# Gray matter correlates of childhood maltreatment: searching for replicability in a multi-cohort brain-wide association study

**DOI:** 10.1101/2024.08.15.608132

**Authors:** Janik Goltermann, Nils Winter, Susanne Meinert, Dominik Grotegerd, Anna Kraus, Kira Flinkenflügel, Luisa Altegoer, Judith Krieger, Elisabeth J. Leehr, Joscha Böhnlein, Linda M. Bonnekoh, Maike Richter, Tim Hahn, Lukas Fisch, Marius Gruber, Marco Hermesdorf, Klaus Berger, Volker Arolt, Katharina Brosch, Frederike Stein, Florian Thomas-Odenthal, Paula Usemann, Lea Teutenberg, Vincent Hammes, Hamidreza Jamalabadi, Nina Alexander, Benjamin Straube, Andreas Jansen, Igor Nenadić, Tilo Kircher, Nils Opel, Udo Dannlowski

## Abstract

Childhood maltreatment effects on cerebral gray matter have been frequently discussed as a neurobiological pathway for depression. However, localizations are highly heterogeneous, and recent reports have questioned the replicability of mental health neuroimaging findings. Here, we investigate the replicability of gray matter correlates of maltreatment (measured retrospectively via the Childhood Trauma Questionnaire) across three large adult cohorts (total N=3225). Pooling cohorts revealed maltreatment-related gray matter reductions, with most extensive effects when not controlling for depression diagnosis (maximum partial R^2^=.022). However, none of these effects significantly replicated across cohorts. Non-replicability was consistent across a variety of maltreatment subtypes and operationalizations, as well as subgroup analyses with and without depression, and stratified by sex. In this work we show that there is little evidence for the replicability of gray matter correlates of childhood maltreatment, when adequately controlling for psychopathology. This underscores the need to focus on replicability research in mental health neuroimaging.

## Introduction

Childhood maltreatment (CM) has been identified to be one of the most important risk factors for the development of affective disorders ^1,2^ and is associated with chronic disease trajectories and poorer treatment outcomes in major depressive disorder (MDD) ^1,3^. Within the past two decades a plethora of neuroimaging studies has repeatedly suggested that experiences of abuse and neglect during childhood are associated with neurobiological alterations in adults^4–8^. Brain regions where these effects have been localized overlap with neural correlates of MDD, giving rise to the notion that neurobiological alterations may mediate the unfavorable effects of CM on clinical trajectories ^9,10^. Thus, studying the neurobiological correlates of CM could give insights into the mechanistic processes of its clinical consequences, potentially informing the optimization of treatments or preventative measures for this population ^11^.

In adults, CM effects on gray matter structure have been observed in an array of regions, with most frequent findings implying the hippocampus, amygdala, dorsolateral prefrontal cortex, insula and anterior cingulate cortex ^8,12–14^. However, the investigation of CM-associated gray matter alterations has yielded considerable heterogeneity in findings regarding localization of effects. Importantly, large-scale consortium studies and meta-analyses do not find these aforementioned regions, but rather report a multitude of other areas to be associated with CM, including the postcentral gyrus and occipital regions ^13^, the median cingulate gyri and supplementary motor area ^15^, the cerebellum and striatum ^16^, as well as the precuneus ^17^.

This heterogeneity could result from the diversity of measurement instruments (e.g., different retrospective self-report scales vs. prospective ratings) and operationalizations of CM (e.g., continuous vs. categorical), as well as different subtypes of maltreatment being studied separately^18–20^. Regarding subtypes of CM there has been considerable debate whether neural correlates could be specific to individual types of experiences. The increasingly influential dimensional model of adversity postulates that different dimensions of CM, such as threat-related and deprivation-related experiences or the unpredictability of one’s environment, underly differential neurobiological processes, consequently leading to differential neural correlates ^21,22^. Evidence for this model in children and adolescents has been accumulated over several studies ^23^. In contrast, other scholars have suggested the relevance of dividing CM experiences even further and have argued that brain alterations are aligned to these experiences in a very specific manner, such as parental verbal abuse impacting gray matter within the auditory cortex or sexual abuse being associated with cortical thinning within the somatosensory cortex ^8^. Another potential source of heterogeneity could stem from varying sample characteristics, differing in diagnoses, the degree of psychopathology and the severity of CM exposure^24^. Furthermore, different statistical approaches have been used. One statistical challenge is a strong phenomenological co-occurrence with mental health problems. Often, psychiatric diagnosis is statistically controlled for which leads to reduced power to detect maltreatment effects because both constructs strongly covary ^1^ and both explain shared variance in neurobiological alterations^9^. On the other side, if not controlling for diagnosis, neurobiological effects due to maltreatment or due to depression are impossible to disentangle. Moreover, evidence suggests that the neural correlates of CM may differ by sex^25–27^, underscoring the importance of carefully considering sex as a factor in these analyses.

The recent debate around questionable replicability in the neuroimaging domain due to underpowered samples and publication bias suggests the possibility of substantial false-positive findings within the previous body of evidence ^28,29^. This notion is supported by evidence for considerable publication bias in meta-analyzed findings of gray matter correlates of CM ^12^. In fact, large-scale neuroimaging consortia, such as the ENIGMA consortium (Frodl et al.^26^; n=3036) or the UK-Biobank (Gheorghe et al.^16^; n=6751), have yielded much smaller effect sizes compared to studies with smaller samples, and have failed to replicate frequently reported associations of CM with the hippocampus or amygdala. However, these consortia still rely exclusively on segmented volumetric brain measures, thus losing spatial resolution, which may account for lower sensitivity to find gray matter alterations, posing a limitation to these findings.

In summary, inconclusive previous findings may result from variability in CM operationalizations, investigated clinical and non-clinical subgroups, varying statistical approaches, insufficient spatial resolution or simply because of false-positive results originating from underpowered studies.

Systematic investigations of the replicability of these neural correlates do not exist to date. To shed light on this heterogeneity and re-evaluate our knowledge about the neurobiological underpinnings of adverse childhood experiences, we investigated the cross-cohort replicability of gray matter correlates of CM. We therefore utilized three large-scale, deeply phenotyped clinical cohort datasets, with a broad range of self-reported maltreatment experiences, in combination with high-resolution voxel-based morphometry (VBM). These rich datasets were assessed and processed in standardized pipelines harmonized across cohorts. We conducted subgroup analyses and probed different operationalizations and subtypes of maltreatment. Additional analyses stratified for sex were run for all models to account for potential sex-specific neural correlates of CM. Replicability was assessed by the spatial overlap of significant findings between our three cohorts, in addition to analyzing all cohorts together in a pooled model. Across all models we tested the hypothesis that CM is associated with lower gray matter volume (GMV).

Here, we show that there is little evidence for the replicability of gray matter correlates of childhood maltreatment, across well-powered adult cohorts, using retrospective self-report measures.

Consistent non-replicability is presented across all maltreatment operationalizations (including CM subtypes and severe forms of CM), subgroup analyses (including individuals with or without MDD) and in additional analyses stratified by sex. The largest evidence for maltreatment-associated gray matter is found when not adequately controlling for confounding MDD diagnosis. In contrast, the association between childhood maltreatment and depression is found across a variety of different clinical characteristics and replicates consistently across cohorts.

## Results

### Associations of childhood maltreatment with demographic and clinical characteristics

CTQ scales were highly interrelated with each other and they showed a pattern of small positive associations with age and small negative associations with education years (Figure 1a). Furthermore, within the MDD participants CTQ scales showed a pattern of weak to moderate associations with previous and current clinical characteristics (Figure 1a). Overall, the relationship between CM reports and demographic and clinical variables was highly similar across the three cohorts, except that age and number of inpatient treatments were not consistently associated with CTQ scales within the BiDirect cohort (Supplementary Figure S1-S3). Participants with a MDD diagnosis reported significantly more severe CM, as compared to HC participants (Figure 1b, Table S5). This was found across all CM subtypes and highly consistent across all cohorts (Figure S2-S5). Largest differences were found for the emotional abuse and neglect subscales (up to r_rank-biserial_=.517).

**Figure 1.**
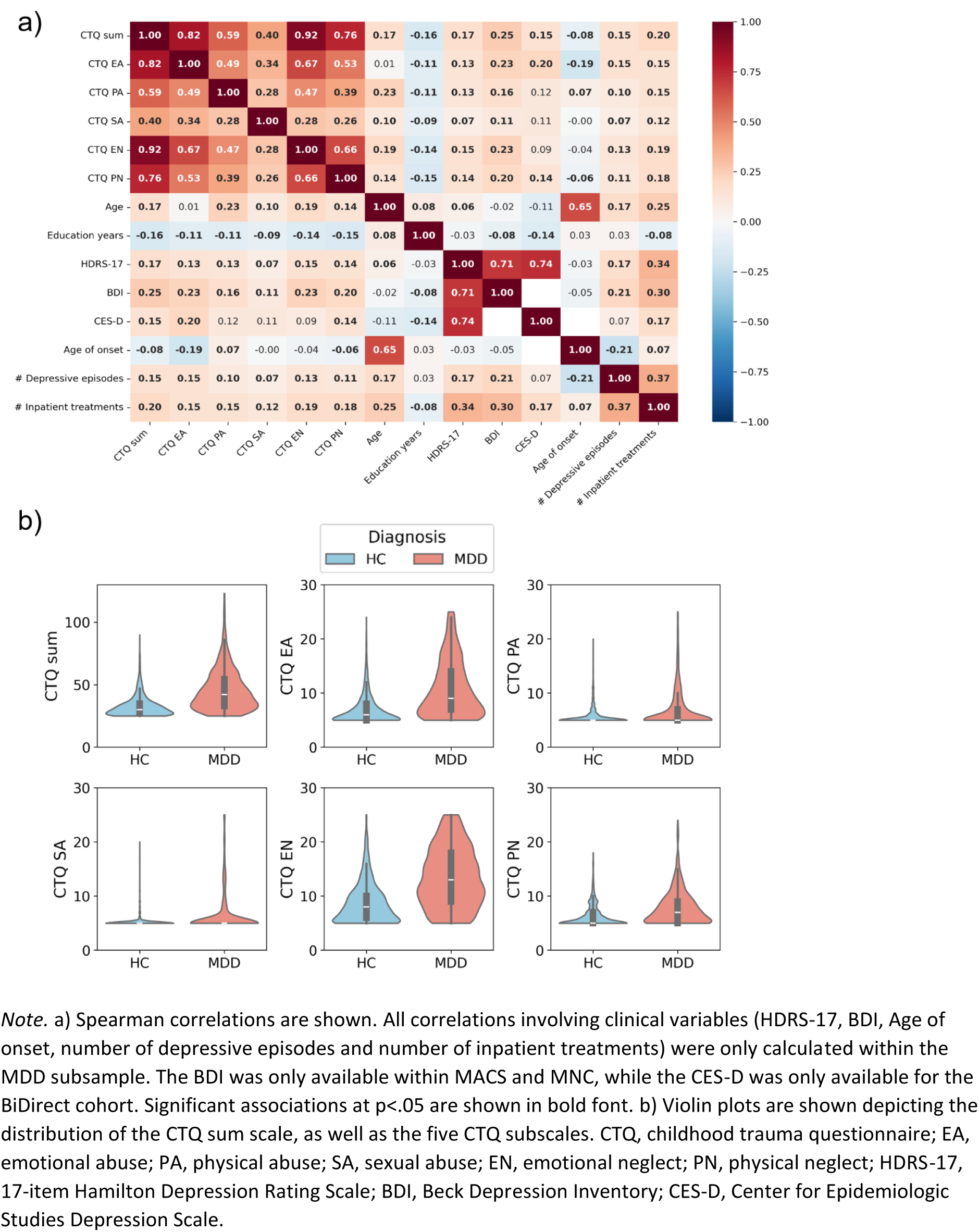
Associations between CTQ scales, demographic variables and clinical variables.

### Gray matter associations in the pooled sample across all cohorts

A total of 15 different statistical models were conducted for all brain-wide analyses. All conducted models are described in Table 1. Results using the full sample from pooling all cohorts together are presented at a conservative significance threshold of p_FWE_<.05, corrected at the voxel-level. Findings from the pooled analyses are shown in Table 2 and Figure 2.

**Figure 2.**
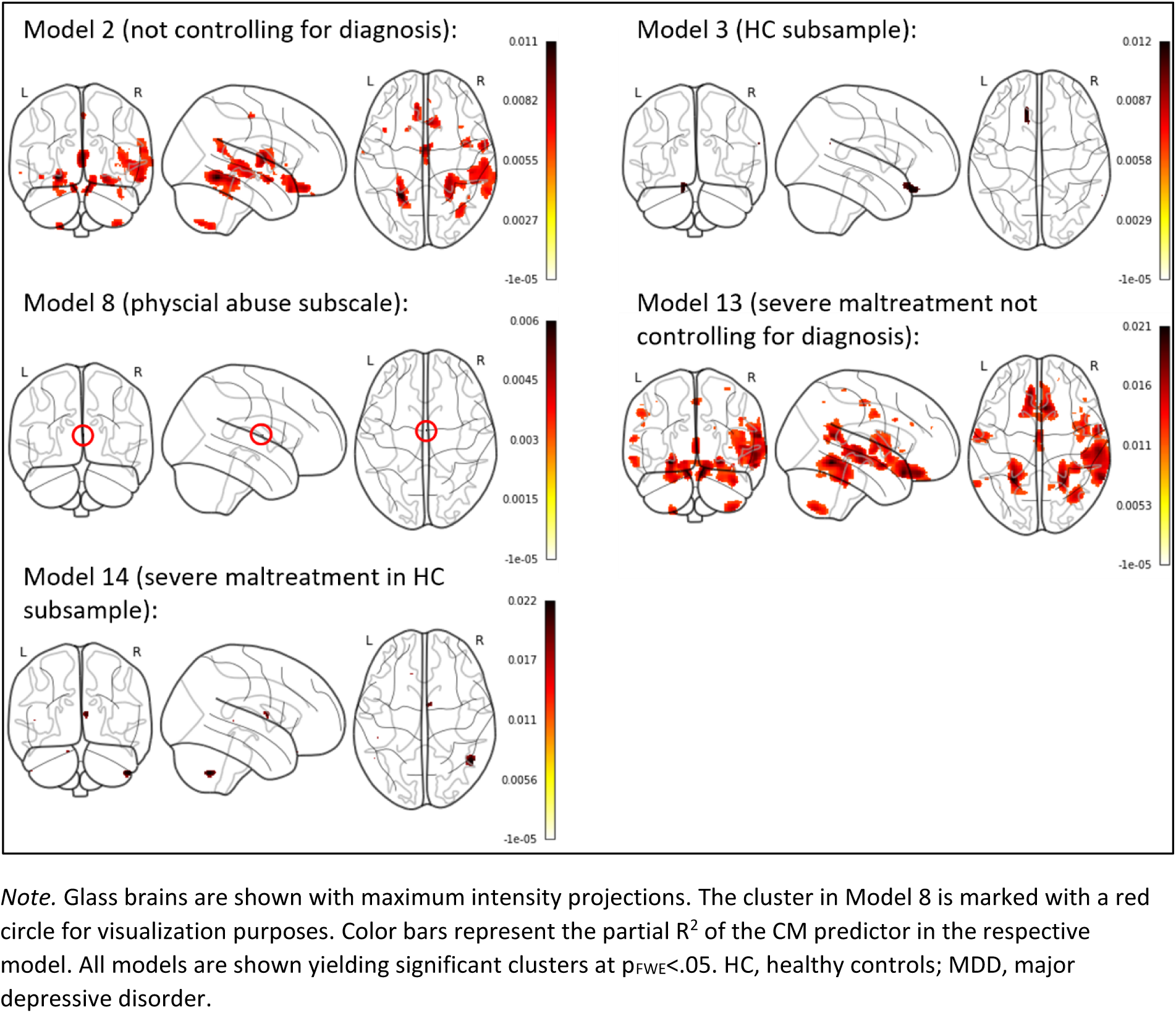
Significant clusters from pooled analysis at p_FWE_<.05.

**Table 1.** Summary of all conducted statistical models and the respective included sample sizes.

| Model | CM<br>operationalization | Controlled for<br>MDD diagnosis | Subsamples | Sample size - n |  |  |  |
| --- | --- | --- | --- | --- | --- | --- | --- |
|  |  |  |  | MACS | MNC | BiDirect | Pooled |
| Model 1 | CTQ sum | yes | HC/MDD | 1752 | 916 | 557 | 3225 |
| Model 2 | CTQ sum | no | HC/MDD | 1752 | 916 | 557 | 3225 |
| Model 3 | CTQ sum | - | HC | 930 | 647 | 321 | 1898 |
| Model 4 | CTQ sum | - | MDD | 822 | 269 | 236 | 1327 |
| Model 5 | Abuse/threat | yes | HC/MDD | 1752 | 916 | 557 | 3225 |
| Model 6 | Neglect/deprivation | yes | HC/MDD | 1752 | 916 | 557 | 3225 |
| Model 7 | EA subscale sum | yes | HC/MDD | 1752 | 916 | 557 | 3225 |
| Model 8 | PA subscale sum | yes | HC/MDD | 1752 | 916 | 557 | 3225 |
| Model 9 | SA subscale sum | yes | HC/MDD | 1752 | 916 | 557 | 3225 |
| Model 10 | EN subscale sum | yes | HC/MDD | 1752 | 916 | 557 | 3225 |
| Model 11 | PN subscale sum | yes | HC/MDD | 1752 | 916 | 557 | 3225 |
| Model 12 | Extreme groups<br>(none/severe) | yes | HC/MDD | None: 644 | None: 369 | None: 213 | None: 1226 |
|  |  |  |  | Severe: 348 | Severe: 145 | Severe: 98 | Severe: 591 |
| Model 13 | Extreme groups<br>(none/severe) | no | HC/MDD | None: 644 | None: 369 | None: 213 | None: 1226 |
|  |  |  |  | Severe: 348 | Severe: 145 | Severe: 98 | Severe: 591 |
| Model 14 | Extreme groups<br>(none/severe) | - | HC | None: 500 | None: 324 | None: 165 | None: 989 |
|  |  |  |  | Severe: 51 | Severe: 41 | Severe: 17 | Severe: 109 |
| Model 15 | Extreme groups<br>(none/severe) | - | MDD | None: 144 | None: 45 | None: 48 | None: 237 |
|  |  |  |  | Severe: 297 | Severe: 104 | Severe: 81 | Severe: 482 |

*Note.* In all models we additionally controlled for age, sex and total intracranial volume. The extreme group comparisons are based on cutoff-based categorizations of severity. For Models with comparison of extreme groups (Models 10-12) sample sizes are given for both the group with ‘none to minimal’ maltreatment (labelled ‘none’) and the group with ‘severe’ maltreatment. CM, childhood maltreatment; CTQ, Childhood Trauma Questionnaire; HC, healthy controls; MDD, major depressive disorder; MACS, Marburg Münster Affective Disorders Cohort Study; MNC, Münster Neuroimaging Cohort; BiDirect, BiDirect study cohort; EA, emotional abuse; PA, physical abuse; SA, sexual abuse; PN, physical neglect; EN, emotional neglect.

**Table 2.**
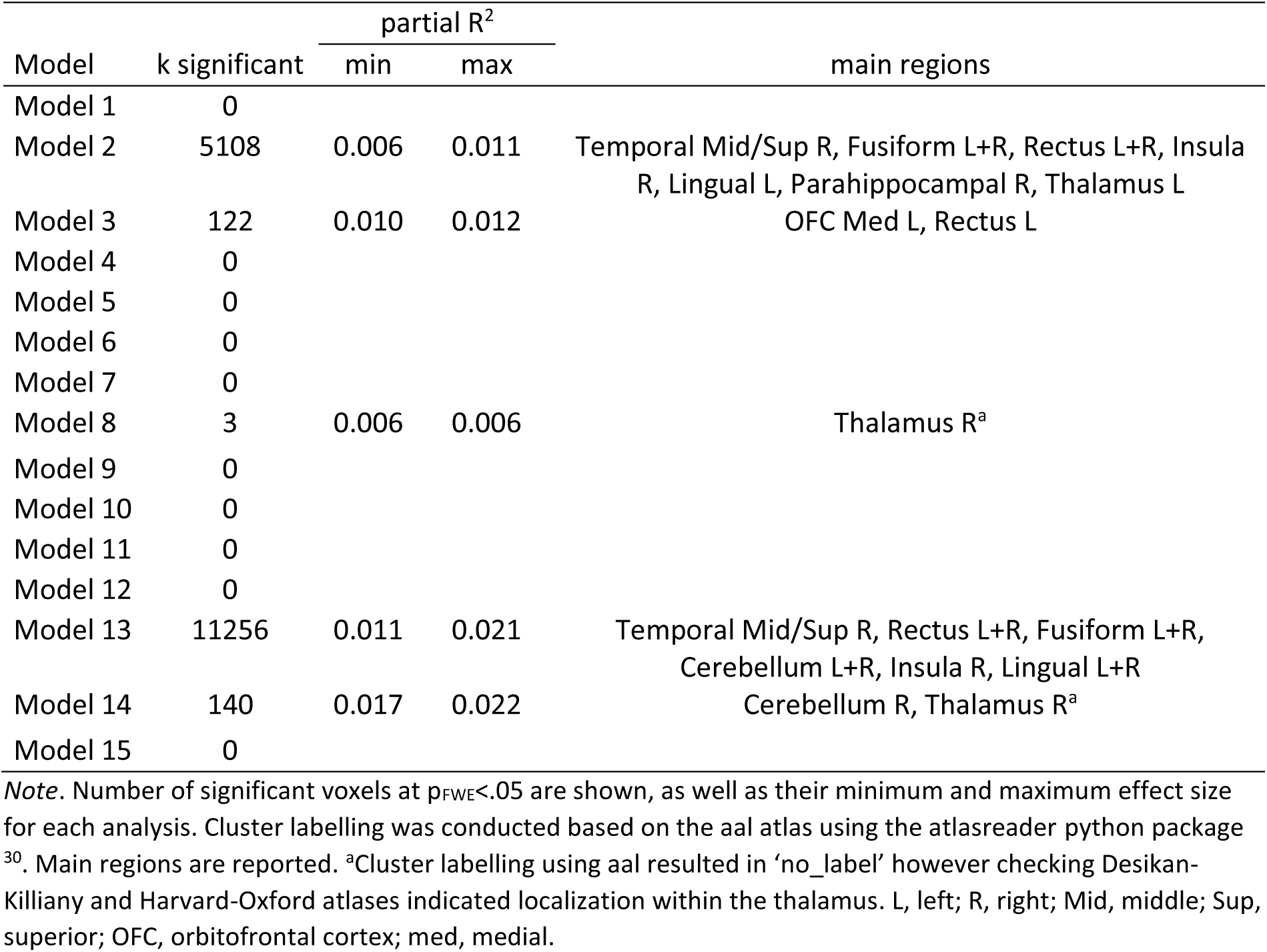
Results summary for pooled cohorts (n=3225) at a significance level of p_FWE_<.05.

When controlling for MDD diagnosis (Model 1) no voxels with a significant CM association were found. Dropping MDD diagnosis as covariate (Model 2) yielded significant widespread clusters (total k=5108), located mainly within superior and middle temporal areas, a bilateral fusiform and lingual complex, the thalamus, as well as in the orbitofrontal cortex and the insula. Subgroup analyses revealed small significant clusters in HC individuals when using CTQ sum as a predictor (Model 3; total k=122) within the medial orbitofrontal cortex, while no clusters survived the FWE-correction within the MDD sample (Model 4). Regarding subtypes of CM, no CTQ subscales were associated with GMV surpassing an FWE-corrected threshold, except a small cluster emerging when using physical abuse as a predictor (Model 8; total k=3 within the thalamus). Similar results were obtained when investigating individuals with ‘severe’ maltreatment: again, the model without controlling for MDD diagnosis yielded widespread reductions in the group with severe maltreatment as compared to the group with ‘none to minimal’ maltreatment in widespread clusters (Model 13; total k=11256). This effect was also found in much smaller localized clusters within HC samples only (Model 14; total k=140). Effect sizes across models when pooling cohorts ranged between partial R^2^=.006 and partial R^2^=.022.

Pooled analyses stratified by sex yielded similar results, however with some additional clusters emerging in female subsamples when investigating severe CM in HC and MDD samples, while controlling for diagnosis (Model 12; total k=1847). Overall, there was a pattern of more models yielding significant effects (and larger clusters) in the female subsamples as compared to male subsamples. Results stratified by sex are shown in Table S6-S7.

### Replicability of gray matter associations across single cohorts

The same 15 models conducted in the pooled cohorts were also fitted in each cohort separately, using liberal uncorrected significance thresholds of p_unc_<.001 and p_unc_<.01. Within single cohorts, both significance thresholds and each statistical model yielded significant voxels in at least one of the three cohorts. In turn, each of the three cohorts produced significant voxels in most of the statistical models. The highest number of significant voxels was observed in model 2 and model 13 – both models where HC and MDD samples were included but diagnosis was not included as a covariate. A detailed summary of cohort-wise results across models, including additional analyses stratified by sex is shown in supplementary Tables S8-S13. Across probed models and across single cohorts the nominally significant voxels were widespread throughout the brain, including the cerebellum, temporal and frontal areas, subcortical areas and somatosensory cortices.

Investigating replicability revealed that there was not one voxel that was congruently significant (i.e., replicable) at a threshold of p_unc_<.001 in all three cohorts. This finding was consistent across all probed statistical models, including HC and MDD subgroup analyses, testing subtypes of CM and comparing groups with severe CM and no CM. Similarly, comparing pairs of cohorts also yielded no voxels that regionally overlapped between any pairwise cohort combinations, for most of the tested models. Only two models yielded marginal pairwise overlap in significance at this threshold: when testing the physical neglect subscale of the CTQ (Model 11) there was a small overlap between the MNC and BiDirect cohorts located within the supramarginal gyrus (overlap k=3; Dice=.002).

Furthermore, there was an overlap of k=2 voxels (Dice=.001) between the MACS and the BiDirect cohort in Model 13 (comparing extreme groups without controlling for diagnosis). This extent of replicability was not significant (p_FDR_>.509), as indicated by permutation-based null-distributions of overlap across cohort-combinations.

When rerunning the replicability analyses using an even more liberal threshold of p_unc_<.01 the observed spatial overlap in significant voxels was increased across models. The only models yielding any overlap across all three cohorts at this threshold was model 2 (CTQ sum; not controlling for MDD diagnosis), with converging significance in k=4 voxels, and model 13 (k=12 voxels; comparing extreme groups of CM, not controlling for MDD diagnosis). Pairwise cohort combinations yielded additional spatial overlap in significance across models, with maximum overlap in model 13 (k=1329, Dice=0.081). All observed overlap of any cohort combination was non-significant. This was consistent across all models (all p_FDR_>.150), as indicated by permutation-based null-distributions of overlap across cohort-combinations.

Replicability results were largely consistent when rerunning all analyses stratified by sex. A summary of the extent of spatial overlap of effects across cohorts, as well as the significance of this replicability is shown in Table 3 and supplementary Tables S14-S18. Significant clusters across significance thresholds, cohorts and statistical models are shown in Figure 3 and supplementary Figures S6-S20, with additional results stratified by sex presented in supplementary Figures S21-S50.

**Figure 3.**
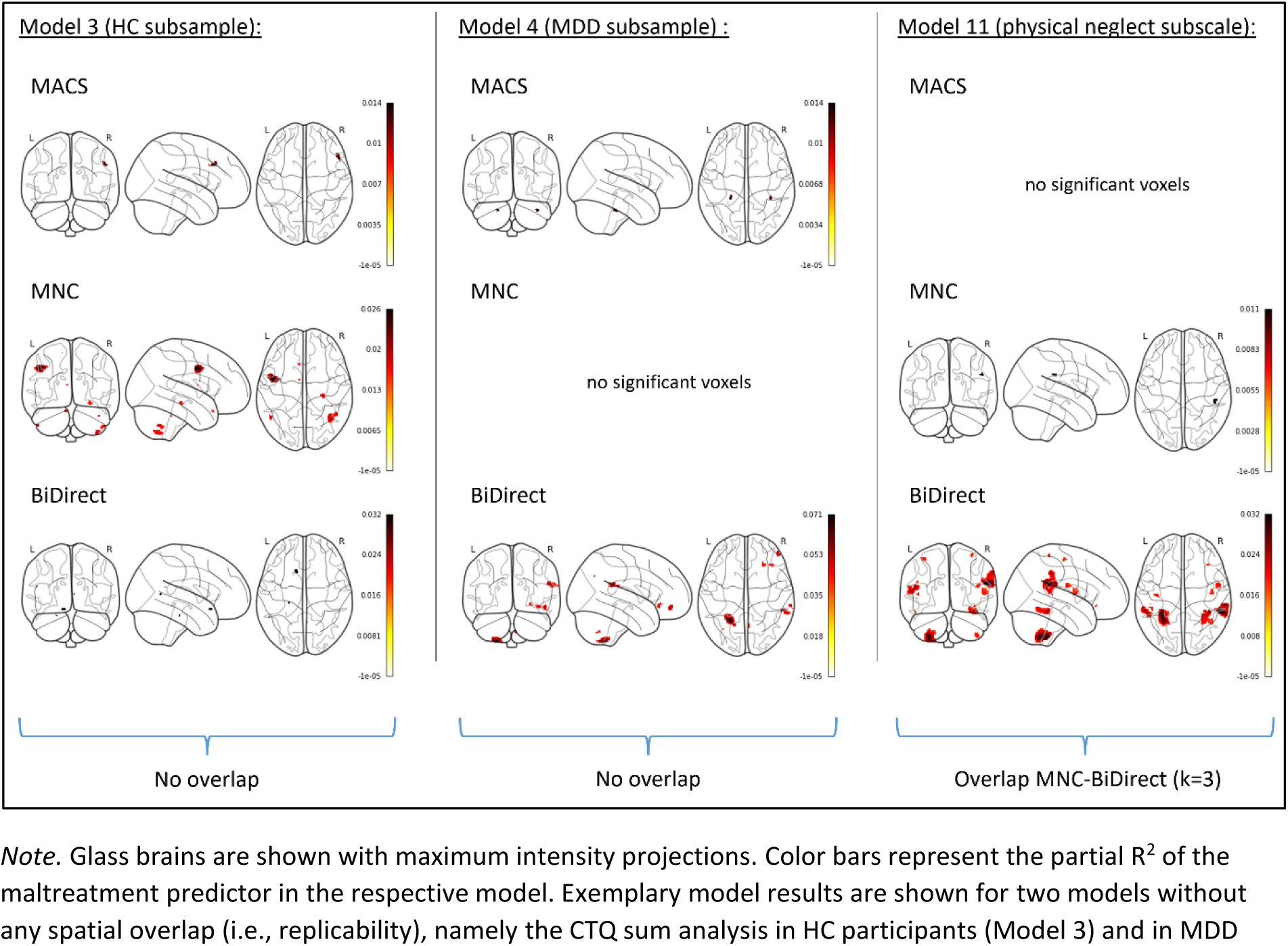
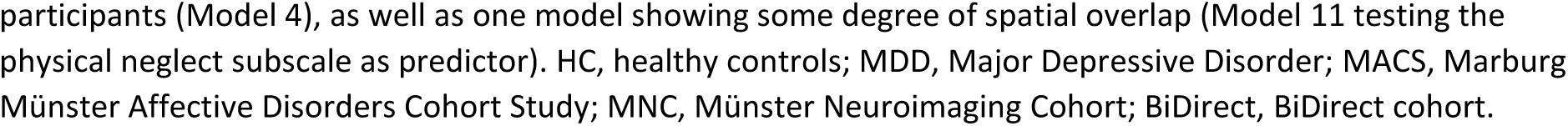
Significant clusters across cohort-wise analyses at p_unc_<.001 for exemplary models (replicability analysis)

**Table 3.** Replicability across cohorts indicated by spatial overlap in significance at a level of p_unc_<.001.

|  | MACS-MNC |  | MACS-BiDirect |  | MNC-BiDirect |  | MACS-MNC-BiDirect |  |
| --- | --- | --- | --- | --- | --- | --- | --- | --- |
|  | k overlap | DICE | k overlap | DICE | k overlap | DICE | k overlap | DICE |
| Model 1 | 0 | 0 | 0 | 0 | 0 | 0 | 0 | - |
| Model 2 | 0 | 0 | 0 | 0 | 0 | 0 | 0 | - |
| Model 3 | 0 | 0 | 0 | 0 | 0 | 0 | 0 | - |
| Model 4 | 0 | 0 | 0 | 0 | 0 | 0 | 0 | - |
| Model 5 | 0 | 0 | 0 | 0 | 0 | 0 | 0 | - |
| Model 6 | 0 | 0 | 0 | 0 | 0 | 0 | 0 | - |
| Model 7 | 0 | 0 | 0 | 0 | 0 | 0 | 0 | - |
| Model 8 | 0 | 0 | 0 | 0 | 0 | 0 | 0 | - |
| Model 9 | 0 | 0 | 0 | 0 | 0 | 0 | 0 | - |
| Model 10 | 0 | 0 | 0 | 0 | 0 | 0 | 0 | - |
| Model 11 | 0 | 0 | 0 | 0 | 3<sup>a</sup> | 0.002 | 0 | - |
| Model 12 | 0 | 0 | 0 | 0 | 0 | 0 | 0 | - |
| Model 13 | 0 | 0 | 2<sup>b</sup> | 0.001 | 0 | 0 | 0 | - |
| Model 14 | 0 | 0 | 0 | 0 | 0 | 0 | 0 | - |
| Model 15 | 0 | 0 | 0 | 0 | 0 | 0 | 0 | - |

*Note.* The overlap in significant voxels is presented across all statistical models and all cohort combinations. The DICE score is presented for pairwise combinations with any voxels overlapping. MACS, Marburg Münster Affective Disorders Cohort Study; MNC, Münster Neuroimaging Cohort; BiDirect, BiDirect study cohort.
<sup>a</sup>Corresponding significance of overlap: $p = .011$ , $p_{FDR} = .509$ . <sup>b</sup>Corresponding significance of overlap: $p = .017$ , $p_{FDR} = .509$ .

## Discussion

Findings implicating long-term effects of CM on the structural morphology of the brain have been frequently published over the last decades and are central to a neurobiological model of how environmental risk is conveyed to psychopathology. However, in an unprecedented replication effort we present evidence that localized brain-wide associations between gray matter structure and various operationalizations of CM are essentially non-replicable. This lack of replicability was consistently shown for a wide variety of common statistical approaches, across non-clinical and clinical, as well as sex-stratified subgroups and across a variety of operationalizations of CM. Central limitations arise due to the concrete assessment method of CM utilized in this study (retrospectively via the CTQ) and due to low demographic (particularly ethnic) variability of the included samples. The extensive non-replicability of CM-related gray matter effects were contrasted by highly replicable negative associations of CM with MDD diagnosis and with various measures of current and previous depression severity.

In our pooled analysis we found small significant effects within HC subsamples and when investigating physical abuse. Furthermore, in the pooled analysis we identified large and widespread clusters when not controlling for confounding MDD diagnosis. Notably, the localized clusters seemingly associated with CM in this latter analysis largely overlap with clusters that were identified to be associated with a lifetime MDD diagnosis in a systematic case-control study using the same cohorts ^31^.

When using liberal uncorrected thresholds, we found a vast array of regions seemingly associated with CM across the different cohorts and statistical models. In isolation, each of these results could easily have been the basis of a publication just like several smaller existing studies in the field, including previous publications from our own group ^4,9^. Importantly, these widespread significant effects for each single cohort should not be interpreted as solid evidence for effects due to the liberal significance thresholds and resulting massively inflated alpha error (i.e., false-positives). In our most liberal analyses, some extent of replicability was observed, particularly when investigating HC and MDD samples together while not controlling for MDD diagnosis. However, the identified overlap was small even for pairwise combinations of cohorts. Furthermore, permutation-based significance testing of this descriptive overlap indicated that it was not higher than expected by chance. Overall, our findings suggest that gray matter reductions associated with CM are non-replicable. Similar results were obtained when rerunning all analyses stratified by sex. Even highly cited previous reports of brain structural correlates of CM, such as associations with lower GMV within the hippocampus ^4,9^ could not be confirmed.

Null findings are always difficult to interpret due to a multitude of potential reasons for failing to detect an effect. Potential reasons for false-negatives can stem from the specific measurement and operationalization of the predictor or dependent variable, the specific statistical approach (e.g., inclusion of covariates), insufficient statistical power, the sample selection and, regarding replicability, differences between specific cohort characteristics. In the following we will discuss each of these potential sources of effect variability.

Despite of its common use, the CTQ has been criticized for neglecting the timing of maltreatment ^32^ and showing low agreement with prospective CM measures ^33,34^, the latter likely due to memory and reporting biases ^35^. Although depressive states are thought to bias childhood reports, we found CTQ scores to be highly stable over two years within the MACS and MNC cohorts, with no systematic association with changes in depressive symptoms ^36^. Regardless, retrospective CM measures generally show stronger associations with later psychopathology than prospective measures, potentially measure different entities ^37^. Timing of exposure may critically moderate CM’s effects on clinical ^38–40^ and neurobiological endpoints ^41–43^. Thus, it is unclear whether our findings, based on the retrospective CTQ, generalize to prospectively assessed CM measures or those incorporating timing information. While the general critique of the CTQ may be valid and posits an important limitation to the current findings, it is notable that the CTQ is also the instrument which has been used in most of the referenced studies, which have reported GMV associations with CM ^4,9,44–47^, ensuring methodological comparability with our study. Some of the studies reporting on significant effects use alternative assessment instruments to measure CM, such as interview measures ^44^, or CTQ data from adolescent or young adult samples, potentially less affected by memory biases ^44,45,47^. Furthermore, our analyses are cross-sectional, and while longitudinal studies are rare, some have shown how CM may affect brain development and links to psychopathology over time ^45,48–50^. Future studies should expand our current findings and investigate the replicability of CM neural correlates using alternative instruments, designs and samples.

On the side of the dependent variable, we used state-of-the-art voxel-wise GMV assessments. Meta-research on neuroimaging replicability suggests that the researchers’ degrees of freedom regarding scanning parameters, preprocessing and quality control pipelines contribute to low reproducibility and replicability ^51,52^. While such differences could influence findings, our procedures were closely harmonized across cohorts, making this an unlikely explanation for low replicability in our study. Further, it remains unclear whether our null findings generalize to other imaging modalities, such as functional or structural connectivity, or alternative measures of gray matter structure (e.g., cortical surface or thickness), warranting further investigation.

Similarly, several decisions are required regarding the operationalization of predictors and statistical modeling. In previous studies, CM (even when based on the CTQ) was operationalized in different ways (total sum score, subscale scores, different cutoff-based categories), and confounding psychopathology was differently addressed. To enable more robust interpretations, we employed a comprehensive approach that included a range of common statistical models. While our list of models is not exhaustive and our conclusions are limited to these specific approaches, the consistent finding of poor replicability across all tested models is striking.

Insufficient statistical power may partly account for low replicability, especially for some subgroup analyses. Although our study represents the largest replicability investigation of brain alterations associated with CM to date, it may still lack power to detect small effects typical in biological psychology and psychiatry, which rarely exceed 2.5% explained variance ^28,53^. This limitation is particularly relevant for some subgroup analyses (smallest subsample size n=129). However, our attained sample size and statistical power are well within the realm of previous meta-analyses ^12,13^ and large-scale consortia analyses ^17,54^.

Sample selection is a key source of variability. While our cohorts encompass a broad range of maltreatment severity and clinical characteristics, they are relatively homogeneous demographically, consisting of German individuals of Western European ancestry with relatively high education levels. This homogeneity enhances conditions for replicability but limits the generalizability of our null findings to other populations. Differences between cohorts may also contribute to low replicability. For instance, the BiDirect cohort was considerably older, while the MNC cohort included only acutely depressed inpatient MDD patients, compared to a mix of outpatient and remitted individuals in the other cohorts. Differences in current depression severity and illness history were also observed across MDD samples. However, the extent of non-replicability across all pairwise cohort comparisons suggests that cohort-specific differences alone are unlikely to fully explain our null findings.

All these aspects pose potential sources of effect variability and could account for false-negative findings. However, previous studies reporting CM associations were highly comparable regarding utilized methodology. It should be noted that it is possible that conventional conceptualizations of clearly localizable gray matter reductions due to CM on a group-level could be too simplistic.

Recently, machine learning and normative modelling approaches have been increasingly promoted following the notion that the concrete shape of neurobiological consequences in the brain may be highly individual ^55,56^.

The absence of evidence cannot directly be interpreted as evidence for the absence of a phenomenon ^57^. However, the extent of non-replicability of gray matter correlates of CM still appears disconcerting. Notably, this is in contrast to the replicability of GMV reductions linked to lifetime MDD, observed using a similar approach ^31^. Replicability is a fundamental principle of the scientific process and essential for accumulating scientific knowledge ^58^. However, non-replicability of published findings is a growing concern across disciplines, such as cell biology ^59^, genetics ^60,61^, oncology ^62^, epidemiology ^63^ and psychology ^64^. Factors contributing to this *replication crisis* include publication pressure, bias toward positive results, and analytic flexibility ^65,66^. Neuroimaging, with its high analytic flexibility, numerous tests, and small, underpowered samples, is particularly susceptible to overestimated effect sizes and non-replicability ^67,68^. Recent research shows that thousands of participants may be needed for robust, replicable brain-wide associations due to small true effect sizes ^28^, which has sparked ongoing debates in the field^69–74^. Our study supports this view, suggesting that low replicability may be a broader issue, not limited to this research question alone. Notably, no consensus exists on how to define “successful” replicability in voxel-based neuroimaging. We contribute to this by formalizing and testing cross-sample replicability of voxel-based analyses.

Concerns about low replicability have led to the development of open science policies, such as preregistration of hypotheses and analysis plans, as well as comprehensive disclosure of analysis code and results ^75^, to achieve transparency and reduce biases. Accordingly, scholars have increasingly advocated for these practices to enhance replicability in neuroimaging research ^76^. However, replications and open science practices remain very rare in neuroimaging ^77^. Additionally, approaches like cross-validation, which assess the generalizability of statistical findings to independent data, can help identify overestimated effect sizes and non-replicability in smaller samples ^78^.

Our findings underline the importance of taking a step back and shifting the focus towards increasing and investigating the replicability and generalizability of presumably established research findings.

Various open science practices are available for this: 1) preregistrations of hypotheses and analyses, 2) transparent sharing of analysis code and methods, and the publication of comprehensive (i.e., non-thresholded) results ^79^, as well as 3) the mere execution of direct and conceptual replication studies, and 4) the publication of null findings. Open science practices should be routinely adopted in neuroimaging research on mental disorders to increase replicability and thus maximize the potential for clinical translation.

## Methods

### Participants

Samples from three large-scale independent cohorts were included in the present analysis: the Marburg Münster Affective Disorders Cohort Study (MACS), the Münster Neuroimaging cohort (MNC) and the BiDirect cohort. All three cohorts include adults (age 18-65 years) with and without mental disorders. Recruitment was restricted to individuals proficient in the German language and with western European ancestry (as the cohorts were originally conceptualized for genetic analyses). For the current analyses we included healthy control (HC) individuals, as well as individuals with a lifetime MDD diagnosis. In total, a sum of n=3225 participants were included (HC: n=1898; MDD: 1327). Identical exclusion criteria were applied for all three independent cohorts: 1) duplicate cases resulting from individuals that were included in more than one of the utilized cohorts, 2) presence of a lifetime bipolar disorder, psychosis spectrum disorder or substance dependencies (other psychiatric comorbidities were permitted), 3) severe head trauma or severe/chronic somatic illness (e.g., Parkinson’s disease, multiple sclerosis, stroke, myocardial infarction), 4) missing MRI data and image artefacts diagnosed during quality control, 5) missing data in the CTQ.

For details on the methods and general inclusion criteria of the study samples we refer to previous publications (MACS: Kircher et al.^80^; Vogelbacher et al.^81^; MNC: Dannlowski et al.^82^; Opel et al.^6^; BiDirect: Teismann et al.^83^) and to the supplements. The final samples included in the current analyses comprised n=1752 participants from MACS (HC: n=930; MDD: n=822), n=916 participants from MNC (HC: n=647; MDD: n=269), and n=557 participants from BiDirect (HC: n=321; MDD: n=236). Detailed sample characteristics of the three cohorts, including demographics, reports of CM and clinical characteristics, are described in Table S1 and Table S2. Differences between cohorts in sample characteristics are shown in Table S3, while differences in clinical characteristics between the MDD subsamples of the cohorts are shown in Table S4. Age distributions across cohorts and diagnosis groups are shown in Figure S1.

Of note, findings regarding gray matter correlates of CM have been previously published using MNC data at earlier stages of data assessments. However, these analyses only included a fraction of the data available for the current analysis (the largest sample including n=170 subjects).^4,9^

The study was approved by the local Institutional Review Board of the medical faculties of the University of Marburg and the University of Münster and written informed consent was obtained before participation. Patients received financial compensation for their participation.

### Assessment of childhood maltreatment and clinical characterization

CM was assessed using the German version of the Childhood Trauma Questionnaire (CTQ) ^84,85^. The CTQ is a 25-item retrospective self-report questionnaire capturing five different subtypes of CM, namely emotional abuse, physical abuse, sexual abuse, emotional neglect, and physical neglect. Each of these subtypes can be scored separately using a sum score of the five corresponding items, while the sum of these subscale scores amounts to the total CTQ score (expressing the total severity or load of experienced maltreatment). In addition, a categorical scoring of the CTQ has been introduced based on validated subscale cutoff values, dividing scores in each subscale into different severity categories from ‘none to minimal’ to ‘severe’ ^86^. As described in detail below, here we utilize the total and subscale sum scores, as well as categorical cutoffs for group comparisons. The CTQ has been used in several hundreds of studies across different nationalities and validated on clinical and non-clinical populations ^87^. It has been extensively tested for its psychometric properties in several languages and geographic contexts ^85,87–94^.

Lifetime clinical diagnosis was assessed using structured clinical interviews by trained study personnel in each cohort. Within MACS and MNC the German version of the Structured Clinical Interview for the DSM-IV (SCID)^95^ was used. Within the BiDirect cohort the Mini International Neuropsychiatric Interview (MINI)^96^ was used, also based on the DSM-IV criteria. Further clinical characterization was done using a variety of standardized clinical interviews, rating scales and self-report questionnaires, capturing information on current remission status, current depression severity and previous course of disease (see supplements).

### Structural image acquisition and processing

T1-weighted high-resolution anatomical brain images were acquired using 3T MRI scanner using highly harmonized scanning protocols across all three cohorts. For the MACS sample two different MRI scanners were used at the recruitment sites in Marburg (Tim Trio, Siemens, Erlangen, Germany; combined with a 12-channel head matrix Rx-coil) and Münster (Prisma, Siemens, Erlangen, Germany; combined with a 20-channel head matrix Rx-coil). MNC and BiDirect samples were both scanned using a Gyroscan Intera scanner with Achieva update (both by Philips Medical Systems, Best, The Netherlands).

Image preprocessing was conducted using the CAT12-toolbox^97^ (https://neuro-jena.github.io/cat/) using default parameters equally for all cohorts. Briefly, images were bias-corrected, tissue classified, and normalized to MNI-space using linear (12-parameter affine) and non-linear transformations, within a unified model including high-dimensional geodesic shooting normalization ^98^. The modulated gray matter images were smoothed with a Gaussian kernel of 8 mm FWHM. Absolute threshold masking with a threshold value of 0.1 was used for all second level analyses as recommended for VBM analyses (http://www.neuro.uni-jena.de/cat/). Image quality was assessed by visual inspection as well as by using the check for homogeneity function implemented in the CAT12 toolbox. Image acquisition and processing for all study samples were extensively described elsewhere ^48,81,83^.

Image harmonization was conducted using the neuroCombat toolbox in python ^99^ with default parameters to control for differences in scanner hardware and corresponding effects on brain images. This procedure allows the specification of ‘biological covariates’ that are excluded from harmonization in order to preserve desired variance of potentially confounding variables. We defined CTQ sum, age, sex, total intracranial volume (TIV) and MDD diagnosis as such covariates. The harmonization process was conducted across six different scanner groups: MACS Münster scanner, MACS Marburg scanner before and after body coil change, MNC scanner before and after gradient coil change and the BiDirect scanner setting.

### Statistical analysis

Associations between CM reports, demographic and clinical variables were investigated using spearman correlations (due to highly non-normal distributions in the CTQ scales) and Man-Whitney-U tests. For the latter, rank-biserial correlations were calculated as a measure of effect size.

Brain-wide associations between CM and voxel-wise GMV were tested using general linear models in a mass-univariate VBM approach. The available cohorts were investigated in two different steps: In a first step we pooled all cohorts together in order to harvest the maximum sample size and thus the maximum available statistical power. For this pooled analysis we used a voxel-wise family-wise error (FWE)-corrected significance threshold of p_FWE_<.05.

In a second step we investigated the cross-cohort replicability by analyzing each of the cohorts separately using two liberal uncorrected significance thresholds of p_unc_<.001 and p_unc_<.01. Here, we examined the spatial convergence (i.e., overlap) of significant voxels across the cohorts as conjunctive criteria (convergence either across any subset of two cohorts or across all three cohorts). Note that these liberal significance thresholds should not be used by themselves for statistical inference due to a massively inflated alpha error from the mass-univariate testing. However, we defined these liberal thresholds as minimum thresholds for effects to be recognized as replicable.

Importantly, the probability of finding the same voxel in two or even three cohorts constitutes a higher threshold than a voxel becoming significant just in a single cohort. In numbers, a threshold of p<.001 exceeded in each of the three single cohorts, results in an effective false-positive rate of p<.001³ = .000000001. No extent threshold for minimum cluster size was used in any analysis. In order to test the significance of the replicability (i.e., overlap in effects) we applied permutation testing, permuting the respective predictor label for each cohort k=1000 times subsequently obtaining a null distribution for the overlap analysis (how much overlap between cohorts can be expected by chance). Obtained p-values were FDR-corrected using the Benjamini-Hochberg procedure^100^ across 15 models for which overlap was investigated across four different cohort-combinations (resulting in correction of sets of 60 tests).

The following statistical models were probed for 1) pooled analyses and 2) cross-cohort replicability analyses, to delineate the conditions under which VBM associations may become evident (an overview of all statistical models is presented in Table 1). In all models age, sex and TIV were included as covariates. Additionally, lifetime MDD diagnosis was included as a control variable in all models unless stated otherwise:

– In **Model 1** we used CTQ sum as the main predictor of interest, while additionally controlling for lifetime MDD diagnosis, as this variable is highly confounded with CM. Thus, in this model we tested the effect of CM on GMV beyond any effect of diagnosis.
– The model described above may not be sufficiently sensitive to detect CM effects due to substantial shared variance with the MDD diagnosis effect being partialized out. Therefore, we further tested a second model (**Model 2**) removing MDD diagnosis as a covariate. This allowed us to obtain a liberal estimate for the association between CM and gray matter, which however is not clearly separable from any MDD diagnosis effects.
– To investigate CM effects independently from a confounding MDD diagnosis effect, we further conducted subgroup analyses within HC (**Model 3**) and MDD (**Model 4**) samples separately.
– Based on the dimensional model of adversity ^21,22^ we probed associations between CM and GMV specifically for the abuse subscales (**Model 5**) and the neglect subscales (**Model 6**) of the CTQ, to differentiate between threat– and deprivation-related experiences.
– In order to delineate effects of specific subtypes of CM in further detail, we tested a series of models with each of the five CTQ subscale sum scores as predictors respectively (**Models 7-11**).
– Lastly, we investigated the effects of severe forms of CM. This was done by identifying participants exceeding the subscale cutoff score for severe CM in any CTQ subscale, as defined and validated by Bernstein and colleagues.^101^ This group of individuals with severe CM is contrasted with a control group that does not exceed any subscale cutoff (only CM reported within the range of “none to minimal”). With this we accounted for the notion that CM associations with VBM may become evident particularly in individuals with severe experiences of CM. In our sample n=591 individuals (HC: n=109; MDD: n=482) fulfilled the criteria for severe maltreatment in at least one of the CTQ subscales, while n=1226 individuals (HC: n=989; MDD: n=237) fell into the category “none to minimal” maltreatment experiences. The extreme group comparisons were done in the full sample (HC and MDD), in one model controlling for MDD diagnosis (**Model 12**) and in another model dropping MDD diagnosis as a covariate (**Model 13**). The distribution of severe CM across diagnostic groups was significantly uneven (Chi^2^=645.71; p<.001, OR=18.45, 95%-CI: [14.348, 23.732). This strongly unequal distribution led us to conduct severity group analyses additionally in HC (**Model 14**) and MDD (**Model 15**) subgroups separately.

To account for potential sex-specific effects all analyses were additionally rerun stratified by female and male sex, as self-reported by the participants. One-sided negative contrasts were tested in all models (i.e., CM associated with lower gray matter volume) due to poor evidence for potential positive associations between CM and VBM ^12,13,15,23^. Partial R² was calculated based on t-maps and reported as an effect size for the partialized percentage of explained variance of the respective CM predictor for each analysis. Analyses were conducted using python (version 3.9.12). Analyses were not preregistered.

## Data availability

Comprehensive non-thresholded statistical estimates are made openly available via the OSF (https://osf.io/j8d9r/?view_only=9edf436ab18f4e8db9ef4c71c4ac356c). Individual raw data is not published due to current EU data protection regulations and the sensitive nature of clinical MRI data but can be made available in form of summary statistics or anonymized aggregation of voxel-wise data upon reasonable request to the corresponding author.

## Code availability

The code used for analysis is publicly available in an Open Science Framework (OSF) repository (https://osf.io/j8d9r/?view_only=9edf436ab18f4e8db9ef4c71c4ac356c), to foster transparency and reproducibility of our analyses ^29^.

## Supporting information

Supplemental methods and results

## Funding and Disclosures

This project was in part supported by the SFB/TRR 393 funded by the Deutsche Forschungsgemeinschaft (DFG, German Research Foundation; project-ID 521379614). The MACS and MNC studies are also funded by the DFG (grant FOR2107 DA1151/5-1 and DA1151/5-2 to UD; SFB-TRR58, Projects C09 and Z02 to UD), the Interdisciplinary Center for Clinical Research (IZKF) of the medical faculty of Munster (grant Dan3/012/17 to UD), IMF Munster RE 22 17 07 to JR and the Deanery of the Medical Faculty of the University of Munster. SM received funding by the “Innovative Medizinische Forschung” (IMF) of the medical faculty of Münster (ME122205) and the Else-Kröner Fresenius Stiftung (EKFS; 2023_EKEA.153). JG also received funding by the IMF (GO122301). TH was supported by the German Research Foundation (DFG grants HA7070/2-2, HA7070/3, HA7070/4). The BiDirect study is funded by German Federal Ministry of Education and Research Grant Nos. 01ER0816, 01ER1506, and 01ER1205. Biomedical financial interests or potential conflicts of interest: TK received unrestricted educational grants from Servier, Janssen, Recordati, Aristo, Otsuka, neuraxpharm. This cooperation has no relevance to the work that is covered in the manuscript.

## Notes

https://osf.io/j8d9r/?view_only=9edf436ab18f4e8db9ef4c71c4ac356c

### Summary of Updates

The Combat image harmonization was revised, now including CTQ sum as an additional 'biological covariate'. Furthermore, we added further analyses, now also including derivates of the 'abuse' and 'neglect' subscales of the CTQ, as well as all analyses rerun stratified by sex. The rest of the manuscript was streamlined.

