## Supplemental methods and results for "Gray matter correlates of childhood maltreatment: searching for replicability in a multi-cohort brain-wide association study"

**Supplements**


**Table S1:** *Demographic and clinical characteristics across all three cohorts*

**Table S2:** *Summary statistics for reported experiences of childhood maltreatment in the CTQ across cohorts*

**Table S3:** *Pairwise cohort comparison in sample characteristics including HC and MDD participants*

**Table S4:** *Pairwise cohort comparison of clinical characteristics of the MDD subgroups*

**Table S5:** *Differences in CTQ scales across HC and MDD groups in pooled sample and cohort-wise*

**Table S6:** *Results summary for pooled cohorts (n=3225) at a significance level of* $p_{FWE}<.05$ *–* ***sex-stratified for female subsample***

**Table S7:** *Results summary for pooled cohorts (n=3225) at a significance level of* $p_{FWE}<.05$ *–* ***sex-stratified for male subsample***

**Table S8:** *Results summary of single cohorts at a cohort-wise significance level of* $\boldsymbol{p_{unc}<.001}$

**Table S9:** *Results summary of single cohorts at a cohort-wise significance level of* $\boldsymbol{p_{unc}<.001}$ ***– sex-stratified for female subsample***

**Table S10:** *Results summary of single cohorts at a cohort-wise significance level of* $\boldsymbol{p_{unc}<.001}$ ***– sex-stratified for male subsample***

**Table S11:** *Results summary of single cohorts at a cohort-wise significance level of* $\boldsymbol{p_{unc}<.01}$

**Table S12:** *Results summary of single cohorts at a cohort-wise significance level of* $\boldsymbol{p_{unc}<.01}$ ***– sex-stratified for female subsample***

**Table S13:** *Results summary of single cohorts at a cohort-wise significance level of* $\boldsymbol{p_{unc}<.01}$ ***– sex-stratified for male subsample***

**Table S14:** *Replicability across cohorts indicated by spatial overlap in significance at a cohort-wise level of* $\boldsymbol{p_{unc}<.001}$ ***– sex-stratified for female subsample***

**Table S15:** *Replicability across cohorts indicated by spatial overlap in significance at a cohort-wise level of* $\boldsymbol{p_{unc}<.001}$ ***– sex-stratified for male subsample***

**Table S16:** *Replicability across cohorts indicated by spatial overlap in significance at a cohort-wise level of* $\boldsymbol{p_{unc}<.01}$

**Table S17:** *Replicability across cohorts indicated by spatial overlap in significance at a cohort-wise level of* $\boldsymbol{p_{unc}<.01}$ ***– sex-stratified for female subsample***

**Table S18:** *Replicability across cohorts indicated by spatial overlap in significance at a level of* $\boldsymbol{p_{unc}<.01}$ ***– sex-stratified for male subsample***

**Figure S1:** *Age distribution across cohorts and diagnosis groups*

**Figure S2:** *Association between CTQ scales, demographic variables and clinical variables within the* ***MACS*** *cohort*

**Figure S3:** *Association between CTQ scales, demographic variables and clinical variables within the* ***MNC*** *cohort*

**Figure S4:** *Association between CTQ scales, demographic variables and clinical variables within the **BiDirect** cohort*

**Figure S5:** *Distributions of CTQ scales across HC and MDD groups stratified by cohorts*

**Figure S6:** *Significant clusters across cohort-wise analyses (replicability analysis) for **model 1** – CTQ sum in HC and MDD samples controlling for MDD diagnosis*

**Figure S7:** *Significant clusters across cohort-wise analyses (replicability analysis) for **model 2** – CTQ sum as a predictor in HC and MDD samples without controlling for MDD diagnosis*

**Figure S8:** *Significant clusters across cohort-wise analyses (replicability analysis) for **model 3** – CTQ sum as a predictor in HC subsamples*

**Figure S9:** *Significant clusters across cohort-wise analyses (replicability analysis) for **model 4** – CTQ sum as a predictor in MDD subsamples*

**Figure S10:** *Significant clusters across cohort-wise analyses (replicability analysis) for **model 5** – CTQ abuse subscales as a predictor in HC and MDD samples controlling for MDD diagnosis*

**Figure S11:** *Significant clusters across cohort-wise analyses (replicability analysis) for **model 6** – CTQ neglect subscales as a predictor in HC and MDD samples controlling for MDD diagnosis*

**Figure S12:** *Significant clusters across cohort-wise analyses (replicability analysis) for **model 7** – CTQ subscale emotional abuse as a predictor in HC and MDD samples controlling for MDD diagnosis*

**Figure S13:** *Significant clusters across cohort-wise analyses (replicability analysis) for **model 8** – CTQ subscale physical abuse as a predictor in HC and MDD samples controlling for MDD diagnosis*

**Figure S14:** *Significant clusters across cohort-wise analyses (replicability analysis) for **model 9** – CTQ subscale sexual abuse as a predictor in HC and MDD samples controlling for MDD diagnosis*

**Figure S15:** *Significant clusters across cohort-wise analyses (replicability analysis) for **model 10** – CTQ subscale emotional neglect as a predictor in HC and MDD samples controlling for MDD diagnosis*

**Figure S16:** *Significant clusters across cohort-wise analyses (replicability analysis) for **model 11** – CTQ subscale physical neglect as a predictor in HC and MDD samples controlling for MDD diagnosis*

**Figure S17:** *Significant clusters across cohort-wise analyses (replicability analysis) for **model 12** – CTQ extreme severity compared to none in HC and MDD samples controlling for MDD diagnosis*

**Figure S18:** *Significant clusters across cohort-wise analyses (replicability analysis) for **model 13** – CTQ extreme severity compared to none in HC and MDD samples without controlling for MDD diagnosis*

**Figure S19:** *Significant clusters across cohort-wise analyses (replicability analysis) for **model 14** – CTQ extreme severity compared to none in HC subsamples*

**Figure S20:** *Significant clusters across cohort-wise analyses (replicability analysis) for **model 15** – CTQ extreme severity compared to none in MDD subsamples*

**Figure S21:** *Significant clusters across cohort-wise analyses (replicability analysis) for **model 1** – CTQ sum in HC and MDD samples controlling for MDD diagnosis – **sex-stratified for female subsample***

**Figure S22:** *Significant clusters across cohort-wise analyses (replicability analysis) for **model 2** – CTQ sum as a predictor in HC and MDD samples without controlling for MDD diagnosis – **sex-stratified for female subsample***

**Figure S23:** *Significant clusters across cohort-wise analyses (replicability analysis) for* ***model 3*** *– CTQ sum as a predictor in HC subsamples –* ***sex-stratified for female subsample***

**Figure S24:** *Significant clusters across cohort-wise analyses (replicability analysis) for* ***model 4*** *– CTQ sum as a predictor in MDD subsamples –* ***sex-stratified for female subsample***

**Figure S25:** *Significant clusters across cohort-wise analyses (replicability analysis) for* ***model 5*** *– CTQ abuse subscales as a predictor in HC and MDD samples controlling for MDD diagnosis –* ***sex-stratified for female subsample***

**Figure S26:** *Significant clusters across cohort-wise analyses (replicability analysis) for* ***model 6*** *– CTQ neglect subscales as a predictor in HC and MDD samples controlling for MDD diagnosis –* ***sex-stratified for female subsample***

**Figure S27:** *Significant clusters across cohort-wise analyses (replicability analysis) for* ***model 7*** *– CTQ subscale emotional abuse as a predictor in HC and MDD samples controlling for MDD diagnosis –* ***sex-stratified for female subsample***

**Figure S28:** *Significant clusters across cohort-wise analyses (replicability analysis) for* ***model 8*** *– CTQ subscale physical abuse as a predictor in HC and MDD samples controlling for MDD diagnosis –* ***sex-stratified for female subsample***

**Figure S29:** *Significant clusters across cohort-wise analyses (replicability analysis) for* ***model 9*** *– CTQ subscale sexual abuse as a predictor in HC and MDD samples controlling for MDD diagnosis –* ***sex-stratified for female subsample***

**Figure S30:** *Significant clusters across cohort-wise analyses (replicability analysis) for* ***model 10*** *– CTQ subscale emotional neglect as a predictor in HC and MDD samples controlling for MDD diagnosis –* ***sex-stratified for female subsample***

**Figure S31:** *Significant clusters across cohort-wise analyses (replicability analysis) for* ***model 11*** *– CTQ subscale physical neglect as a predictor in HC and MDD samples controlling for MDD diagnosis –* ***sex-stratified for female subsample***

**Figure S32:** *Significant clusters across cohort-wise analyses (replicability analysis) for* ***model 12*** *– CTQ extreme severity compared to none in HC and MDD samples controlling for MDD diagnosis –* ***sex-stratified for female subsample***

**Figure S33:** *Significant clusters across cohort-wise analyses (replicability analysis) for* ***model 13*** *– CTQ extreme severity compared to none in HC and MDD samples without controlling for MDD diagnosis –* ***sex-stratified for female subsample***

**Figure S34:** *Significant clusters across cohort-wise analyses (replicability analysis) for* ***model 14*** *– CTQ extreme severity compared to none in HC subsamples –* ***sex-stratified for female subsample***

**Figure S35:** *Significant clusters across cohort-wise analyses (replicability analysis) for* ***model 15*** *– CTQ extreme severity compared to none in MDD subsamples –* ***sex-stratified for female subsample***

**Figure S36:** *Significant clusters across cohort-wise analyses (replicability analysis) for* ***model 1*** *– CTQ sum in HC and MDD samples controlling for MDD diagnosis –* ***sex-stratified for male subsample***

**Figure S37:** *Significant clusters across cohort-wise analyses (replicability analysis) for* ***model 2*** *– CTQ sum as a predictor in HC and MDD samples without controlling for MDD diagnosis –* ***sex-stratified for male subsample***

**Figure S38:** *Significant clusters across cohort-wise analyses (replicability analysis) for* ***model 3*** *– CTQ sum as a predictor in HC subsamples –* ***sex-stratified for male subsample***

**Figure S39:** *Significant clusters across cohort-wise analyses (replicability analysis) for* ***model 4*** *– CTQ sum as a predictor in MDD subsamples –* ***sex-stratified for male subsample***

**Figure S40:** *Significant clusters across cohort-wise analyses (replicability analysis) for* ***model 5*** *– CTQ abuse subscales as a predictor in HC and MDD samples controlling for MDD diagnosis –* ***sex-stratified for male subsample***

**Figure S41:** *Significant clusters across cohort-wise analyses (replicability analysis) for* ***model 6*** *– CTQ neglect subscales as a predictor in HC and MDD samples controlling for MDD diagnosis –* ***sex-stratified for male subsample***

**Figure S42:** *Significant clusters across cohort-wise analyses (replicability analysis) for* ***model 7*** *– CTQ subscale emotional abuse as a predictor in HC and MDD samples controlling for MDD diagnosis –* ***sex-stratified for male subsample***

**Figure S43:** *Significant clusters across cohort-wise analyses (replicability analysis) for* ***model 8*** *– CTQ subscale physical abuse as a predictor in HC and MDD samples controlling for MDD diagnosis –* ***sex-stratified for male subsample***

**Figure S44:** *Significant clusters across cohort-wise analyses (replicability analysis) for* ***model 9*** *– CTQ subscale sexual abuse as a predictor in HC and MDD samples controlling for MDD diagnosis –* ***sex-stratified for male subsample***

**Figure S45:** *Significant clusters across cohort-wise analyses (replicability analysis) for* ***model 10*** *– CTQ subscale emotional neglect as a predictor in HC and MDD samples controlling for MDD diagnosis –* ***sex-stratified for male subsample***

**Figure S46:** *Significant clusters across cohort-wise analyses (replicability analysis) for* ***model 11*** *– CTQ subscale physical neglect as a predictor in HC and MDD samples controlling for MDD diagnosis –* ***sex-stratified for male subsample***

**Figure S47:** *Significant clusters across cohort-wise analyses (replicability analysis) for* ***model 12*** *– CTQ extreme severity compared to none in HC and MDD samples controlling for MDD diagnosis –* ***sex-stratified for male subsample***

**Figure S48:** *Significant clusters across cohort-wise analyses (replicability analysis) for* ***model 13*** *– CTQ extreme severity compared to none in HC and MDD samples without controlling for MDD diagnosis –* ***sex-stratified for male subsample***

**Figure S49:** *Significant clusters across cohort-wise analyses (replicability analysis) for* ***model 14*** *– CTQ extreme severity compared to none in HC subsamples –* ***sex-stratified for male subsample***

**Figure S50:** *Significant clusters across cohort-wise analyses (replicability analysis) for* ***model 15*** *– CTQ extreme severity compared to none in MDD subsamples –* ***sex-stratified for male subsample***

*Cohort characteristics and inclusion criteria*

The existing Marburg-Münster Affective Disorders Cohort Study (MACS), Münster Neuroimaging Cohort (MNC) and BiDirect study were utilized for analysis. The cohorts include adult participants with age 18-65 years (35-65 years for the BiDirect study) and were all created to investigate affective disorders. Largest groups within each of the three cohorts comprise healthy control participants (HC) free from any lifetime history of mental disorders and participants with a major depressive disorder (MDD) diagnosis. Additional smaller patient groups with bipolar disorder (MACS and MNC), psychosis spectrum disorders (schizoaffective and schizophrenia; MACS), as well as comorbid cardiovascular diseases (BiDirect) were specifically recruited for the cohorts, and psychiatric comorbidities were generally permitted with some exceptions regarding brain organic disorders or substance use disorders. MACS and BiDirect recruited patient populations from psychiatric hospitals and outpatient centers in and around Münster, Germany (MACS and BiDirect) and Marburg, Germany (MACS). For the MNC, inpatient populations were included only, recruited also at psychiatric hospitals in Münster, Germany.

*Scanner harmonization procedure*

Data from different MRI scanners was included in the current analysis. In addition, data collection of the available cohorts took place over the course of several years (MNC being the oldest and still ongoing study, with study start in October 2009). Thus, hardware changes had to be conducted over the course of the study within the MACS and MNC study, potentially contributing to further scanner hardware effects. Notably, the three cohorts utilized here were already closely aligned with regard to scanning sequence parameters, thus reducing scanner effects to a minimum. The neuroCombat harmonization tool aims to neutralize scanner effects by estimating regional location (i.e., mean) and scaling (i.e., variance) parameters across all scanner groups (Fortin et al., 2018; https://github.com/Jfortin1/neuroCombat). By default, these parameters estimated using empirical bayes, thus pooling information across voxels. These parameters are then applied to adjust voxel intensities to achieve comparable distributions across scanner groups. This procedure allows the specification of “biological covariates”, for which associated variability will be preserved from the harmonization. This is particularly important when scanner groups differ with regard to variables that are associated with the biological outcome (e.g., age). As recommended in the Combat manual we included CTQ sum (as the primary variable of interest), as well as all utilized covariates (i.e., age, sex, total intracranial volume and MDD diagnosis) as biological covariates to avoid that the variability of gray matter intensities associated with these variables is ‘harmonized out’.

*Measures for clinical characterization*

A Structured Clinical Interview for the DSM-IV (SCID) (Wittchen et al., 1997) was used within MACS and MNC, while the Mini International Neuropsychiatric Interview (MINI) (Lecrubier et al., 1997) was used within the BiDirect cohort. Both measures, the SCID and the MINI, were used to deduct lifetime psychiatric diagnoses. The current remission status of MDD patients was assessed using the SCID for MACS and MNC. For the BiDirect cohort a CES-D cutoff was used as a proxy to categorize patients into remitted (CES-D < 20) or acute.

For additional clinical characterization regarding the current depression severity the Beck Depression Inventory (BDI) (Beck & Steer, 1987) was available for MACS and MNC cohorts, while the Center of Epidemiological Studies Depression Scale (CES-D) (Eaton et al., 2004) was available for the BiDirect cohort. The 17-item version of the Hamilton Depression Rating Scale (HDRS-17) was available in all three cohorts as a standardized clinical rating instrument for depression severity (Hamilton, 1960). Further, an additional interview-based characterization of the previous course of disease was available for MACS and MNC regarding estimates for the total number of psychiatric hospitalization and the number of previous depressive episodes. No comparable measure for the previous disease course was available for the BiDirect sample. For a summary of the clinical characterization of all cohorts see Table S1.

**Table S1**

*Demographic and clinical characteristics across all three cohorts*

| | HC | | | MDD | | |
|---|---|---|---|---|---|---|
| | count | mean | SD | count | mean | SD |
| *MACS (n=1752)* | | | | | | |
| Age | 930 | 34.23 | 12.96 | 822 | 36.50 | 13.13 |
| Sex (f/m) | 597/333 | | | 537/285 | - | - |
| Education years | 927 | 13.98 | 2.57 | 818 | 13.20 | 2.71 |
| Remission (y/n) | - | - | - | 452/369 | - | - |
| HDRS-17 | 926 | 1.38 | 2.04 | 819 | 8.43 | 6.47 |
| BDI | 926 | 4.11 | 4.28 | 811 | 17.59 | 11.12 |
| CES-D | - | - | - | - | - | - |
| Age of onset | - | - | - | 817 | 25.82 | 12.49 |
| # Depressive episodes | - | - | - | 782 | 3.87 | 6.08 |
| # Inpatient treatments | - | - | - | 813 | 1.55 | 1.98 |
| | count | mean | SD | count | mean | SD |
| *MNC (n=916)* | | | | | | |
| Age | 647 | 34.79 | 12.12 | 269 | 36.99 | 13.36 |
| Sex (f/m) | 359/288 | | | 148/121 | - | - |
| Education years | 643 | 14.64 | 2.51 | 268 | 13.73 | 2.71 |
| Remission (y/n) | - | - | - | 1/268 | - | - |
| HDRS-17 | 520 | 0.85 | 1.40 | 267 | 19.21 | 4.12 |
| BDI | 647 | 2.32 | 3.12 | 266 | 28.20 | 9.14 |
| CES-D | - | - | - | - | - | - |
| Age of onset | - | - | - | 260 | 25.56 | 12.78 |
| # Depressive episodes | - | - | - | 269 | 4.42 | 5.39 |
| # Inpatient treatments | - | - | - | 249 | 2.45 | 2.71 |
| | count | mean | SD | count | mean | SD |
| *BiDirect (n=557)* | | | | | | |
| Age | 321 | 52.61 | 7.97 | 236 | 50.63 | 7.06 |
| Sex (f/m) | 171/150 | | | 143/93 | - | - |
| Education years | 321 | 15.21 | 2.59 | 234 | 14.35 | 2.40 |
| Remission (y/n) | - | - | - | 75/161 | - | - |
| HDRS-17 | - | - | - | 236 | 13.08 | 6.61 |
| BDI | - | - | - | - | - | - |
| CES-D | 321 | 7.84 | 5.90 | 236 | 26.33 | 12.36 |
| Age of onset | - | - | - | - | - | - |
| # Depressive episodes | - | - | - | 225 | 5.89 | 9.49 |
| # Inpatient treatments | - | - | - | 233 | 1.56 | 1.30 |

*Note*. HC, healthy controls; MDD, major depressive disorder; MACS, Marburg Münster Affective Disorders Cohort Study; MNC, Münster Neuroimaging Cohort; BiDirect, BiDirect study cohort; f, female; m, male; y, yes; n, no; HDRS-17, 17-item Hamilton Depression Rating Scale; BDI, Beck Depression Inventory; CES-D, Center for Epidemiologic Studies Depression Scale.

**Table S2**

*Summary statistics for reported experiences of childhood maltreatment in the CTQ across cohorts*

| | HC | | | MDD | | |
|---|---|---|---|---|---|---|
| | count | mean | SD | count | mean | SD |
| *MACS (n=1725)* | | | | | | |
| CTQ sum | 930 | 32.68 | 8.82 | 822 | 45.13 | 15.65 |
| EA subscale sum | 930 | 7.06 | 2.88 | 822 | 10.96 | 5.11 |
| PA subscale sum | 930 | 5.60 | 1.54 | 822 | 6.78 | 3.14 |
| SA subscale sum | 930 | 5.29 | 1.48 | 822 | 6.38 | 3.43 |
| EN subscale sum | 930 | 8.53 | 3.72 | 822 | 13.18 | 5.36 |
| PN subscale sum | 930 | 6.20 | 1.89 | 822 | 7.82 | 3.09 |
| | count | mean | SD | count | mean | SD |
| *MNC (n=916)* | | | | | | |
| CTQ sum | 647 | 32.90 | 8.14 | 269 | 46.91 | 17.06 |
| EA subscale sum | 647 | 6.98 | 2.70 | 269 | 11.16 | 5.44 |
| PA subscale sum | 647 | 5.59 | 1.72 | 269 | 7.40 | 3.90 |
| SA subscale sum | 647 | 5.22 | 0.94 | 269 | 6.46 | 3.82 |
| EN subscale sum | 647 | 8.80 | 3.64 | 269 | 13.60 | 5.54 |
| PN subscale sum | 647 | 6.31 | 2.00 | 269 | 8.29 | 3.27 |
| | count | mean | SD | count | mean | SD |
| *BiDirect (n=557)* | | | | | | |
| CTQ sum | 321 | 32.69 | 7.64 | 236 | 44.97 | 17.14 |
| EA subscale sum | 321 | 6.42 | 2.17 | 236 | 10.06 | 5.20 |
| PA subscale sum | 321 | 5.85 | 1.51 | 236 | 6.90 | 3.45 |
| SA subscale sum | 321 | 5.19 | 0.86 | 236 | 6.36 | 3.59 |
| EN subscale sum | 321 | 8.95 | 3.97 | 236 | 13.88 | 5.68 |
| PN subscale sum | 321 | 6.27 | 1.97 | 236 | 7.78 | 3.00 |

*Note*. CTQ, Childhood Trauma Questionnaire; HC, healthy controls; MDD, major depressive disorder; MACS, Marburg Münster Affective Disorders Cohort Study; MNC, Münster Neuroimaging Cohort; BiDirect, BiDirect study cohort; EA, emotional abuse; PA, physical abuse; SA, sexual abuse; PN, physical neglect; EN, emotional neglect.

**Table S3**

*Pairwise cohort comparison in sample characteristics including HC and MDD participants*

| | MACS-MNC | | MACS-BiDirect | | MNC-BiDirect | |
|---|---|---|---|---|---|---|
| | Test statistic | p-value | Test statistic | p-value | Test statistic | p-value |
| Sex | 21.9432 | <.001 | 25.7993 | <.001 | 0.9449 | .331 |
| MDD diagnosis | 75.9345 | <.001 | 3.3405 | .068 | 25.4202 | <.001 |
| Age | 813601 | .554 | 811138 | <.001 | 79976 | <.001 |
| Education years | 934558 | <.001 | 599155 | <.001 | 235718 | .024 |
| CTQ sum | 753976 | 0.010 | 475857 | .378 | 245900 | .244 |
| CTQ EA | 727820 | <.001 | 414523 | <.001 | 271148 | .038 |
| CTQ PA | 772544 | .054 | 521764 | .004 | 227639 | <.001 |
| CTQ SA | 776172 | .021 | 476570 | .176 | 252777 | .605 |
| CTQ EN | 763920 | .041 | 501968 | .304 | 235560 | .013 |
| CTQ PN | 788585 | .445 | 482996 | .707 | 253297 | .811 |

*Note.* Cohort differences in the distribution of categorical variables (sex and MDD diagnosis) were tested using $chi^2$ tests (the $chi^2$ value is given as a test statistic). Cohort mean differences in continuous variables were tested using Mann-Whitney-U tests (U-values are given as test statistic) due to normality violations in all continuous variables according to Shapiro-Wilk tests significant at $p<.05$. MDD, major depressive disorder; MACS, Marburg Münster Affective Disorders Cohort Study; MNC, Münster Neuroimaging Cohort; BiDirect, BiDirect study cohort; EA, emotional abuse; PA, physical abuse; SA, sexual abuse; PN, physical neglect; EN, emotional neglect.

**Table S4**

*Pairwise cohort comparison of clinical characteristics of the MDD subgroups*

| | MACS-MNC | | MACS-BiDirect | | MNC-BiDirect | |
|---|---|---|---|---|---|---|
| | Test statistic | p-value | Test statistic | p-value | Test statistic | p-value |
| Remission | 247.2073 | <.001 | 38.7962 | <.001 | 94.5569 | <.001 |
| HDRS-17 | 20555.5 | <.001 | 58872 | <.001 | 48559 | <.001 |
| BDI | 50433 | <.001 | - | - | - | - |
| Age of onset | 108473 | .6044 | - | - | - | - |
| # Depressive episodes | 91229 | <.001 | 75183 | <.001 | 29558.5 | 0.651 |
| # Inpatient treatments | 67923.5 | <.001 | 82246 | .001 | 36381 | <.001 |

*Note*. Cohort differences in the distribution of categorical variables (remission) were tested using $chi^2$ tests (the $chi^2$ value is given as a test statistic). Cohort mean differences in continuous variables were tested using Mann-Whitney-U tests (U-values are given as test statistic) due to normality violations in all continuous variables according to Shapiro-Wilk tests significant at $p<.05$. CES-D values were not compared across cohorts as they were only available for the BiDirect cohort. MDD, major depressive disorder; MACS, Marburg Münster Affective Disorders Cohort Study; MNC, Münster Neuroimaging Cohort; BiDirect, BiDirect study cohort; HDRS-17, 17-item Hamilton Depression Rating Scale; BDI, Beck Depression Inventory.

**Table S5**

*Differences in CTQ scales across HC and MDD groups in pooled sample and cohort-wise*

| Cohort | CTQ scale | U Statistic | p-value | Effect size r |
|---|---|---|---|---|
| Pooled | CTQ sum | 569455.5 | <0.001 | 0.548 |
| | CTQ EA | 628867.5 | <0.001 | 0.501 |
| | CTQ PA | 974777.5 | <0.001 | 0.226 |
| | CTQ SA | 1052632.5 | <0.001 | 0.164 |
| | CTQ EN | 608515.5 | <0.001 | 0.517 |
| | CTQ PN | 794500 | <0.001 | 0.369 |
| MACS | CTQ sum | 169297 | <0.001 | 0.557 |
| | CTQ EA | 187176 | <0.001 | 0.510 |
| | CTQ PA | 296571 | <0.001 | 0.224 |
| | CTQ SA | 315886.5 | <0.001 | 0.174 |
| | CTQ EN | 181692 | <0.001 | 0.525 |
| | CTQ PN | 241310 | <0.001 | 0.369 |
| MNC | CTQ sum | 37554.5 | <0.001 | 0.568 |
| | CTQ EA | 42979.5 | <0.001 | 0.506 |
| | CTQ PA | 62214 | <0.001 | 0.285 |
| | CTQ SA | 73644 | <0.001 | 0.154 |
| | CTQ EN | 41962 | <0.001 | 0.518 |
| | CTQ PN | 51077 | <0.001 | 0.413 |
| BiDirect | CTQ sum | 18761 | <0.001 | 0.505 |
| | CTQ EA | 20450 | <0.001 | 0.460 |
| | CTQ PA | 32585.5 | 0.0014 | 0.140 |
| | CTQ SA | 32396 | <0.001 | 0.145 |
| | CTQ EN | 18389 | <0.001 | 0.515 |
| | CTQ PN | 24819 | <0.001 | 0.345 |

*Note. Mann-Whitney U tests were calculated to test if differences in CTQ scales were significant across HC and MDD groups. A rank-biserial correlation coefficient is given as an effect size for this difference.*

**Table S6**

*Results summary for pooled cohorts (n=3225) at a significance level of $p_{FWE}$<.05 –* ***sex-stratified for female subsample***

| Model | k significant | Partial $R^2$ Min | Partial $R^2$ Max | Main regions |
|---|---|---|---|---|
| Model 1 | 0 | - | - | - |
| Model 2 | 632 | 0.010 | 0.014 | Temporal Mid/Sup R, Cerebellum R, Insula L |
| Model 3 | 212 | 0.017 | 0.021 | Insula L+R, Temporal Inf Orb 2 L, Temporal Pole Sup R, Temporal Mid/Sup R |
| Model 4 | 0 | - | - | - |
| Model 5 | 0 | - | - | - |
| Model 6 | 0 | - | - | - |
| Model 7 | 0 | - | - | - |
| Model 8 | 0 | - | - | - |
| Model 9 | 0 | - | - | - |
| Model 10 | 0 | - | - | - |
| Model 11 | 0 | - | - | - |
| Model 12 | 1847 | 0.017 | 0.025 | Cerebellum L+R, Temporal Mid L, Rectus R, Caudate R, Insula R, Temporal Pole Sup R |
| Model 13 | 6343 | 0.017 | 0.029 | Temporal Sup L+R, Temporal Pole Sup R, Lingual L, Amygdala R, Rectus L, Lingual R |
| Model 14 | 566 | 0.029 | 0.041 | Insula L+R, Cerebellum L, Temporal Mid L, Frontal Inf Orb 2 L, Temporal Mid/Sup L+R |
| Model 15 | 15 | 0.041 | 0.045 | 'no label' |

*Note*. Number of significant voxels at $p_{FWE}$<.05 are shown, as well as their minimum and maximum effect size for each analysis. Cluster labelling was conducted based on the aal atlas using the atlasreader python package (Notter et al., 2019). Main regions are reported. L, left; R, right; Mid, middle; Sup, superior; Inf, inferior; Orb orbital.

**Table S7**

*Results summary for pooled cohorts (n=3225) at a significance level of $p_{FWE}$<.05 – **sex-stratified for male subsample***

| | | partial $R^2$ | | |
|---|---|---|---|---|
| Model | k significant | Min | Max | Main regions |
| Model 1 | 0 | - | - | - |
| Model 2 | 312 | 0.015 | 0.020 | Fusiform L+R, Lingual L, Temporal Sup R |
| Model 3 | 0 | - | - | - |
| Model 4 | 0 | - | - | - |
| Model 5 | 0 | - | - | - |
| Model 6 | 0 | - | - | - |
| Model 7 | 0 | - | - | - |
| Model 8 | 0 | - | - | - |
| Model 9 | 0 | - | - | - |
| Model 10 | 0 | - | - | - |
| Model 11 | 0 | - | - | - |
| Model 12 | 0 | - | - | - |
| Model 13 | 9 | 0.028 | 0.030 | Fusiform L |
| Model 14 | 0 | - | - | - |
| Model 15 | 0 | - | - | - |

*Note*. Number of significant voxels at $p_{FWE}$<.05 are shown, as well as their minimum and maximum effect size for each analysis. Cluster labelling was conducted based on the aal atlas using the atlasreader python package (Notter et al., 2019). Main regions are reported. L, left; R, right; Sup, superior.

**Table S8**

*Results summary of single cohorts at a cohort-wise significance level of* $\boldsymbol{p_{unc}<.001}$

| Model | Cohort | k significant | Partial $R^2$ Min | Partial $R^2$ Max | Main regions |
|---|---|---|---|---|---|
| Model 1 | MACS | 0 | - | - | - |
| Model 1 | MNC | 8 | 0.01 | 0.011 | Precentral L, Supramarginal R |
| Model 1 | BiDirect | 1299 | 0.017 | 0.029 | Cerebellum L, Temporal Sup R |
| Model 2 | MACS | 1614 | 0.005 | 0.011 | Temporal Mid/Sup R, Rectus R |
| Model 2 | MNC | 378 | 0.01 | 0.018 | Thalamus L, Hippocampus L |
| Model 2 | BiDirect | 3388 | 0.017 | 0.037 | Cerebellum L, Lingual L |
| Model 3 | MACS | 63 | 0.01 | 0.014 | Frontal Mid 2 R, Frontal Inf Oper R |
| Model 3 | MNC | 646 | 0.015 | 0.026 | Precentral L, Cerebellum R |
| Model 3 | BiDirect | 36 | 0.03 | 0.032 | Rectus L, Cerebellum L |
| Model 4 | MACS | 42 | 0.012 | 0.014 | Cerebellum R, Cerebellum L |
| Model 4 | MNC | 0 | - | - | - |
| Model 4 | BiDirect | 630 | 0.04 | 0.071 | Cerebellum L, Temporal Sup R |
| Model 5 | MACS | 0 | - | - | - |
| Model 5 | MNC | 5 | 0.011 | 0.011 | Postcentral R |
| Model 5 | BiDirect | 377 | 0.017 | 0.023 | Cerebellum L, Temporal Sup R |
| Model 6 | MACS | 0 | - | - | - |
| Model 6 | MNC | 38 | 0.01 | 0.014 | Supramarginal R |
| Model 6 | BiDirect | 1461 | 0.017 | 0.028 | Cerebellum L, Fusiform R |
| Model 7 | MACS | 0 | - | - | - |
| Model 7 | MNC | 0 | - | - | - |
| Model 7 | BiDirect | 375 | 0.017 | 0.023 | Cerebellum L, Precuneus L |
| Model 8 | MACS | 282 | 0.005 | 0.007 | Cerebellum R, Precentral R |
| Model 8 | MNC | 63 | 0.01 | 0.012 | Calcarine L, Postcentral R |
| Model 8 | BiDirect | 1283 | 0.017 | 0.026 | Cerebellum R, Precentral L |
| Model 9 | MACS | 0 | - | - | - |
| Model 9 | MNC | 102 | 0.01 | 0.012 | Frontal Mid 2 R, Precentral R |
| Model 9 | BiDirect | 491 | 0.017 | 0.024 | Cerebellum L, Lingual L |
| Model 10 | MACS | 21 | 0.005 | 0.006 | Cerebellum R |
| Model 10 | MNC | 10 | 0.01 | 0.013 | Supramarginal R |
| Model 10 | BiDirect | 356 | 0.017 | 0.026 | Cerebellum L, Temporal Mid L |
| Model 11 | MACS | 0 | - | - | - |
| Model 11 | MNC | 29 | 0.01 | 0.011 | 'no label' |
| Model 11 | BiDirect | 3233 | 0.017 | 0.032 | Cerebellum L, Supramarginal R |
| Model 12 | MACS | 0 | - | - | - |
| Model 12 | MNC | 246 | 0.019 | 0.025 | Frontal Sup 2 R, Cerebellum R |
| Model 12 | BiDirect | 956 | 0.031 | 0.042 | Cerebellum L, Supramarginal R |
| Model 13 | MACS | 2403 | 0.01 | 0.017 | Temporal Sup R, Rectus R |
| Model 13 | MNC | 675 | 0.019 | 0.033 | Thalamus R, Temporal Sup R |
| Model 13 | BiDirect | 3031 | 0.031 | 0.057 | Cerebellum L, Supramarginal R |
| Model 14 | MACS | 45 | 0.017 | 0.02 | Temporal Inf R, Lingual L |

| | | | | | |
|---|---|---|---|---|---|
| Model 14 | MNC | 481 | 0.026 | 0.037 | Temporal Sup R, Cerebellum R |
| Model 14 | BiDirect | 146 | 0.052 | 0.074 | Rectus L, Precentral L |
| Model 15 | MACS | 59 | 0.022 | 0.027 | Cerebellum L |
| Model 15 | MNC | 1135 | 0.064 | 0.135 | Frontal Sup 2 R, Rectus R |
| Model 15 | BiDirect | 261 | 0.074 | 0.113 | Frontal Inf Orb 2 R, Parietal Sup R |

*Note*. Number of significant voxels at uncorrected p<.001 are shown, as well as their minimum and maximum effect size for each analysis. Cluster labelling was conducted based on the aal atlas using the atlasreader python package (Notter et al., 2019). Main regions of largest clusters are reported. MACS, Marburg Münster Affective Disorders Cohort Study; MNC, Münster Neuroimaging Cohort; BiDirect, BiDirect study cohort.

**Table S9**

*Results summary of single cohorts at a cohort-wise significance level of* $\boldsymbol{p_{unc}}$***<.001 – sex-stratified for female subsample***

| Model | Cohort | k significant | Partial $R^2$ Min | Partial $R^2$ Max | Main regions |
|---|---|---|---|---|---|
| Model 1 | MACS | 0 | - | - | - |
| Model 1 | MNC | 60 | 0.019 | 0.023 | Lingual R, Cerebellum R |
| Model 1 | BiDirect | 535 | 0.033 | 0.051 | Temporal Inf L, Cerebellum L |
| Model 2 | MACS | 1367 | 0.008 | 0.017 | Temporal Sup R, Cerebellum R |
| Model 2 | MNC | 270 | 0.019 | 0.025 | Precentral L, Lingual L |
| Model 2 | BiDirect | 1001 | 0.032 | 0.050 | Cerebellum L, Lingual L |
| Model 3 | MACS | 358 | 0.016 | 0.027 | Temporal Mid/Sup L, Frontal Mid 2 R |
| Model 3 | MNC | 1838 | 0.027 | 0.051 | Temporal Sup R, Temporal Pole Sup L |
| Model 3 | BiDirect | 26 | 0.063 | 0.070 | Parahippocampal R, Temporal Mid L |
| Model 4 | MACS | 0 | - | - | - |
| Model 4 | MNC | 70 | 0.064 | 0.072 | Postcentral R, Occipital Mid L |
| Model 4 | BiDirect | 263 | 0.066 | 0.086 | OFC Post R, Frontal Inf Orb 2 R |
| Model 5 | MACS | 0 | - | - | - |
| Model 5 | MNC | 192 | 0.019 | 0.032 | Frontal Mid 2 R, Temporal Pole Sup R |
| Model 5 | BiDirect | 128 | 0.033 | 0.041 | Temporal Sup R, Temporal Inf L |
| Model 6 | MACS | 65 | 0.008 | 0.010 | Temporal Sup R |
| Model 6 | MNC | 116 | 0.019 | 0.023 | Rectus R, Lingual R |
| Model 6 | BiDirect | 628 | 0.033 | 0.050 | OFC Post R, Temporal Inf L |
| Model 7 | MACS | 0 | - | - | - |
| Model 7 | MNC | 284 | 0.019 | 0.025 | Frontal Mid 2 R, Temporal Pole Sup R |
| Model 7 | BiDirect | 247 | 0.033 | 0.044 | OFC Post R, Temporal Mid R |
| Model 8 | MACS | 99 | 0.008 | 0.010 | Cerebellum L, Precentral R |
| Model 8 | MNC | 960 | 0.019 | 0.030 | Calcarine L, Cuneus L |
| Model 8 | BiDirect | 474 | 0.033 | 0.051 | Paracentral Lobule R, Postcentral R |
| Model 9 | MACS | 0 | - | - | - |
| Model 9 | MNC | 42 | 0.019 | 0.024 | Thalamus R |
| Model 9 | BiDirect | 1045 | 0.033 | 0.051 | Cerebellum R, Parietal Inf L |
| Model 10 | MACS | 393 | 0.008 | 0.012 | Temporal Sup R |
| Model 10 | MNC | 116 | 0.019 | 0.023 | Rectus R, Cingulate Mid L |
| Model 10 | BiDirect | 317 | 0.033 | 0.047 | OFC Post R, Frontal Inf Orb 2 R |
| Model 11 | MACS | 0 | - | - | - |
| Model 11 | MNC | 164 | 0.019 | 0.026 | Occipital Inf R, Fusiform R |
| Model 11 | BiDirect | 1037 | 0.033 | 0.060 | Frontal Inf Orb 2 L, Temporal Sup R |
| Model 12 | MACS | 819 | 0.015 | 0.022 | Cerebellum L, Cerebellum R |
| Model 12 | MNC | 1381 | 0.033 | 0.051 | Parietal Sup L, Cerebellum L |
| Model 12 | BiDirect | 1107 | 0.054 | 0.084 | Cerebellum L, Temporal Inf L |
| Model 13 | MACS | 5010 | 0.015 | 0.031 | Temporal Sup R, Cerebellum R |
| Model 13 | MNC | 209 | 0.033 | 0.047 | Cerebellum R, Precentral L |
| Model 13 | BiDirect | 1133 | 0.054 | 0.081 | Cerebellum L, Supramarginal R |
| Model 14 | MACS | 101 | 0.026 | 0.035 | Insula R, Temporal Mid L |
| Model 14 | MNC | 1023 | 0.046 | 0.066 | Temporal Sup R, Parietal Sup R |
| Model 14 | BiDirect | 1160 | 0.111 | 0.199 | Temporal Inf R, Temporal Mid L |
| Model 15 | MACS | 8 | 0.033 | 0.036 | ‘no label’ |

| | | | | | |
|---|---|---|---|---|---|
| Model 15 | MNC | 269 | 0.111 | 0.162 | Frontal Sup 2 R, Occipital Mid R |
| Model 15 | BiDirect | 865 | 0.103 | 0.149 | Cerebellum L, Frontal Sup 2 R |

*Note*. Number of significant voxels at uncorrected p<.001 are shown, as well as their minimum and maximum effect size for each analysis. Cluster labelling was conducted based on the aal atlas using the atlasreader python package (Notter et al., 2019). Main regions of largest clusters are reported. MACS, Marburg Münster Affective Disorders Cohort Study; MNC, Münster Neuroimaging Cohort; BiDirect, BiDirect study cohort.

**Table S10**

*Results summary of single cohorts at a cohort-wise significance level of* ***$p_{unc}$<.001 – sex-stratified for male subsample***

| | | | Partial $R^2$ | | |
|---|---|---|---|---|---|
| Model | Cohort | k significant | Min | Max | Main regions |
| Model 1 | MACS | 10 | 0.015 | 0.017 | Paracentral Lobule R |
| Model 1 | MNC | 87 | 0.023 | 0.033 | Hippocampus L, Rectus L |
| Model 1 | BiDirect | 718 | 0.036 | 0.061 | Postcentral R, Temporal Mid L |
| Model 2 | MACS | 522 | 0.015 | 0.022 | Temporal Mid R, Paracentral Lobule R |
| Model 2 | MNC | 268 | 0.023 | 0.047 | Rectus L, Thalamus L |
| Model 2 | BiDirect | 989 | 0.036 | 0.065 | Postcentral R, Fusiform R |
| Model 3 | MACS | 71 | 0.029 | 0.034 | Postcentral R, Calcarine L |
| Model 3 | MNC | 159 | 0.033 | 0.041 | Cerebellum R |
| Model 3 | BiDirect | 163 | 0.055 | 0.075 | Postcentral R, Cerebellum R |
| Model 4 | MACS | 0 | - | - | - |
| Model 4 | MNC | 0 | - | - | - |
| Model 4 | BiDirect | 1317 | 0.101 | 0.177 | Precuneus R, Occipital Sup R |
| Model 5 | MACS | 110 | 0.015 | 0.021 | Paracentral Lobule R |
| Model 5 | MNC | 92 | 0.023 | 0.028 | Hippocampus L, Supramarginal R |
| Model 5 | BiDirect | 1766 | 0.036 | 0.074 | Postcentral L, Cingulate Ant L |
| Model 6 | MACS | 47 | 0.016 | 0.023 | Frontal Sup 2 L, Postcentral L |
| Model 6 | MNC | 23 | 0.023 | 0.029 | Rectus L |
| Model 6 | BiDirect | 133 | 0.036 | 0.050 | Postcentral R, Cerebellum R |
| Model 7 | MACS | 100 | 0.015 | 0.019 | Temporal Sup R, Postcentral R |
| Model 7 | MNC | 26 | 0.023 | 0.034 | Rectus L |
| Model 7 | BiDirect | 1441 | 0.036 | 0.064 | Postcentral L, Temporal Mid L |
| Model 8 | MACS | 193 | 0.015 | 0.024 | Paracentral Lobule R |
| Model 8 | MNC | 93 | 0.023 | 0.028 | Thalamus L, Cerebellum L |
| Model 8 | BiDirect | 1945 | 0.036 | 0.069 | Occipital Sup R, Precentral L |
| Model 9 | MACS | 0 | - | - | - |
| Model 9 | MNC | 2219 | 0.023 | 0.037 | Occipital Mid R, Supramarginal R |
| Model 9 | BiDirect | 1 | 0.036 | 0.036 | Frontal Mid 2 R |
| Model 10 | MACS | 39 | 0.015 | 0.022 | Frontal Sup 2 L |
| Model 10 | MNC | 4 | 0.024 | 0.025 | 'no label' |
| Model 10 | BiDirect | 20 | 0.036 | 0.038 | Precuneus R, Occipital Mid L |
| Model 11 | MACS | 39 | 0.015 | 0.017 | Postcentral R, Paracentral Lobule R |
| Model 11 | MNC | 0 | - | - | - |
| Model 11 | BiDirect | 587 | 0.036 | 0.066 | Cerebellum R, Postcentral R |
| Model 12 | MACS | 0 | - | - | - |
| Model 12 | MNC | 154 | 0.043 | 0.050 | Frontal Sup 2 R, Fusiform R |
| Model 12 | BiDirect | 345 | 0.071 | 0.112 | Cuneus R, Frontal Inf Oper R |
| Model 13 | MACS | 7 | 0.028 | 0.030 | Paracentral Lobule R, Cuneus L |
| Model 13 | MNC | 575 | 0.043 | 0.082 | Rectus L, Thalamus L |
| Model 13 | BiDirect | 590 | 0.070 | 0.102 | Lingual R, Frontal Inf Tri R |
| Model 14 | MACS | 106 | 0.050 | 0.070 | Fusiform R, Occipital Mid L |
| Model 14 | MNC | 7 | 0.060 | 0.064 | Temporal Inf R, Cerebellum R |
| Model 14 | BiDirect | 44 | 0.098 | 0.119 | Insula L, Postcentral R |
| Model 15 | MACS | 0 | - | - | - |

| | | | | | |
|---|---|---|---|---|---|
| Model 15 | MNC | 715 | 0.146 | 0.240 | Frontal Sup 2 R, Frontal Sup Med L |
| Model 15 | BiDirect | 162 | 0.242 | 0.336 | Occipital Sup R, Calcarine L |

*Note*. Number of significant voxels at uncorrected p<.001 are shown, as well as their minimum and maximum effect size for each analysis. Cluster labelling was conducted based on the aal atlas using the atlasreader python package (Notter et al., 2019). Main regions of largest clusters are reported. MACS, Marburg Münster Affective Disorders Cohort Study; MNC, Münster Neuroimaging Cohort; BiDirect, BiDirect study cohort.

**Table S11**

*Results summary of single cohorts at a cohort-wise significance level of* $\boldsymbol{p_{unc}<.01}$

| Model | Cohort | k significant | Partial $R^2$ | | Main regions |
|---|---|---|---|---|---|
| | | | Min | Max | |
| Model 1 | MACS | 244 | 0.003 | 0.004 | Cerebellum R, Temporal Sup R |
| Model 1 | MNC | 749 | 0.006 | 0.011 | Precentral L, Supramarginal R |
| Model 1 | BiDirect | 10648 | 0.010 | 0.029 | Cerebellum L, Supramarginal R |
| Model 2 | MACS | 10400 | 0.003 | 0.011 | Temporal Sup R, Rectus R |
| Model 2 | MNC | 3444 | 0.006 | 0.018 | Hippocampus L, Thalamus L |
| Model 2 | BiDirect | 16924 | 0.010 | 0.037 | Cerebellum L, Supramarginal R |
| Model 3 | MACS | 1542 | 0.006 | 0.014 | Frontal Sup 2 R, Frontal Sup Medial L |
| Model 3 | MNC | 5230 | 0.008 | 0.026 | Cerebellum R, Precentral L |
| Model 3 | BiDirect | 2007 | 0.017 | 0.032 | Cerebellum L, Temporal Mid L |
| Model 4 | MACS | 1018 | 0.007 | 0.014 | Cerebellum L, Cerebellum R |
| Model 4 | MNC | 555 | 0.020 | 0.035 | Occipital Mid L, Frontal Inf Tri R |
| Model 4 | BiDirect | 8650 | 0.023 | 0.071 | Cerebellum L, Frontal Inf Orb 2 R |
| Model 5 | MACS | 309 | 0.003 | 0.004 | Cerebellum L, Precentral R |
| Model 5 | MNC | 1501 | 0.006 | 0.011 | Occipital Mid L, Precentral L |
| Model 5 | BiDirect | 8002 | 0.010 | 0.023 | Cerebellum L, Temporal Sup R |
| Model 6 | MACS | 152 | 0.003 | 0.004 | Cerebellum R, Temporal Sup R |
| Model 6 | MNC | 1251 | 0.006 | 0.014 | Precentral L, Supramarginal R |
| Model 6 | BiDirect | 10527 | 0.010 | 0.028 | Cerebellum L, Supramarginal R |
| Model 7 | MACS | 411 | 0.003 | 0.005 | Precentral R, Temporal Sup R |
| Model 7 | MNC | 1052 | 0.006 | 0.010 | Precentral L, Frontal Mid 2 R |
| Model 7 | BiDirect | 6301 | 0.010 | 0.023 | Cerebellum L, Temporal Sup R |
| Model 8 | MACS | 4652 | 0.003 | 0.007 | Cerebellum R, Cerebellum L |
| Model 8 | MNC | 3082 | 0.006 | 0.012 | Calcarine L, Postcentral R |
| Model 8 | BiDirect | 13399 | 0.010 | 0.026 | Cerebellum L, Postcentral L |
| Model 9 | MACS | 0 | - | - | - |
| Model 9 | MNC | 2888 | 0.006 | 0.012 | Frontal Mid 2 R, Thalamus L |
| Model 9 | BiDirect | 6123 | 0.010 | 0.024 | Cerebellum L, Temporal Pole Mid R |
| Model 10 | MACS | 3234 | 0.003 | 0.006 | Cerebellum L, Cerebellum R |
| Model 10 | MNC | 630 | 0.006 | 0.013 | Precentral L, Frontal Mid 2 L |
| Model 10 | BiDirect | 6556 | 0.010 | 0.026 | Cerebellum L, Cerebellum R |
| Model 11 | MACS | 25 | 0.003 | 0.004 | Occipital Sup L, Calcarine L |
| Model 11 | MNC | 1919 | 0.006 | 0.011 | Temporal Sup R, OFC Med L |
| Model 11 | BiDirect | 18007 | 0.010 | 0.032 | Supramarginal R, Temporal Sup L |
| Model 12 | MACS | 833 | 0.005 | 0.010 | Cerebellum R, Cerebellum L |
| Model 12 | MNC | 7423 | 0.011 | 0.025 | Occipital Mid L, Supramarginal R |
| Model 12 | BiDirect | 7313 | 0.018 | 0.042 | Cerebellum L, Supramarginal R |
| Model 13 | MACS | 18463 | 0.005 | 0.017 | Temporal Sup/Mid R, Cerebellum L |
| Model 13 | MNC | 7885 | 0.011 | 0.033 | Temporal Sup R, Caudate L |
| Model 13 | BiDirect | 14534 | 0.018 | 0.057 | Cerebellum L, Supramarginal R |
| Model 14 | MACS | 1313 | 0.010 | 0.020 | Temporal Inf R, Precuneus R |
| Model 14 | MNC | 8318 | 0.015 | 0.037 | Caudate L, Temporal Sup R |
| Model 14 | BiDirect | 4534 | 0.030 | 0.074 | Frontal Inf Tri L, Frontal Mid 2 L |

| | | | | | |
|---|---|---|---|---|---|
| Model 15 | MACS | 1182 | 0.012 | 0.027 | Cerebellum L, Cerebellum R |
| Model 15 | MNC | 9257 | 0.037 | 0.135 | Frontal Sup 2 R, Rectus R |
| Model 15 | BiDirect | 4980 | 0.043 | 0.113 | Cerebellum L, Temporal Sup R |

*Note*. Number of significant voxels at uncorrected p<.001 are shown, as well as their minimum and maximum effect size for each analysis. Cluster labelling was conducted based on the aal atlas using the atlasreader python package (Notter et al., 2019). Main regions of largest clusters are reported. MACS, Marburg Münster Affective Disorders Cohort Study; MNC, Münster Neuroimaging Cohort; BiDirect, BiDirect study cohort.

**Table S12**

*Results summary of single cohorts at a cohort-wise significance level of* ***$p_{unc}$<.01 – sex-stratified for female subsample***

| Model | Cohort | k significant | Partial $R^2$ | | Main regions |
|---|---|---|---|---|---|
| | | | Min | Max | |
| Model 1 | MACS | 341 | 0.005 | 0.008 | Temporal Sup R, Precentral R |
| Model 1 | MNC | 4962 | 0.011 | 0.023 | Lingual R, Calcarine L |
| Model 1 | BiDirect | 6369 | 0.019 | 0.051 | Cerebellum L, Temporal Sup R |
| Model 2 | MACS | 10960 | 0.005 | 0.017 | Temporal Sup R, Cerebellum R |
| Model 2 | MNC | 5658 | 0.011 | 0.025 | Lingual L, Precentral L |
| Model 2 | BiDirect | 8760 | 0.019 | 0.050 | Temporal Sup R, Cerebellum L |
| Model 3 | MACS | 3885 | 0.009 | 0.027 | Temporal Sup L, Frontal Mid 2 R |
| Model 3 | MNC | 15196 | 0.015 | 0.051 | Cerebellum L, Temporal Sup R |
| Model 3 | BiDirect | 2355 | 0.036 | 0.070 | Temporal Mid/Sup L, Lingual R |
| Model 4 | MACS | 364 | 0.010 | 0.017 | Cerebellum R, Cerebellum R |
| Model 4 | MNC | 2183 | 0.037 | 0.072 | Postcentral R, Occipital Mid L |
| Model 4 | BiDirect | 5523 | 0.038 | 0.086 | OFC Post R, Temporal Sup R |
| Model 5 | MACS | 703 | 0.005 | 0.008 | Cerebellum L, Precentral R |
| Model 5 | MNC | 5732 | 0.011 | 0.032 | Frontal Mid 2 R, Temporal Pole Sup R |
| Model 5 | BiDirect | 4823 | 0.019 | 0.041 | Temporal Sup R, Temporal Inf L |
| Model 6 | MACS | 890 | 0.005 | 0.010 | Temporal Sup R, Cerebellum 3 L |
| Model 6 | MNC | 4008 | 0.011 | 0.023 | Occipital Inf R, Cingulate Mid L |
| Model 6 | BiDirect | 7074 | 0.019 | 0.050 | Cerebellum L, OFC Post R |
| Model 7 | MACS | 684 | 0.005 | 0.007 | Cerebellum L, Cerebellum 8 R |
| Model 7 | MNC | 6563 | 0.011 | 0.025 | Temporal Sup L, Frontal Mid 2 R |
| Model 7 | BiDirect | 5404 | 0.019 | 0.044 | OFC Post R, Temporal Inf L |
| Model 8 | MACS | 3442 | 0.005 | 0.010 | Cerebellum L, Cerebellum R |
| Model 8 | MNC | 8804 | 0.011 | 0.030 | Calcarine L, Precentral L |
| Model 8 | BiDirect | 7382 | 0.019 | 0.051 | Paracentral Lobule R, Temporal Sup L |
| Model 9 | MACS | 0 | - | - | - |
| Model 9 | MNC | 2282 | 0.011 | 0.024 | Thalamus L, Frontal Mid 2 R |
| Model 9 | BiDirect | 8756 | 0.019 | 0.051 | Cerebellum L, Parietal Inf L |
| Model 10 | MACS | 2770 | 0.005 | 0.012 | Temporal Sup R, Cerebellum 4 5 L |
| Model 10 | MNC | 3216 | 0.011 | 0.023 | Cingulate Mid L, Rectus R |
| Model 10 | BiDirect | 4491 | 0.019 | 0.047 | Cerebellum L, OFC Post R |
| Model 11 | MACS | 50 | 0.005 | 0.006 | Temporal Pole Sup R, Precentral R |
| Model 11 | MNC | 4802 | 0.011 | 0.026 | Occipital Inf R, Precuneus L |
| Model 11 | BiDirect | 15863 | 0.019 | 0.060 | Temporal Sup R, OFC Post R |
| Model 12 | MACS | 5224 | 0.008 | 0.022 | Cerebellum L, Temporal Mid L |
| Model 12 | MNC | 11483 | 0.019 | 0.051 | Cerebellum L, Parietal Sup L |
| Model 12 | BiDirect | 6591 | 0.031 | 0.084 | Cerebellum L, Temporal Inf L |
| Model 13 | MACS | 25060 | 0.008 | 0.031 | Cerebellum L, Temporal Sup R |
| Model 13 | MNC | 5853 | 0.019 | 0.047 | Precentral L, Parietal Sup L |
| Model 13 | BiDirect | 6656 | 0.031 | 0.081 | Cerebellum L, Fusiform R |
| Model 14 | MACS | 4409 | 0.015 | 0.035 | Cerebellum R, Temporal Mid L |
| Model 14 | MNC | 13393 | 0.026 | 0.066 | Cerebellum L, Temporal Sup R |
| Model 14 | BiDirect | 7063 | 0.064 | 0.199 | Temporal Inf/Mid R, Temporal Mid/Inf L |
| Model 15 | MACS | 2650 | 0.019 | 0.036 | Cerebellum R, Cerebellum L |

| | | | | | |
|---|---|---|---|---|---|
| Model 15 | MNC | 6336 | 0.064 | 0.162 | Postcentral R, Occipital Mid R |
| Model 15 | BiDirect | 4354 | 0.060 | 0.149 | Cerebellum L, OFC Post R |

*Note*. Number of significant voxels at uncorrected p<.001 are shown, as well as their minimum and maximum effect size for each analysis. Cluster labelling was conducted based on the aal atlas using the atlasreader python package (Notter et al., 2019). Main regions of largest clusters are reported. MACS, Marburg Münster Affective Disorders Cohort Study; MNC, Münster Neuroimaging Cohort; BiDirect, BiDirect study cohort.

**Table S13**

*Results summary of single cohorts at a cohort-wise significance level of* $\boldsymbol{p_{unc}}$***<.01 – sex-stratified for male subsample***

| Model | Cohort | k significant | Partial $R^2$ | | Main regions |
|---|---|---|---|---|---|
| | | | Min | Max | |
| Model 1 | MACS | 707 | 0.009 | 0.017 | Postcentral R, Paracentral Lobule R |
| Model 1 | MNC | 1452 | 0.013 | 0.033 | Hippocampus L, Supramarginal R |
| Model 1 | BiDirect | 8774 | 0.021 | 0.061 | Cerebellum R, Fusiform R |
| Model 2 | MACS | 5785 | 0.009 | 0.022 | Paracentral Lobule R, Temporal Mid R |
| Model 2 | MNC | 3009 | 0.013 | 0.047 | Hippocampus L, Thalamus L |
| Model 2 | BiDirect | 10655 | 0.021 | 0.065 | Cerebellum R, Fusiform R |
| Model 3 | MACS | 2899 | 0.016 | 0.034 | Occipital Mid L, Precentral R |
| Model 3 | MNC | 1445 | 0.019 | 0.041 | Cerebellum R, Postcentral R |
| Model 3 | BiDirect | 2678 | 0.032 | 0.075 | Postcentral R, Cerebellum R |
| Model 4 | MACS | 131 | 0.019 | 0.026 | Supp Motor Area R, Temporal Mid R |
| Model 4 | MNC | 1190 | 0.045 | 0.073 | Frontal Sup Med L, Supramarginal R |
| Model 4 | BiDirect | 12785 | 0.059 | 0.177 | Occipital Sup R, Parietal Inf L |
| Model 5 | MACS | 792 | 0.009 | 0.021 | Paracentral Lobule R, Cingulate Mid R |
| Model 5 | MNC | 2591 | 0.013 | 0.028 | Hippocampus L, Cerebellum L |
| Model 5 | BiDirect | 14755 | 0.021 | 0.074 | Cerebellum R, Postcentral L |
| Model 6 | MACS | 641 | 0.009 | 0.023 | Postcentral L, Frontal Sup 2 L |
| Model 6 | MNC | 811 | 0.013 | 0.029 | OFC Med L, Frontal Sup 2 R |
| Model 6 | BiDirect | 4054 | 0.021 | 0.050 | Postcentral R, Cerebellum R |
| Model 7 | MACS | 2591 | 0.009 | 0.019 | Temporal Mid R, Postcentral R |
| Model 7 | MNC | 676 | 0.013 | 0.034 | Cerebellum L, Hippocampus L |
| Model 7 | BiDirect | 13369 | 0.021 | 0.064 | Postcentral L, Precuneus R |
| Model 8 | MACS | 929 | 0.009 | 0.024 | Paracentral Lobule R, Supp Motor Area R |
| Model 8 | MNC | 2202 | 0.013 | 0.028 | Hippocampus L, Thalamus L |
| Model 8 | BiDirect | 14045 | 0.021 | 0.069 | Postcentral L, Cerebellum L |
| Model 9 | MACS | 54 | 0.009 | 0.012 | Paracentral Lobule R, Frontal Sup 2 L |
| Model 9 | MNC | 19142 | 0.013 | 0.037 | Lingual L, Supramarginal R |
| Model 9 | BiDirect | 289 | 0.021 | 0.036 | Frontal Mid 2 R, Frontal Mid 2 L |
| Model 10 | MACS | 1063 | 0.009 | 0.022 | Cerebellum L, Fusiform R |
| Model 10 | MNC | 721 | 0.013 | 0.025 | Lingual L, OFC Med L |
| Model 10 | BiDirect | 3344 | 0.021 | 0.038 | Cerebellum R, Precuneus R |
| Model 11 | MACS | 2258 | 0.009 | 0.017 | Paracentral Lobule R, Occipital Mid L |
| Model 11 | MNC | 573 | 0.013 | 0.022 | OFC Med L, Hippocampus L |
| Model 11 | BiDirect | 7053 | 0.021 | 0.066 | Cerebellum R, Cerebellum L |
| Model 12 | MACS | 26 | 0.016 | 0.019 | Supramarginal R, Paracentral Lobule R |
| Model 12 | MNC | 3261 | 0.025 | 0.050 | Frontal Sup 2 R, Lingual L |
| Model 12 | BiDirect | 4882 | 0.041 | 0.112 | Occipital Sup R, Frontal Inf Tri R |
| Model 13 | MACS | 2624 | 0.016 | 0.030 | Fusiform L, Paracentral Lobule R |
| Model 13 | MNC | 6234 | 0.024 | 0.082 | Rectus L, Hippocampus L |
| Model 13 | BiDirect | 7067 | 0.040 | 0.102 | Frontal Inf Tri R, Lingual R |
| Model 14 | MACS | 1203 | 0.029 | 0.070 | Fusiform R, Lingual L |
| Model 14 | MNC | 2577 | 0.034 | 0.064 | Cerebellum R, Thalamus R |
| Model 14 | BiDirect | 2641 | 0.056 | 0.119 | Postcentral R, Occipital Inf R |
| Model 15 | MACS | 43 | 0.036 | 0.047 | Supp Motor Area R, Postcentral L |

| | | | | | |
|---|---|---|---|---|---|
| Model 15 | MNC | 7320 | 0.086 | 0.240 | Rectus L, Frontal Sup 2 R |
| Model 15 | BiDirect | 4788 | 0.145 | 0.336 | Occipital Sup R, Cuneus L |

*Note*. Number of significant voxels at uncorrected p<.001 are shown, as well as their minimum and maximum effect size for each analysis. Cluster labelling was conducted based on the aal atlas using the atlasreader python package (Notter et al., 2019). Main regions of largest clusters are reported. MACS, Marburg Münster Affective Disorders Cohort Study; MNC, Münster Neuroimaging Cohort; BiDirect, BiDirect study cohort.

**Table S14**

*Replicability across cohorts indicated by spatial overlap in significance at a cohort-wise level of* ***$p_{unc}$<.001 – sex-stratified for female subsample***

| | MACS-MNC | | | | MACS-BiDirect | | | | MNC-BiDirect | | | | MACS-MNC-BiDirect | | | |
|---|---|---|---|---|---|---|---|---|---|---|---|---|---|---|---|---|
| Model | k overlap | p-value | p-value (FDR) | DICE | k overlap | p-value | p-value (FDR) | DICE | k overlap | p-value | p-value (FDR) | DICE | k overlap | p-value | p-value (FDR) | DICE |
| Model 1 | 0 | 1 | 1 | 0 | 0 | 1 | 1 | 0 | 0 | 1 | 1 | 0 | 0 | 1 | 1 | - |
| Model 2 | 0 | 1 | 1 | 0 | 0 | 1 | 1 | 0 | 0 | 1 | 1 | 0 | 0 | 1 | 1 | - |
| Model 3 | 0 | 1 | 1 | 0 | 0 | 1 | 1 | 0 | 0 | 1 | 1 | 0 | 0 | 1 | 1 | - |
| Model 4 | 0 | 1 | 1 | 0 | 0 | 1 | 1 | 0 | 0 | 1 | 1 | 0 | 0 | 1 | 1 | - |
| Model 5 | 0 | 1 | 1 | 0 | 0 | 1 | 1 | 0 | 0 | 1 | 1 | 0 | 0 | 1 | 1 | - |
| Model 6 | 0 | 1 | 1 | 0 | 0 | 1 | 1 | 0 | 0 | 1 | 1 | 0 | 0 | 1 | 1 | - |
| Model 7 | 0 | 1 | 1 | 0 | 0 | 1 | 1 | 0 | 0 | 1 | 1 | 0 | 0 | 1 | 1 | - |
| Model 8 | 0 | 1 | 1 | 0 | 0 | 1 | 1 | 0 | 0 | 1 | 1 | 0 | 0 | 1 | 1 | - |
| Model 9 | 0 | 1 | 1 | 0 | 0 | 1 | 1 | 0 | 0 | 1 | 1 | 0 | 0 | 1 | 1 | - |
| Model 10 | 0 | 1 | 1 | 0 | 0 | 1 | 1 | 0 | 0 | 1 | 1 | 0 | 0 | 1 | 1 | - |
| Model 11 | 0 | 1 | 1 | 0 | 0 | 1 | 1 | 0 | 0 | 1 | 1 | 0 | 0 | 1 | 1 | - |
| Model 12 | 0 | 1 | 1 | 0 | 0 | 1 | 1 | 0 | 0 | 1 | 1 | 0 | 0 | 1 | 1 | - |
| Model 13 | 0 | 1 | 1 | 0 | 0 | 1 | 1 | 0 | 0 | 1 | 1 | 0 | 0 | 1 | 1 | - |
| Model 14 | 0 | 1 | 1 | 0 | 4 | .011 | .659 | 0.006 | 0 | 1 | 1 | 0 | 0 | 1 | 1 | - |
| Model 15 | 0 | 1 | 1 | 0 | 0 | 1 | 1 | 0 | 0 | 1 | 1 | 0 | 0 | 1 | 1 | - |

*Note*. The overlap in significant voxels is presented across all statistical models and all cohort combinations. The DICE score is presented for pairwise combinations with any voxels overlapping. P-values are based on permutation-based null distribution of cohort-wise overlap. FDR-correction was applied using the Benjamini-Hochberg procedure (Benjamini & Hochberg, 1995). MACS, Marburg Münster Affective Disorders Cohort Study; MNC, Münster Neuroimaging Cohort; BiDirect, BiDirect study cohort.

**Table S15**

*Replicability across cohorts indicated by spatial overlap in significance at a cohort-wise level of* ***$p_{unc}$<.001 – sex-stratified for male subsample***

| | MACS-MNC | | | | MACS-BiDirect | | | | MNC-BiDirect | | | | MACS-MNC-BiDirect | | | |
|---|---|---|---|---|---|---|---|---|---|---|---|---|---|---|---|---|
| Model | k overlap | p-value | p-value (FDR) | DICE | k overlap | p-value | p-value (FDR) | DICE | k overlap | p-value | p-value (FDR) | DICE | k overlap | p-value | p-value (FDR) | DICE |
| Model 1 | 0 | 1 | 1 | 0 | 0 | 1 | 1 | 0 | 0 | 1 | 1 | 0 | 0 | 1 | 1 | - |
| Model 2 | 0 | 1 | 1 | 0 | 0 | 1 | 1 | 0 | 0 | 1 | 1 | 0 | 0 | 1 | 1 | - |
| Model 3 | 0 | 1 | 1 | 0 | 0 | 1 | 1 | 0 | 0 | 1 | 1 | 0 | 0 | 1 | 1 | - |
| Model 4 | 0 | 1 | 1 | 0 | 0 | 1 | 1 | 0 | 0 | 1 | 1 | 0 | 0 | 1 | 1 | - |
| Model 5 | 0 | 1 | 1 | 0 | 0 | 1 | 1 | 0 | 0 | 1 | 1 | 0 | 0 | 1 | 1 | - |
| Model 6 | 0 | 1 | 1 | 0 | 0 | 1 | 1 | 0 | 0 | 1 | 1 | 0 | 0 | 1 | 1 | - |
| Model 7 | 0 | 1 | 1 | 0 | 0 | 1 | 1 | 0 | 0 | 1 | 1 | 0 | 0 | 1 | 1 | - |
| Model 8 | 0 | 1 | 1 | 0 | 0 | 1 | 1 | 0 | 0 | 1 | 1 | 0 | 0 | 1 | 1 | - |
| Model 9 | 0 | 1 | 1 | 0 | 0 | 1 | 1 | 0 | 0 | 1 | 1 | 0 | 0 | 1 | 1 | - |
| Model 10 | 0 | 1 | 1 | 0 | 0 | 1 | 1 | 0 | 0 | 1 | 1 | 0 | 0 | 1 | 1 | - |
| Model 11 | 0 | 1 | 1 | 0 | 0 | 1 | 1 | 0 | 0 | 1 | 1 | 0 | 0 | 1 | 1 | - |
| Model 12 | 0 | 1 | 1 | 0 | 0 | 1 | 1 | 0 | 0 | 1 | 1 | 0 | 0 | 1 | 1 | - |
| Model 13 | 0 | 1 | 1 | 0 | 0 | 1 | 1 | 0 | 0 | 1 | 1 | 0 | 0 | 1 | 1 | - |
| Model 14 | 0 | 1 | 1 | 0 | 0 | 1 | 1 | 0 | 0 | 1 | 1 | 0 | 0 | 1 | 1 | - |
| Model 15 | 0 | 1 | 1 | 0 | 0 | 1 | 1 | 0 | 0 | 1 | 1 | 0 | 0 | 1 | 1 | - |

*Note*. The overlap in significant voxels is presented across all statistical models and all cohort combinations. The DICE score is presented for pairwise combinations with any voxels overlapping. P-values are based on permutation-based null distribution of cohort-wise overlap. FDR-correction was applied using the Benjamini-Hochberg procedure (Benjamini & Hochberg, 1995). MACS, Marburg Münster Affective Disorders Cohort Study; MNC, Münster Neuroimaging Cohort; BiDirect, BiDirect study cohort.

**Table S16**

*Replicability across cohorts indicated by spatial overlap in significance at a cohort-wise level of* $\boldsymbol{p_{unc}<.01}$

| | MACS-MNC | | | | MACS-BiDirect | | | | MNC-BiDirect | | | | MACS-MNC-BiDirect | | | |
|---|---|---|---|---|---|---|---|---|---|---|---|---|---|---|---|---|
| Model | k overlap | p-value | p-value (FDR) | DICE | k overlap | p-value | p-value (FDR) | DICE | k overlap | p-value | p-value (FDR) | DICE | k overlap | p-value | p-value (FDR) | DICE |
| Model 1 | 0 | 1 | 1 | 0 | 0 | 1 | 1 | 0 | 6 | 0.380 | 0.911 | 0.001 | 0 | 1 | 1 | - |
| Model 2 | 73 | 0.132 | 0.609 | 0.011 | 329 | 0.038 | 0.228 | 0.024 | 223 | 0.032 | 0.228 | 0.022 | 4 | 0.010 | 0.150 | - |
| Model 3 | 8 | 0.318 | 0.794 | 0.002 | 11 | 0.308 | 0.794 | 0.006 | 12 | 0.303 | 0.794 | 0.003 | 0 | 1 | 1 | - |
| Model 4 | 0 | 1 | 1 | 0 | 0 | 1 | 1 | 0 | 0 | 1 | 1 | 0 | 0 | 1 | 1 | - |
| Model 5 | 0 | 1 | 1 | 0 | 190 | 0.065 | 0.325 | 0.046 | 14 | 0.295 | 0.794 | 0.003 | 0 | 1 | 1 | - |
| Model 6 | 0 | 1 | 1 | 0 | 0 | 1 | 1 | 0 | 18 | 0.292 | 0.794 | 0.003 | 0 | 1 | 1 | - |
| Model 7 | 0 | 1 | 1 | 0 | 38 | 0.216 | 0.746 | 0.011 | 0 | 1 | 1 | 0 | 0 | 1 | 1 | - |
| Model 8 | 50 | 0.190 | 0.731 | 0.013 | 286 | 0.037 | 0.228 | 0.032 | 17 | 0.262 | 0.794 | 0.002 | 0 | 1 | 1 | - |
| Model 9 | 0 | 1 | 1 | 0 | 0 | 1 | 1 | 0 | 0 | 1 | 1 | 0 | 0 | 1 | 1 | - |
| Model 10 | 0 | 1 | 1 | 0 | 48 | 0.185 | 0.731 | 0.010 | 0 | 1 | 1 | 0 | 0 | 1 | 1 | - |
| Model 11 | 0 | 1 | 1 | 0 | 0 | 1 | 1 | 0 | 609 | 0.007 | 0.150 | 0.061 | 0 | 1 | 1 | - |
| Model 12 | 0 | 1 | 1 | 0 | 0 | 1 | 1 | 0 | 346 | 0.026 | 0.223 | 0.047 | 0 | 1 | 1 | - |
| Model 13 | 739 | 0.009 | 0.150 | 0.056 | 1329 | 0.004 | 0.150 | 0.081 | 472 | 0.019 | 0.190 | 0.042 | 12 | 0.013 | 0.156 | - |
| Model 14 | 2 | 0.436 | 1 | 0.000 | 0 | 1 | 1 | 0 | 127 | 0.059 | 0.321 | 0.020 | 0 | 1 | 1 | - |
| Model 15 | 36 | 0.224 | 0.746 | 0.007 | 0 | 1 | 1 | 0 | 46 | 0.195 | 0.731 | 0.006 | 0 | 1 | 1 | - |

*Note*. The overlap in significant voxels is presented across all statistical models and all cohort combinations. The DICE score is presented for pairwise combinations with any voxels overlapping. P-values are based on permutation-based null distribution of cohort-wise overlap. FDR-correction was applied using the Benjamini-Hochberg procedure (Benjamini & Hochberg, 1995). MACS, Marburg Münster Affective Disorders Cohort Study; MNC, Münster Neuroimaging Cohort; BiDirect, BiDirect study cohort.

**Table S17**

*Replicability across cohorts indicated by spatial overlap in significance at a cohort-wise level of* ***$p_{unc}$<.01 – sex-stratified for female subsample***

| | MACS-MNC | | | | MACS-BiDirect | | | | MNC-BiDirect | | | | MACS-MNC-BiDirect | | | |
|---|---|---|---|---|---|---|---|---|---|---|---|---|---|---|---|---|
| Model | k overlap | p-value | p-value (FDR) | DICE | k overlap | p-value | p-value (FDR) | DICE | k overlap | p-value | p-value (FDR) | DICE | k overlap | p-value | p-value (FDR) | DICE |
| Model 1 | 0 | 1 | 1 | 0 | 0 | 1 | 1 | 0 | 0 | 1 | 1 | 0 | 0 | 1 | 1 | - |
| Model 2 | 102 | 0.118 | 0.535 | 0.012 | 464 | 0.018 | 0.360 | 0.047 | 28 | 0.231 | 0.629 | 0.004 | 0 | 1 | 1 | - |
| Model 3 | 51 | 0.168 | 0.539 | 0.005 | 0 | 1 | 1 | 0 | 0 | 1 | 1 | 0 | 0 | 1 | 1 | - |
| Model 4 | 0 | 1 | 1 | 0 | 4 | 0.411 | 1 | 0.001 | 77 | 0.125 | 0.535 | 0.020 | 0 | 1 | 1 | - |
| Model 5 | 0 | 1 | 1 | 0 | 215 | 0.049 | 0.437 | 0.078 | 38 | 0.189 | 0.539 | 0.007 | 0 | 1 | 1 | - |
| Model 6 | 0 | 1 | 1 | 0 | 0 | 1 | 1 | 0 | 2 | 0.455 | 1 | 0.000 | 0 | 1 | 1 | - |
| Model 7 | 0 | 1 | 1 | 0 | 153 | 0.067 | 0.446 | 0.050 | 64 | 0.160 | 0.539 | 0.011 | 0 | 1 | 1 | - |
| Model 8 | 0 | 1 | 1 | 0 | 95 | 0.097 | 0.535 | 0.018 | 53 | 0.140 | 0.539 | 0.007 | 0 | 1 | 1 | - |
| Model 9 | 0 | 1 | 1 | 0 | 0 | 1 | 1 | 0 | 0 | 1 | 1 | 0 | 0 | 1 | 1 | - |
| Model 10 | 1 | 0.472 | 1 | 0.000 | 0 | 1 | 1 | 0 | 0 | 1 | 1 | 0 | 0 | 1 | 1 | - |
| Model 11 | 0 | 1 | 1 | 0 | 0 | 1 | 1 | 0 | 44 | 0.183 | 0.539 | 0.004 | 0 | 1 | 1 | - |
| Model 12 | 47 | 0.182 | 0.539 | 0.006 | 75 | 0.123 | 0.535 | 0.013 | 83 | 0.118 | 0.535 | 0.009 | 0 | 1 | 1 | - |
| Model 13 | 185 | 0.060 | 0.446 | 0.012 | 1655 | 0.002 | 0.120 | 0.104 | 0 | 1 | 1 | 0 | 0 | 1 | 1 | - |
| Model 14 | 239 | 0.032 | 0.437 | 0.027 | 430 | 0.014 | 0.360 | 0.075 | 172 | 0.051 | 0.437 | 0.017 | 0 | 1 | 1 | - |
| Model 15 | 226 | 0.040 | 0.437 | 0.050 | 6 | 0.411 | 1.000 | 0.002 | 51 | 0.179 | 0.539 | 0.010 | 0 | 1 | 1 | - |

*Note*. The overlap in significant voxels is presented across all statistical models and all cohort combinations. The DICE score is presented for pairwise combinations with any voxels overlapping. P-values are based on permutation-based null distribution of cohort-wise overlap. FDR-correction was applied using the Benjamini-Hochberg procedure (Benjamini & Hochberg, 1995). MACS, Marburg Münster Affective Disorders Cohort Study; MNC, Münster Neuroimaging Cohort; BiDirect, BiDirect study cohort.

### Table S18

*Replicability across cohorts indicated by spatial overlap in significance at a level of **$p_{unc}$<.01 – sex-stratified for male subsample***

| | MACS-MNC | | | | MACS-BiDirect | | | | MNC-BiDirect | | | | MACS-MNC-BiDirect | | | |
|---|---|---|---|---|---|---|---|---|---|---|---|---|---|---|---|---|
| Model | k overlap | p-value | p-value (FDR) | DICE | k overlap | p-value | p-value (FDR) | DICE | k overlap | p-value | p-value (FDR) | DICE | k overlap | p-value | p-value (FDR) | DICE |
| Model 1 | 0 | 1 | 1 | 0 | 0 | 1 | 1 | 0 | 3 | 0.462 | 1 | 0.001 | 0 | 1 | 1 | - |
| Model 2 | 93 | 0.102 | 1 | 0.021 | 144 | 0.090 | 1 | 0.018 | 47 | 0.195 | 1 | 0.007 | 0 | 1 | 1 | - |
| Model 3 | 0 | 1 | 1 | 0 | 0 | 1 | 1 | 0 | 0 | 1 | 1 | 0 | 0 | 1 | 1 | - |
| Model 4 | 0 | 1 | 1 | 0 | 0 | 1 | 1 | 0 | 4 | 0.418 | 1 | 0.001 | 0 | 1 | 1 | - |
| Model 5 | 0 | 1 | 1 | 0 | 0 | 1 | 1 | 0 | 13 | 0.333 | 1 | 0.001 | 0 | 1 | 1 | - |
| Model 6 | 0 | 1 | 1 | 0 | 0 | 1 | 1 | 0 | 9 | 0.372 | 1 | 0.004 | 0 | 1 | 1 | - |
| Model 7 | 0 | 1 | 1 | 0 | 12 | 0.340 | 1 | 0.002 | 0 | 1 | 1 | 0 | 0 | 1 | 1 | - |
| Model 8 | 0 | 1 | 1 | 0 | 3 | 0.436 | 1 | 0.000 | 128 | 0.089 | 1 | 0.016 | 0 | 1 | 1 | - |
| Model 9 | 0 | 1 | 1 | 0 | 0 | 1 | 1 | 0 | 12 | 0.249 | 1 | 0.001 | 0 | 1 | 1 | - |
| Model 10 | 0 | 1 | 1 | 0 | 0 | 1 | 1 | 0 | 0 | 1 | 1 | 0 | 0 | 1 | 1 | - |
| Model 11 | 0 | 1 | 1 | 0 | 25 | 0.266 | 1 | 0.005 | 0 | 1 | 1 | 0 | 0 | 1 | 1 | - |
| Model 12 | 0 | 1 | 1 | 0 | 0 | 1 | 1 | 0 | 76 | 0.123 | 1 | 0.019 | 0 | 1 | 1 | - |
| Model 13 | 64 | 0.153 | 1 | 0.014 | 62 | 0.159 | 1 | 0.013 | 90 | 0.109 | 1 | 0.014 | 0 | 1 | 1 | - |
| Model 14 | 0 | 1 | 1 | 0 | 0 | 1 | 1 | 0 | 0 | 1 | 1 | 0 | 0 | 1 | 1 | - |
| Model 15 | 0 | 1 | 1 | 0 | 0 | 1 | 1 | 0 | 40 | 0.239 | 1 | 0.007 | 0 | 1 | 1 | - |

*Note*. The overlap in significant voxels is presented across all statistical models and all cohort combinations. The DICE score is presented for pairwise combinations with any voxels overlapping. P-values for each observed cohort-combination overlap are based on permutation-based null distribution. FDR-correction was applied using the Benjamini-Hochberg procedure (Benjamini & Hochberg, 1995). MACS, Marburg Münster Affective Disorders Cohort Study; MNC, Münster Neuroimaging Cohort; BiDirect, BiDirect study cohort.

**Figure S1**

*Age distribution across cohorts and diagnosis groups*

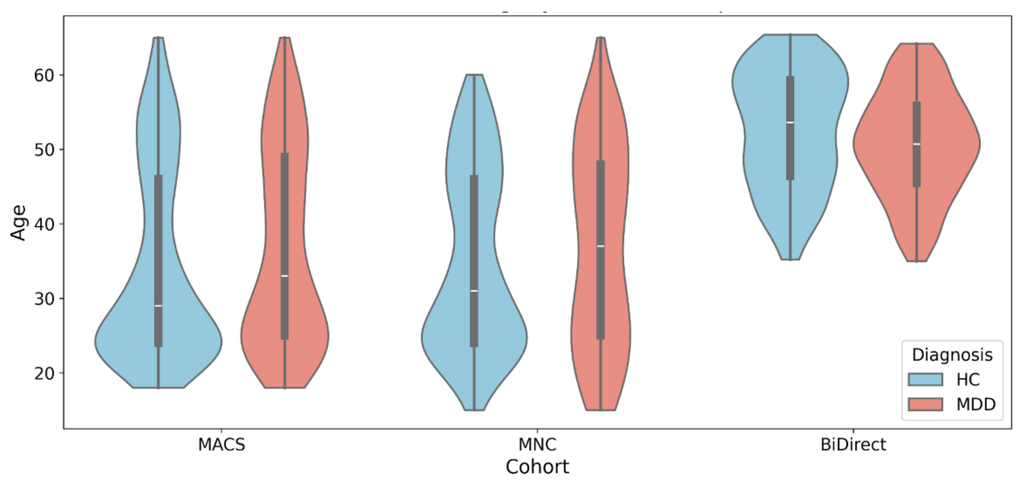


*Note.* HC, healthy controls; MDD, major depressive disorder.

**Figure S2**

*Association between CTQ scales, demographic variables and clinical variables within the **MACS** cohort*

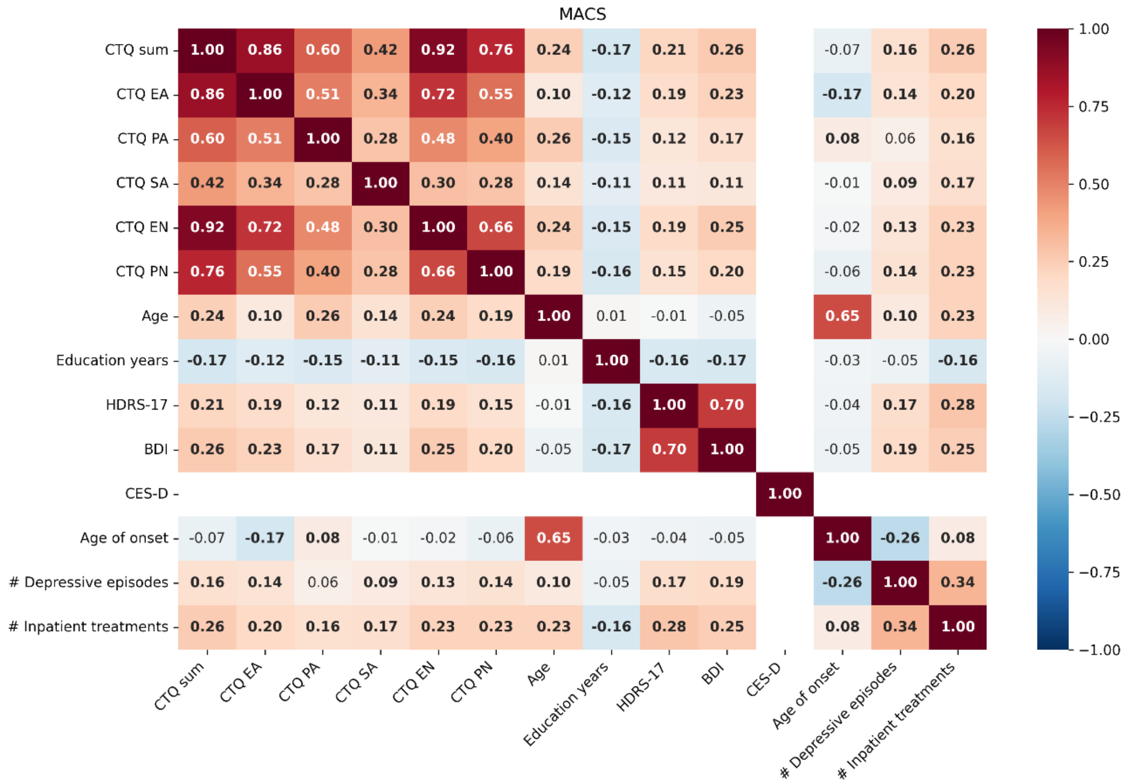


*Note.* Spearman correlations are shown. All correlations involving clinical variables (HDRS-17, BDI, Age of onset, number of depressive episodes and number of inpatient treatments) were only calculated within the MDD subsample. The BDI was only available within MACS and MNC, while the CES-D was only available for the BiDirect cohort. Significant associations at p<.05 are shown in bold font. CTQ, childhood trauma questionnaire; EA, emotional abuse; PA, physical abuse; SA, sexual abuse; EN, emotional neglect; PN, physical neglect; HDRS-17, 17-item Hamilton Depression Rating Scale; BDI, Beck Depression Inventory; CES-D, Center for Epidemiologic Studies Depression Scale.

### Figure S3

*Association between CTQ scales, demographic variables and clinical variables within the **MNC** cohort*

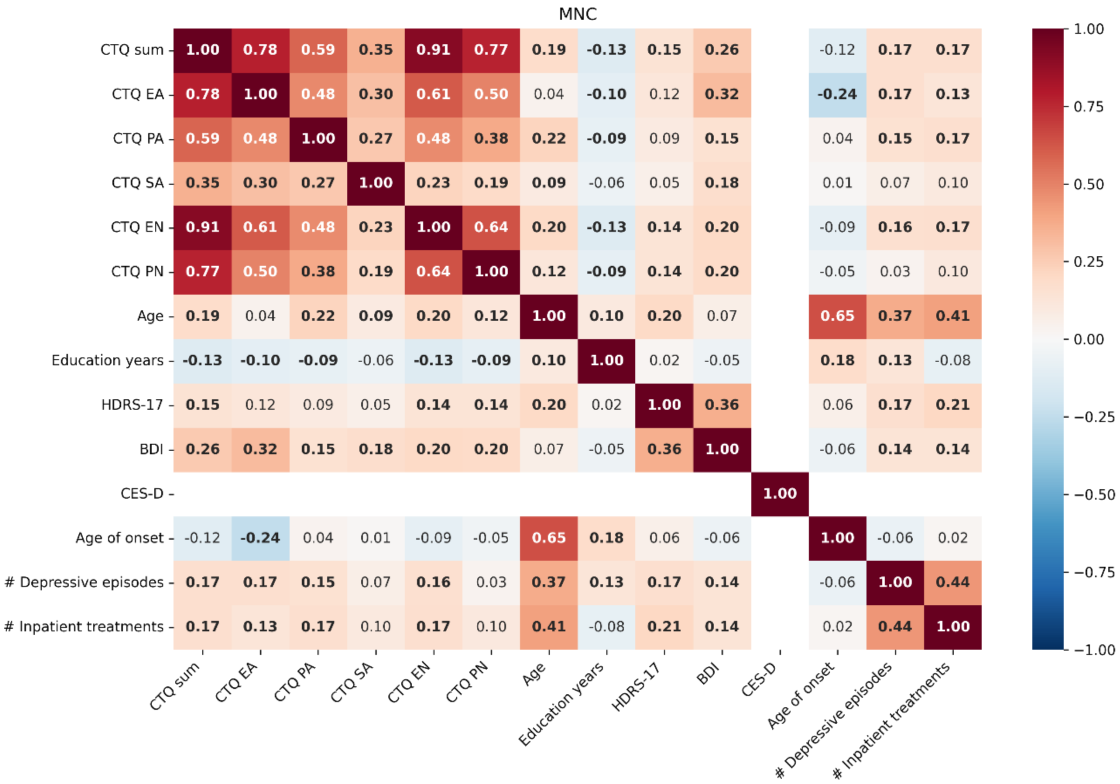


*Note.* Spearman correlations are shown. All correlations involving clinical variables (HDRS-17, BDI, Age of onset, number of depressive episodes and number of inpatient treatments) were only calculated within the MDD subsample. The BDI was only available within MACS and MNC, while the CES-D was only available for the BiDirect cohort. Significant associations at p<.05 are shown in bold font. CTQ, childhood trauma questionnaire; EA, emotional abuse; PA, physical abuse; SA, sexual abuse; EN, emotional neglect; PN, physical neglect; HDRS-17, 17-item Hamilton Depression Rating Scale; BDI, Beck Depression Inventory; CES-D, Center for Epidemiologic Studies Depression Scale.

### Figure S4

*Association between CTQ scales, demographic variables and clinical variables within the* ***BiDirect*** *cohort*

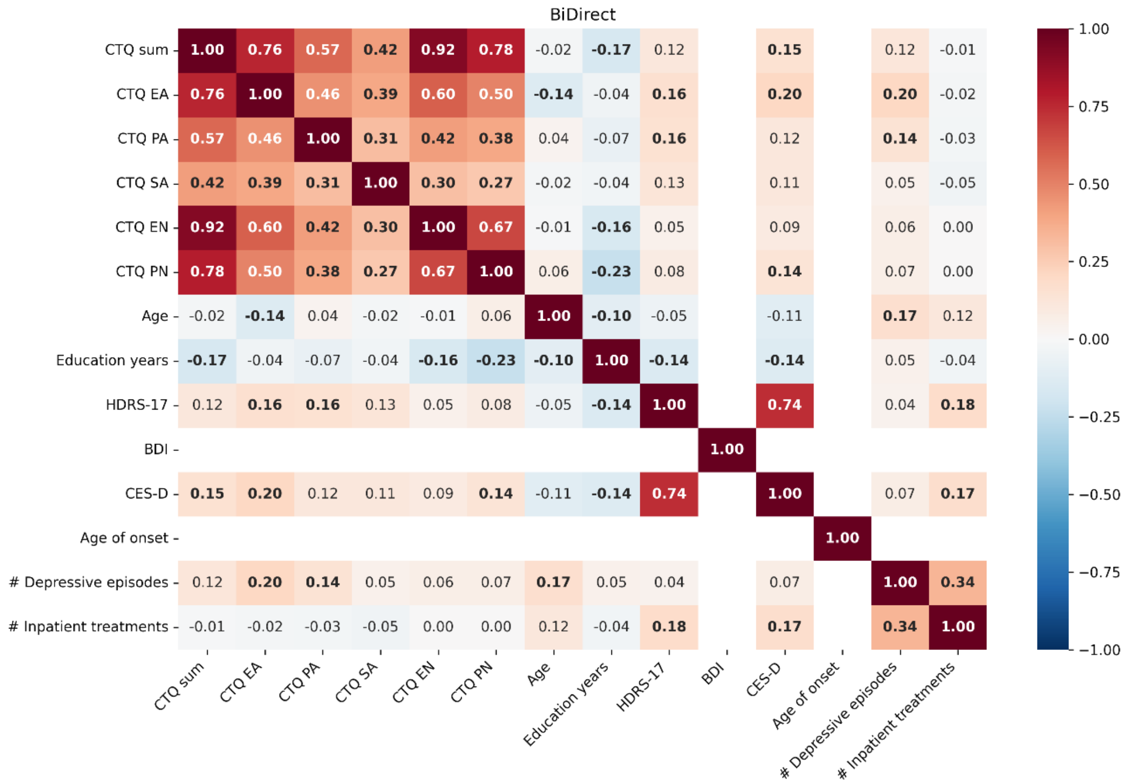


*Note.* Spearman correlations are shown. All correlations involving clinical variables (HDRS-17, BDI, Age of onset, number of depressive episodes and number of inpatient treatments) were only calculated within the MDD subsample. The BDI was only available within MACS and MNC, while the CES-D was only available for the BiDirect cohort. Significant associations at $p<.05$ are shown in bold font. CTQ, childhood trauma questionnaire; EA, emotional abuse; PA, physical abuse; SA, sexual abuse; EN, emotional neglect; PN, physical neglect; HDRS-17, 17-item Hamilton Depression Rating Scale; BDI, Beck Depression Inventory; CES-D, Center for Epidemiologic Studies Depression Scale.

**Figure S5**

*Distributions of CTQ scales across HC and MDD groups stratified by cohorts*

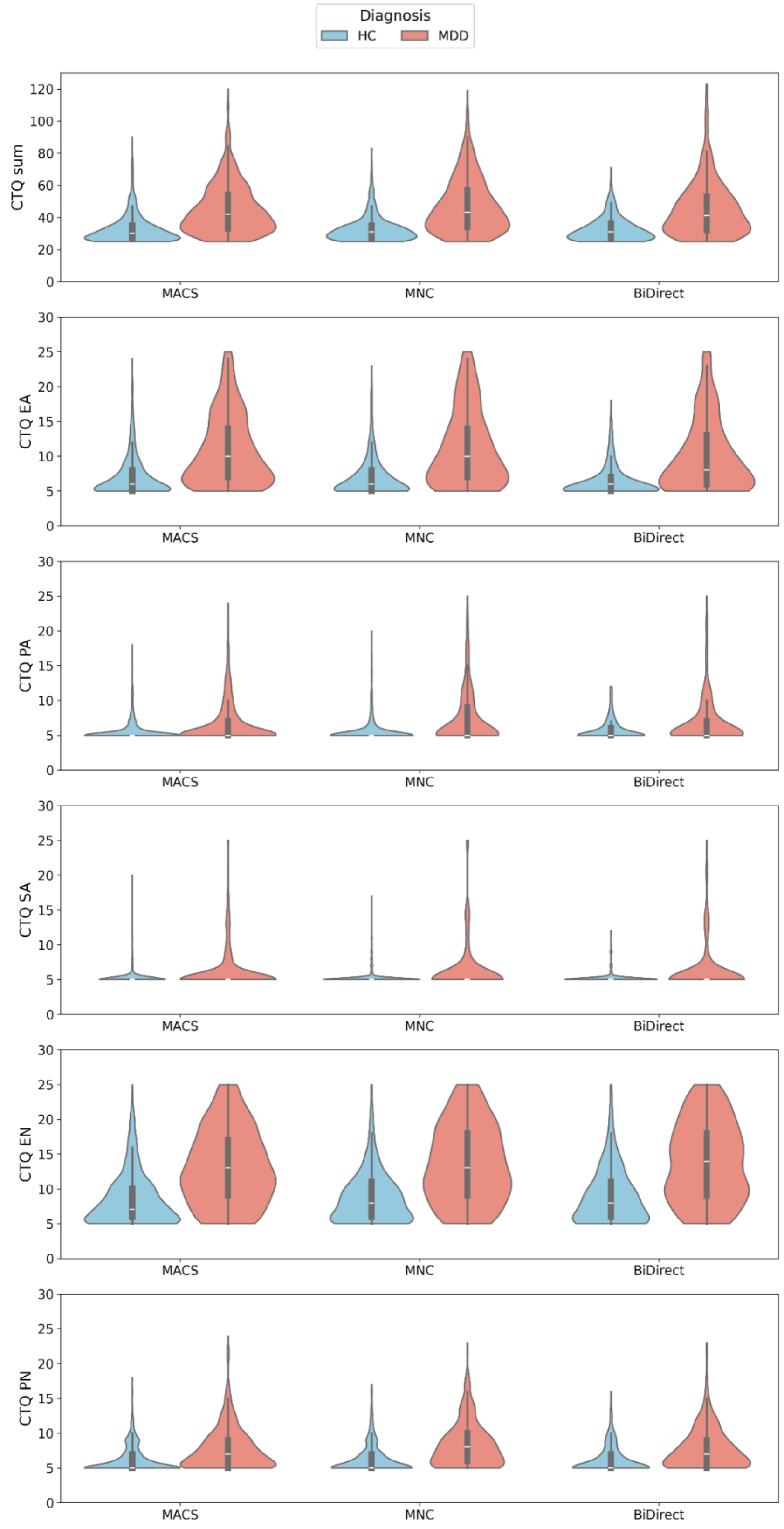


*Note.* Violin plots are shown depicting the distribution of the CTQ sum scale, as well as the five CTQ subscales. HC, healthy controls; MDD, major depressive disorder; CTQ, childhood trauma questionnaire; EA, emotional abuse; PA, physical abuse; SA, sexual abuse; EN, emotional neglect; PN, physical neglect.

**Figure S6**

*Significant clusters across cohort-wise analyses (replicability analysis) for* ***model 1*** *– CTQ sum in HC and MDD samples controlling for MDD diagnosis*

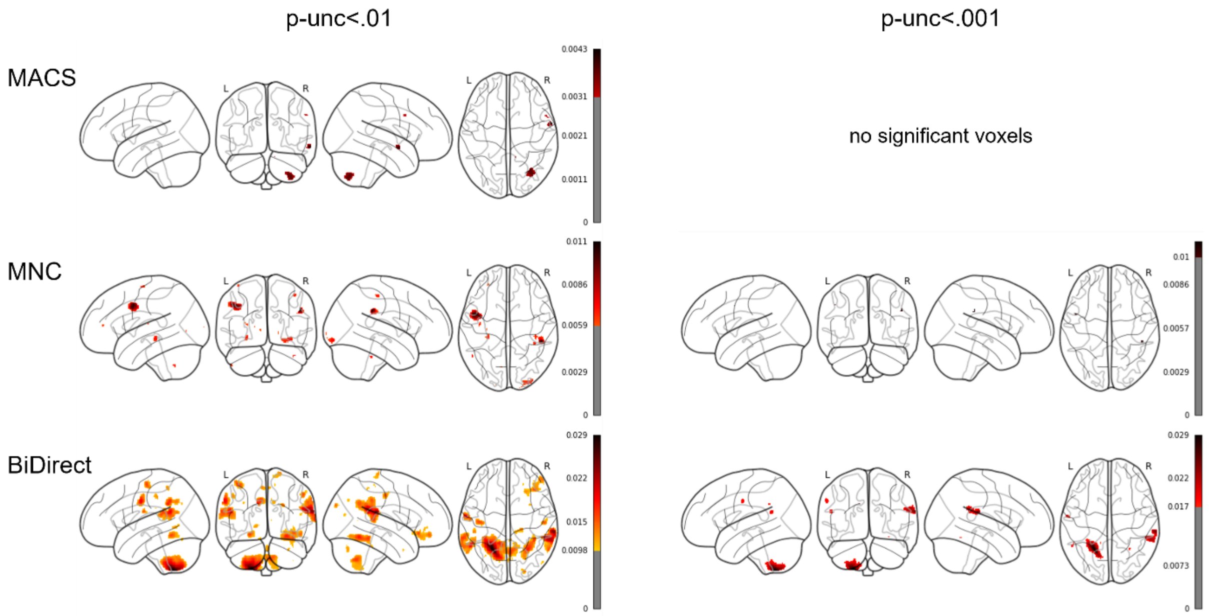


*Note.* Glass brains are shown with maximum intensity projections. Color bars represent the partial $R^2$ of the maltreatment predictor. Results are shown thresholded using the uncorrected thresholds $p_{unc}$<.01 (left column) and $p_{unc}$<.001 (right column).

**Figure S7**

*Significant clusters across cohort-wise analyses (replicability analysis) for* ***model 2*** *– CTQ sum as a predictor in HC and MDD samples without controlling for MDD diagnosis*

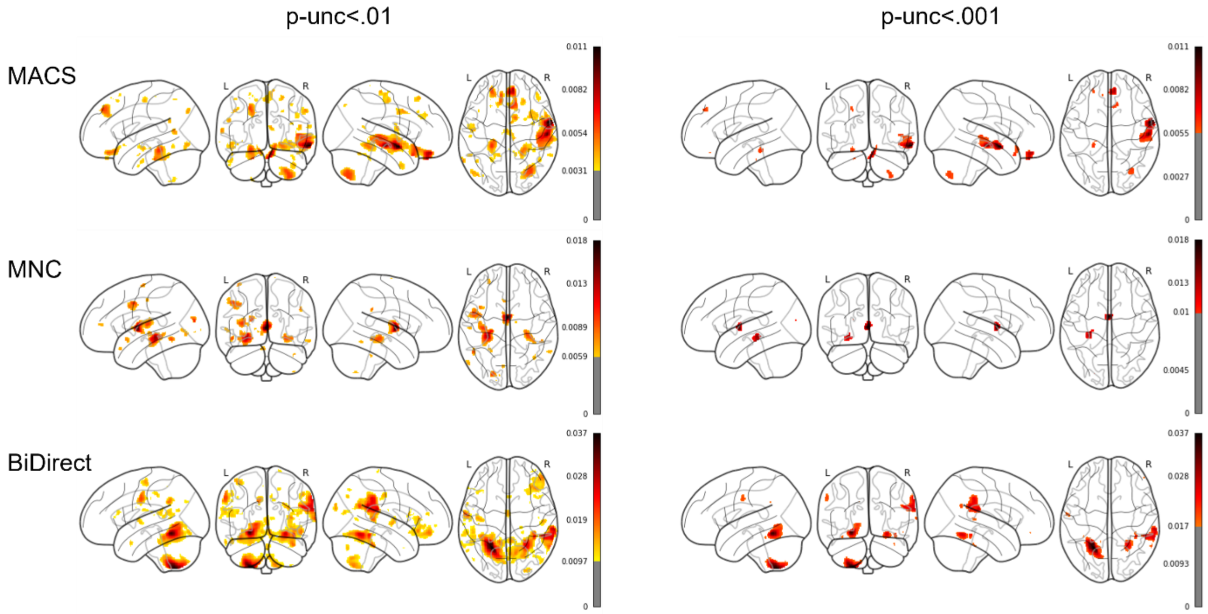


*Note.* Glass brains are shown with maximum intensity projections. Color bars represent the partial $R^2$ of the maltreatment predictor. Results are shown thresholded using the uncorrected thresholds $p_{unc}<.01$ (left column) and $p_{unc}<.001$ (right column).

**Figure S8**

*Significant clusters across cohort-wise analyses (replicability analysis) for* ***model 3*** *– CTQ sum as a predictor in HC subsamples*

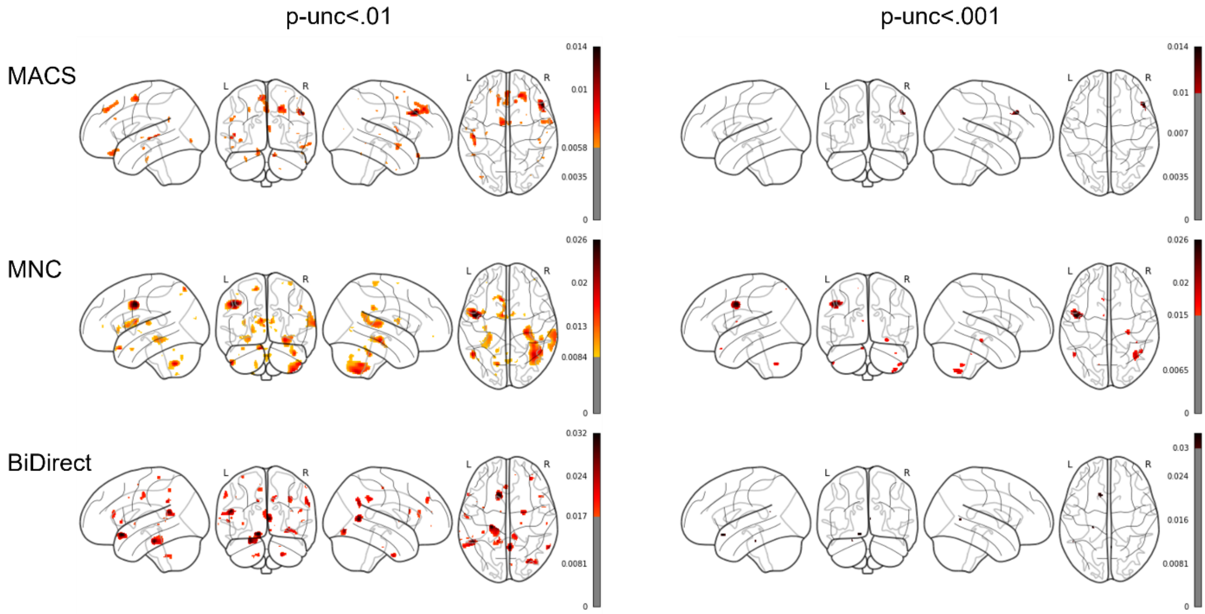


*Note.* Glass brains are shown with maximum intensity projections. Color bars represent the partial $R^2$ of the maltreatment predictor. Results are shown thresholded using the uncorrected thresholds $p_{unc}$<.01 (left column) and $p_{unc}$<.001 (right column).

**Figure S9**

*Significant clusters across cohort-wise analyses (replicability analysis) for* ***model 4*** *– CTQ sum as a predictor in MDD subsamples*

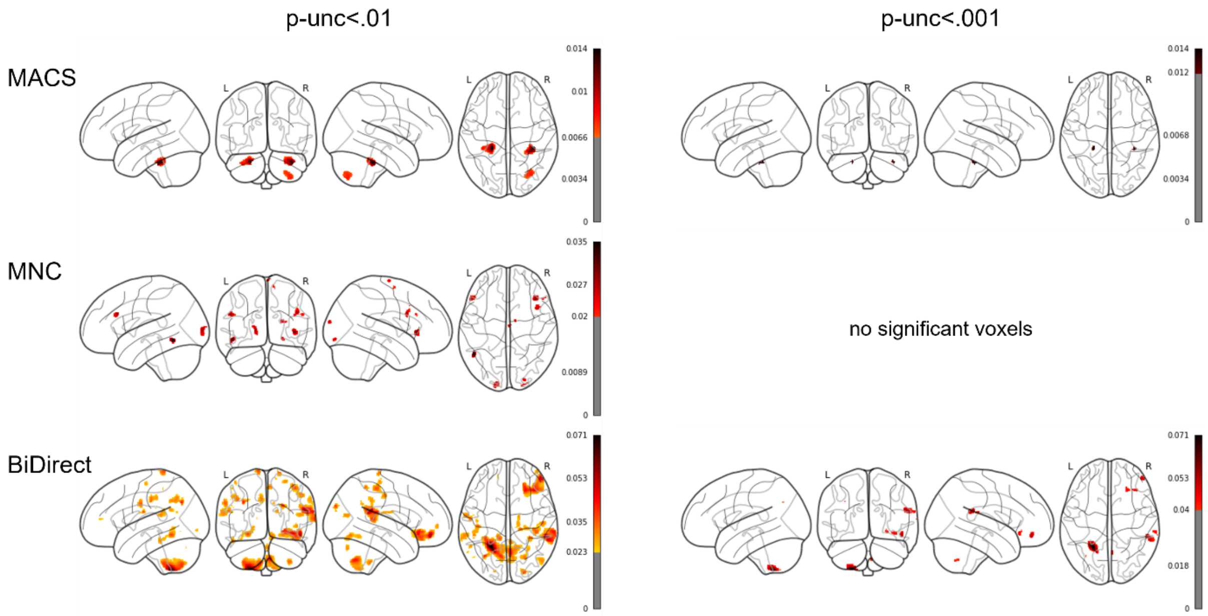


*Note.* Glass brains are shown with maximum intensity projections. Color bars represent the partial $R^2$ of the maltreatment predictor. Results are shown thresholded using the uncorrected thresholds $p_{unc}<.01$ (left column) and $p_{unc}<.001$ (right column).

**Figure S10**

*Significant clusters across cohort-wise analyses (replicability analysis) for* ***model 5*** *– CTQ abuse subscales as a predictor in HC and MDD samples controlling for MDD diagnosis*

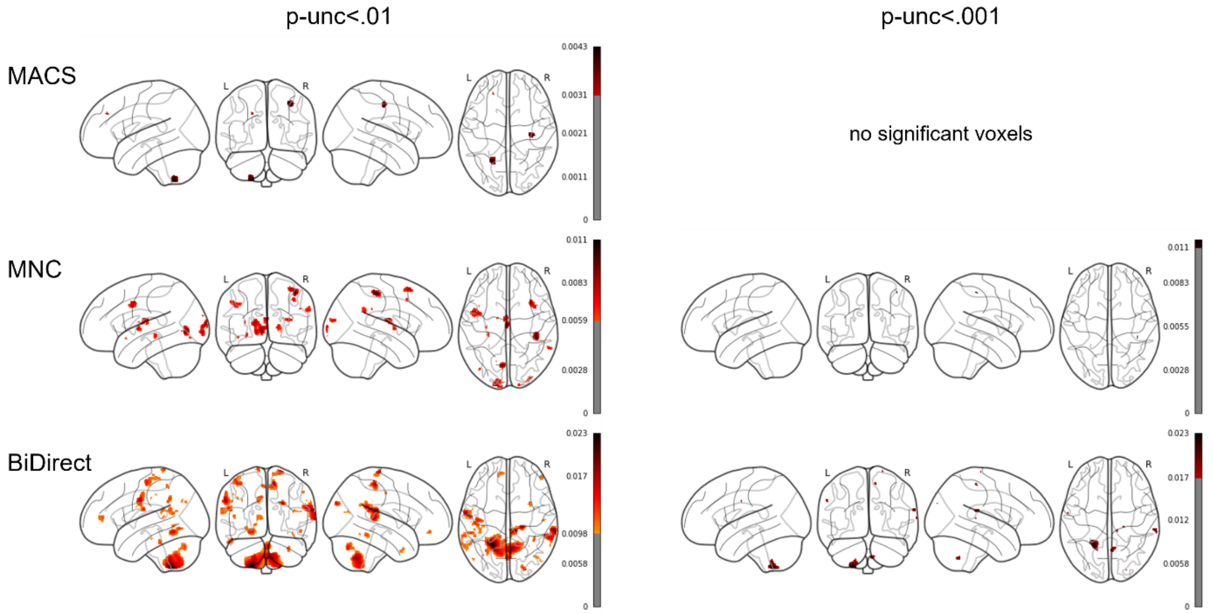


*Note.* Glass brains are shown with maximum intensity projections. Color bars represent the partial $R^2$ of the maltreatment predictor. Results are shown thresholded using the uncorrected thresholds $p_{unc}$<.01 (left column) and $p_{unc}$<.001 (right column).

**Figure S11**

*Significant clusters across cohort-wise analyses (replicability analysis) for **model 6** – CTQ neglect subscales as a predictor in HC and MDD samples controlling for MDD diagnosis*

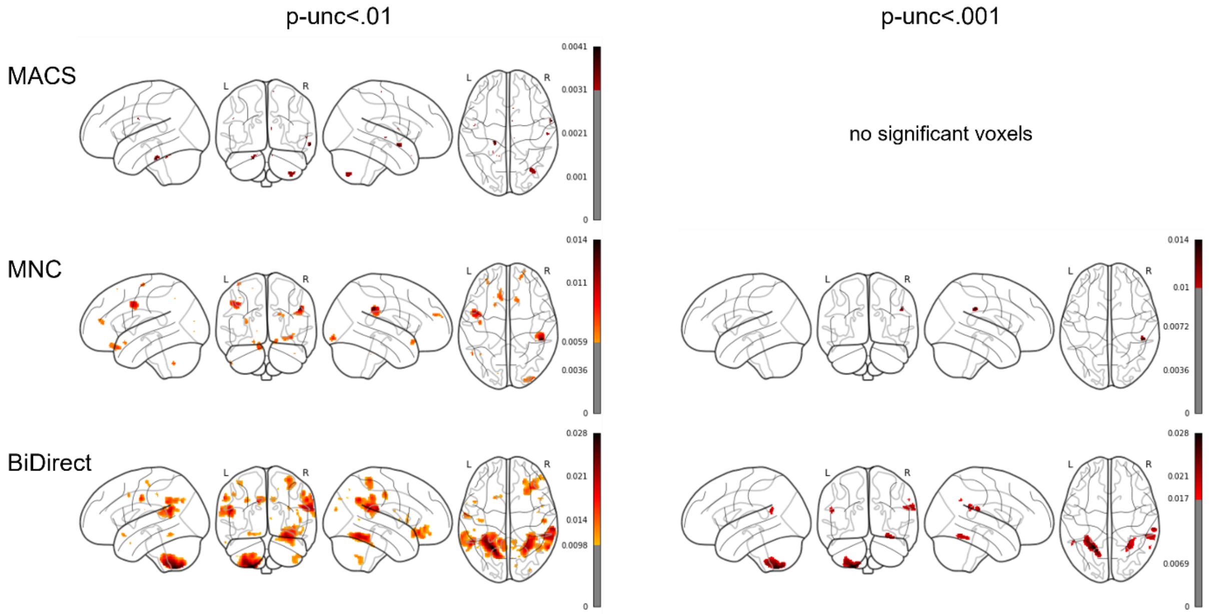


*Note.* Glass brains are shown with maximum intensity projections. Color bars represent the partial $R^2$ of the maltreatment predictor. Results are shown thresholded using the uncorrected thresholds $p_{unc}<.01$ (left column) and $p_{unc}<.001$ (right column).

**Figure S12**

*Significant clusters across cohort-wise analyses (replicability analysis) for **model 7** – CTQ subscale emotional abuse as a predictor in HC and MDD samples controlling for MDD diagnosis*

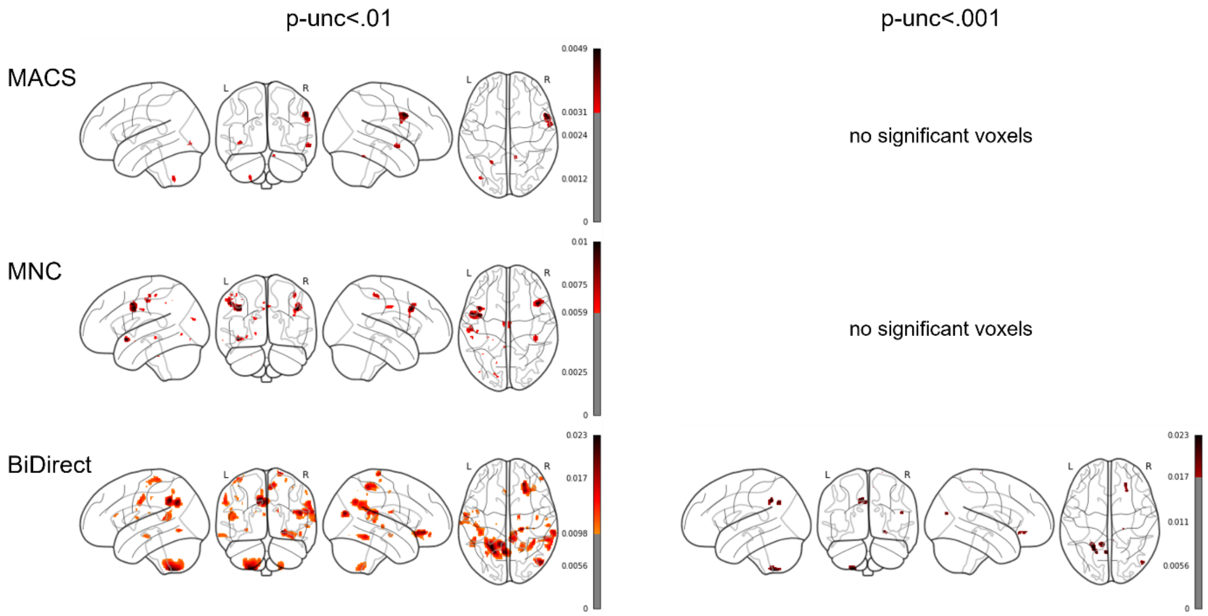


*Note.* Glass brains are shown with maximum intensity projections. Color bars represent the partial $R^2$ of the maltreatment predictor. Results are shown thresholded using the uncorrected thresholds $p_{unc}$<.01 (left column) and $p_{unc}$<.001 (right column).

**Figure S13**

*Significant clusters across cohort-wise analyses (replicability analysis) for* ***model 8*** *– CTQ subscale physical abuse as a predictor in HC and MDD samples controlling for MDD diagnosis*

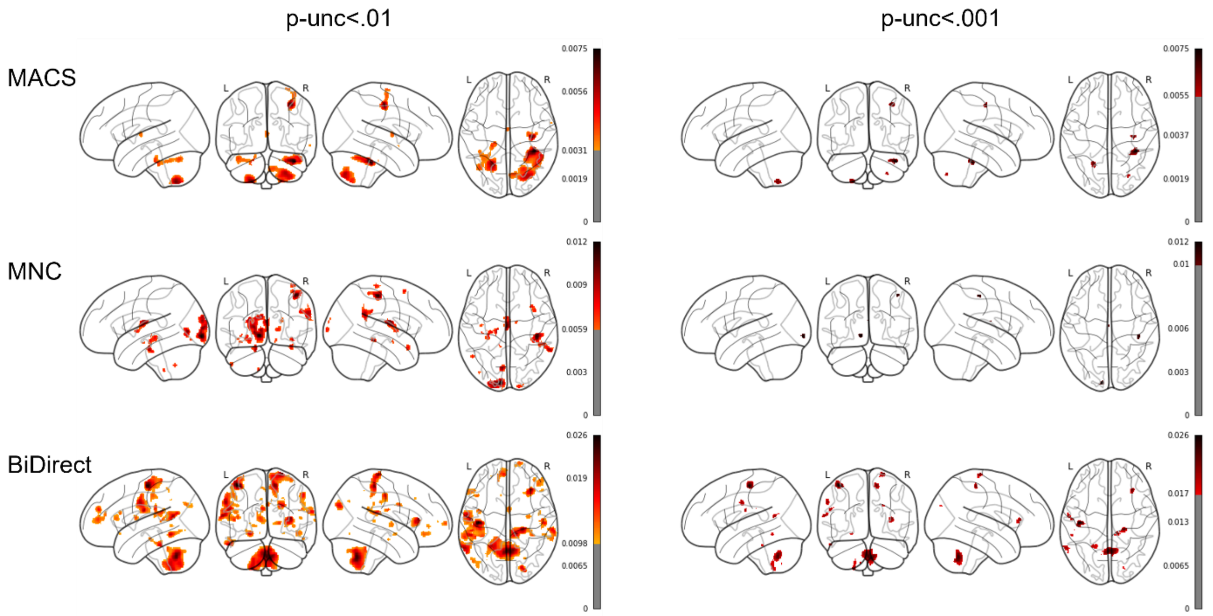


*Note.* Glass brains are shown with maximum intensity projections. Color bars represent the partial $R^2$ of the maltreatment predictor. Results are shown thresholded using the uncorrected thresholds $p_{unc}$<.01 (left column) and $p_{unc}$<.001 (right column).

**Figure S14**

*Significant clusters across cohort-wise analyses (replicability analysis) for* ***model 9*** *– CTQ subscale sexual abuse as a predictor in HC and MDD samples controlling for MDD diagnosis*

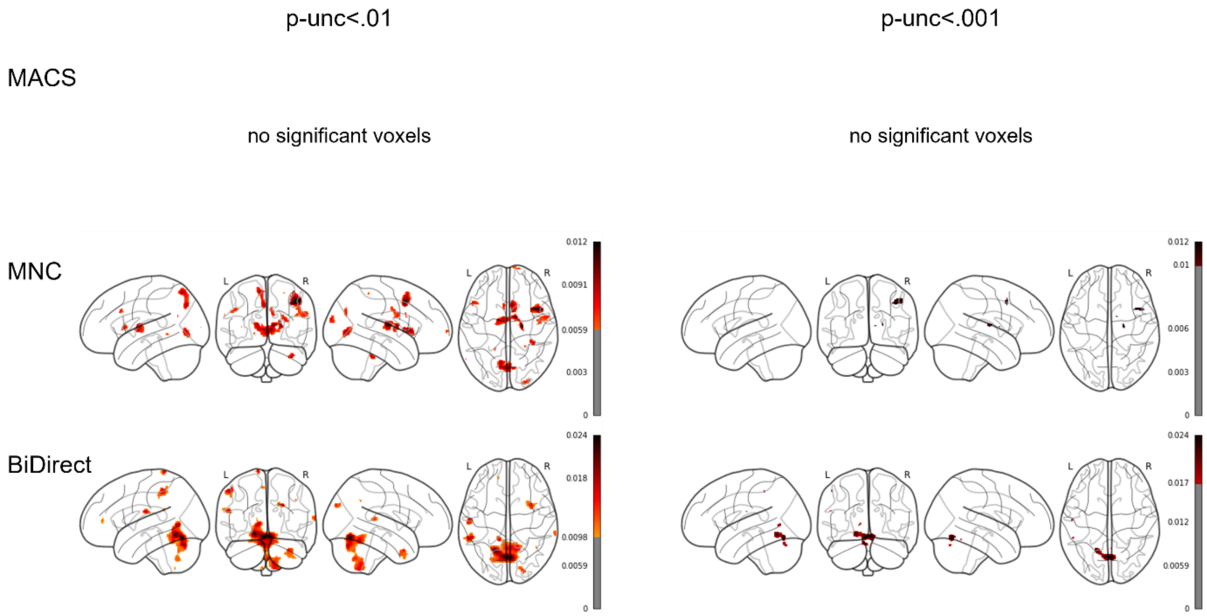


*Note.* Glass brains are shown with maximum intensity projections. Color bars represent the partial $R^2$ of the maltreatment predictor. Results are shown thresholded using the uncorrected thresholds $p_{unc}<.01$ (left column) and $p_{unc}<.001$ (right column).

**Figure S15**

*Significant clusters across cohort-wise analyses (replicability analysis) for **model 10** – CTQ subscale emotional neglect as a predictor in HC and MDD samples controlling for MDD diagnosis*

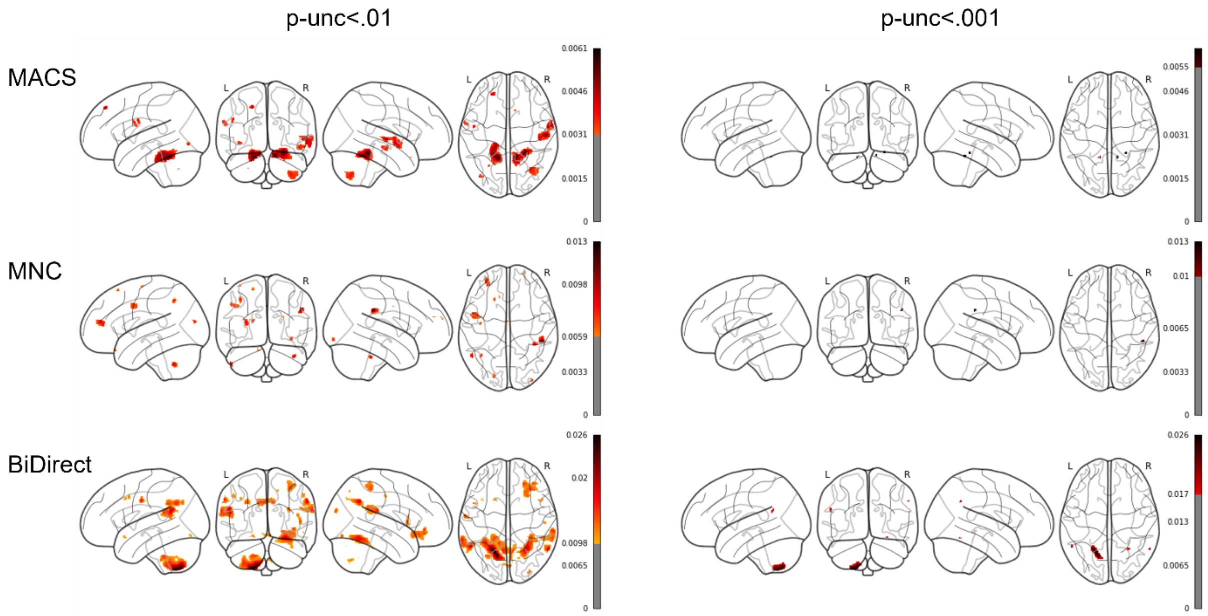


*Note.* Glass brains are shown with maximum intensity projections. Color bars represent the partial $R^2$ of the maltreatment predictor. Results are shown thresholded using the uncorrected thresholds $p_{unc}<.01$ (left column) and $p_{unc}<.001$ (right column).

**Figure S16**

*Significant clusters across cohort-wise analyses (replicability analysis) for **model 11** – CTQ subscale physical neglect as a predictor in HC and MDD samples controlling for MDD diagnosis*

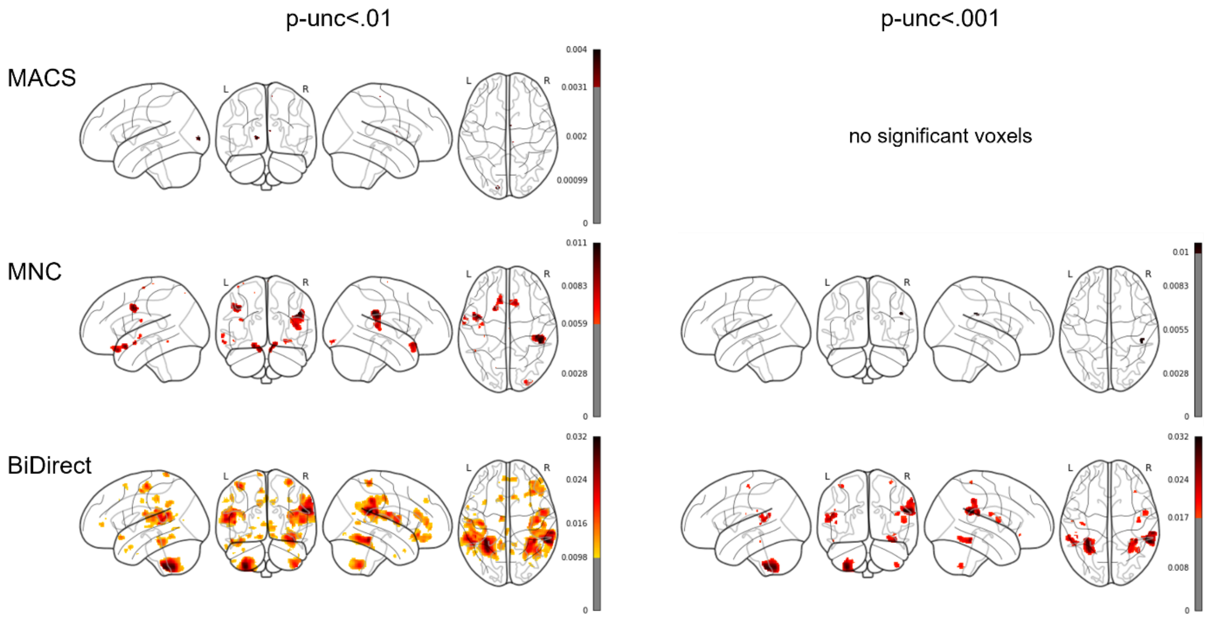


*Note.* Glass brains are shown with maximum intensity projections. Color bars represent the partial $R^2$ of the maltreatment predictor. Results are shown thresholded using the uncorrected thresholds $p_{unc}<.01$ (left column) and $p_{unc}<.001$ (right column).

**Figure S17**

*Significant clusters across cohort-wise analyses (replicability analysis) for* ***model 12*** *– CTQ extreme severity compared to none in HC and MDD samples controlling for MDD diagnosis*

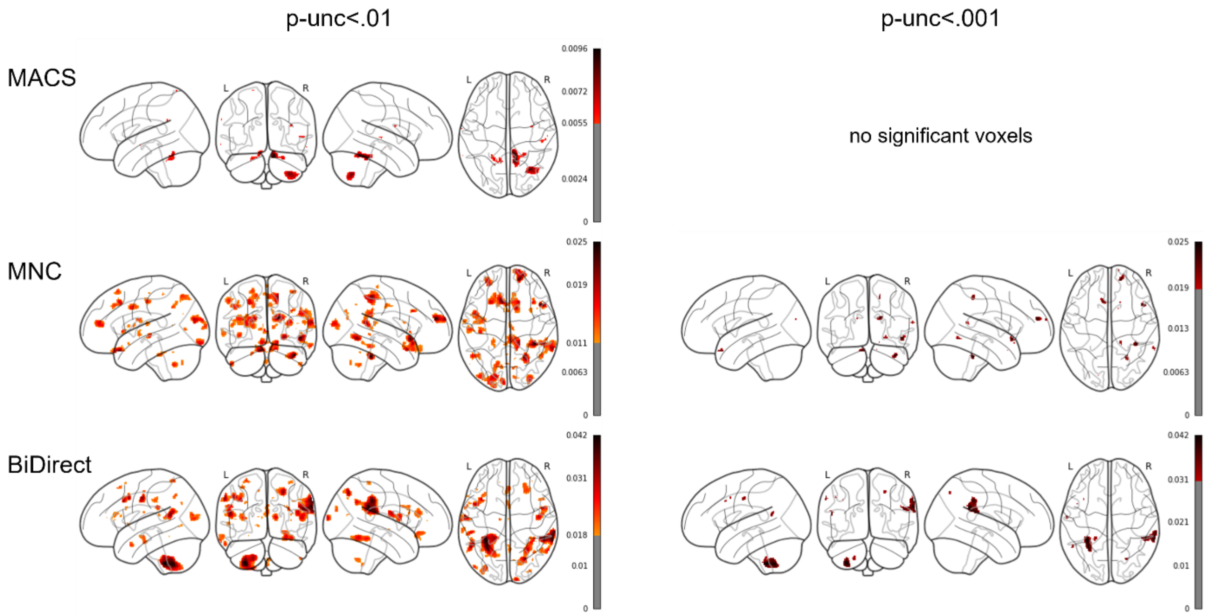


*Note.* Glass brains are shown with maximum intensity projections. Color bars represent the partial $R^2$ of the maltreatment predictor. Results are shown thresholded using the uncorrected thresholds $p_{unc}<.01$ (left column) and $p_{unc}<.001$ (right column).

**Figure S18**

*Significant clusters across cohort-wise analyses (replicability analysis) for* ***model 13*** *– CTQ extreme severity compared to none in HC and MDD samples without controlling for MDD diagnosis*

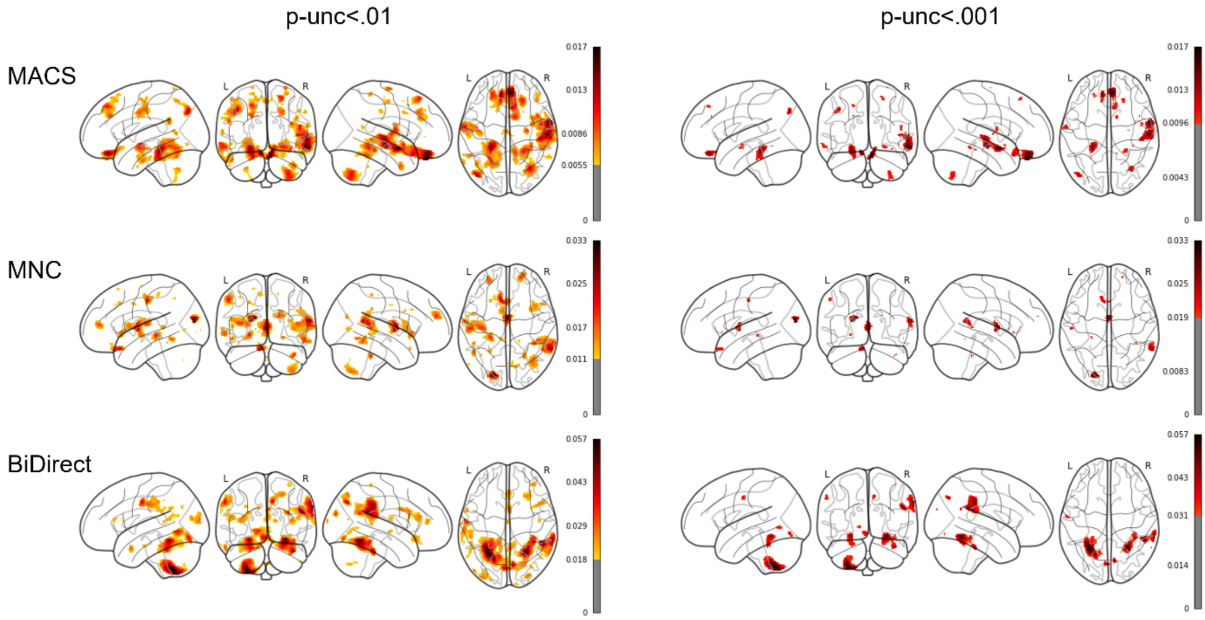


*Note.* Glass brains are shown with maximum intensity projections. Color bars represent the partial $R^2$ of the maltreatment predictor. Results are shown thresholded using the uncorrected thresholds $p_{unc}$<.01 (left column) and $p_{unc}$<.001 (right column).

**Figure S19**

*Significant clusters across cohort-wise analyses (replicability analysis) for* ***model 14*** *– CTQ extreme severity compared to none in HC subsamples*

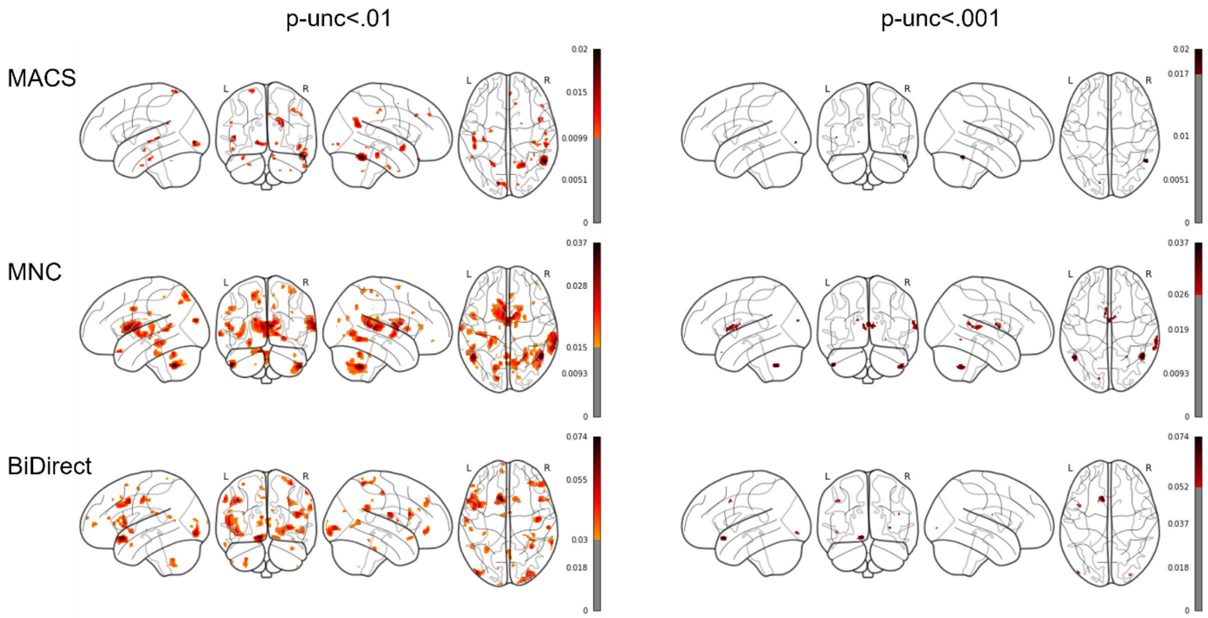


*Note.* Glass brains are shown with maximum intensity projections. Color bars represent the partial $R^2$ of the maltreatment predictor. Results are shown thresholded using the uncorrected thresholds $p_{unc}$<.01 (left column) and $p_{unc}$<.001 (right column).

**Figure S20**

*Significant clusters across cohort-wise analyses (replicability analysis) for* ***model 15*** *– CTQ extreme severity compared to none in MDD subsamples*

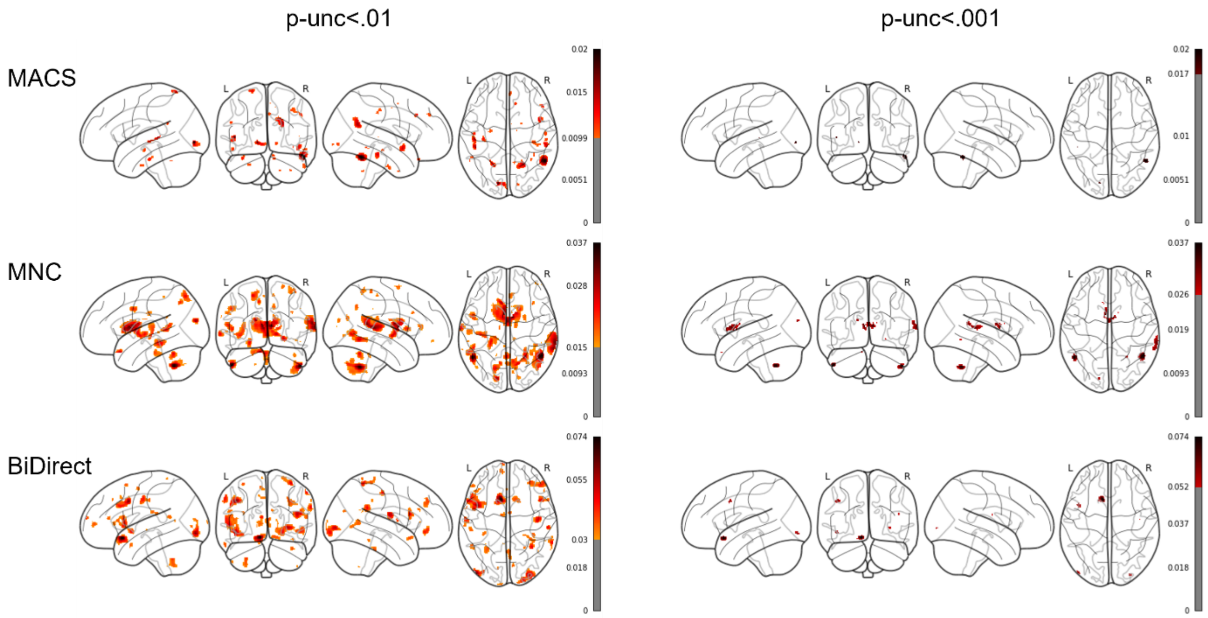


*Note.* Glass brains are shown with maximum intensity projections. Color bars represent the partial $R^2$ of the maltreatment predictor. Results are shown thresholded using the uncorrected thresholds $p_{unc}$<.01 (left column) and $p_{unc}$<.001 (right column).

**Figure S21**

*Significant clusters across cohort-wise analyses (replicability analysis) for* ***model 1*** *– CTQ sum in HC and MDD samples controlling for MDD diagnosis –* ***sex-stratified for female subsample***

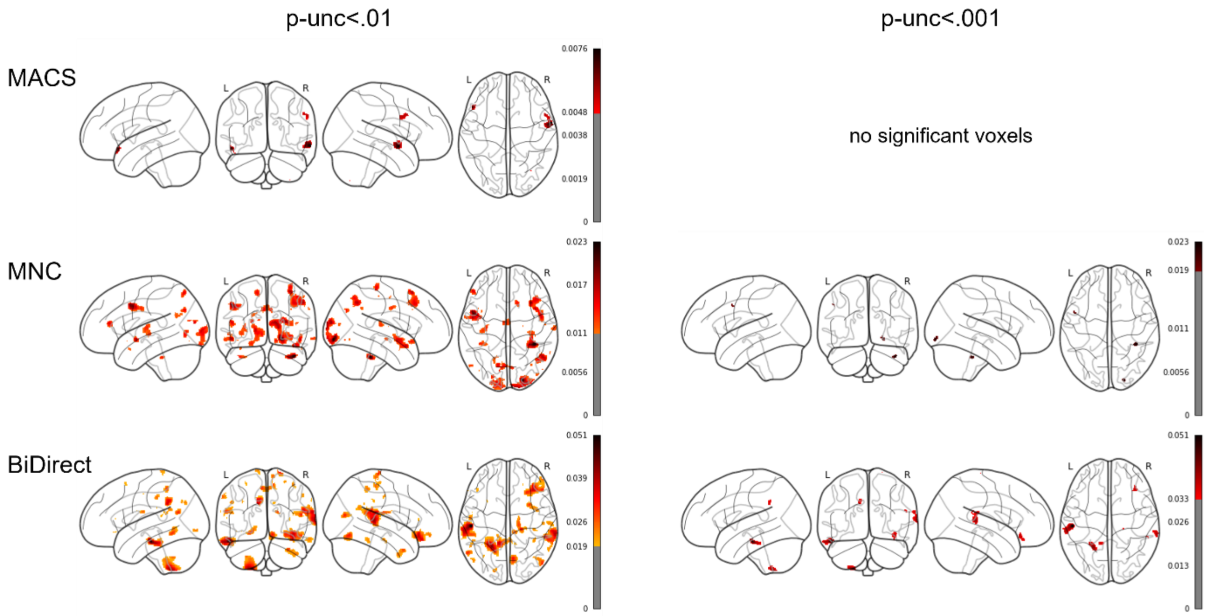


*Note.* Glass brains are shown with maximum intensity projections. Color bars represent the partial $R^2$ of the maltreatment predictor. Results are shown thresholded using the uncorrected thresholds $p_{unc}<.01$ (left column) and $p_{unc}<.001$ (right column).

**Figure S22**

*Significant clusters across cohort-wise analyses (replicability analysis) for **model 2** – CTQ sum as a predictor in HC and MDD samples without controlling for MDD diagnosis – **sex-stratified for female subsample***

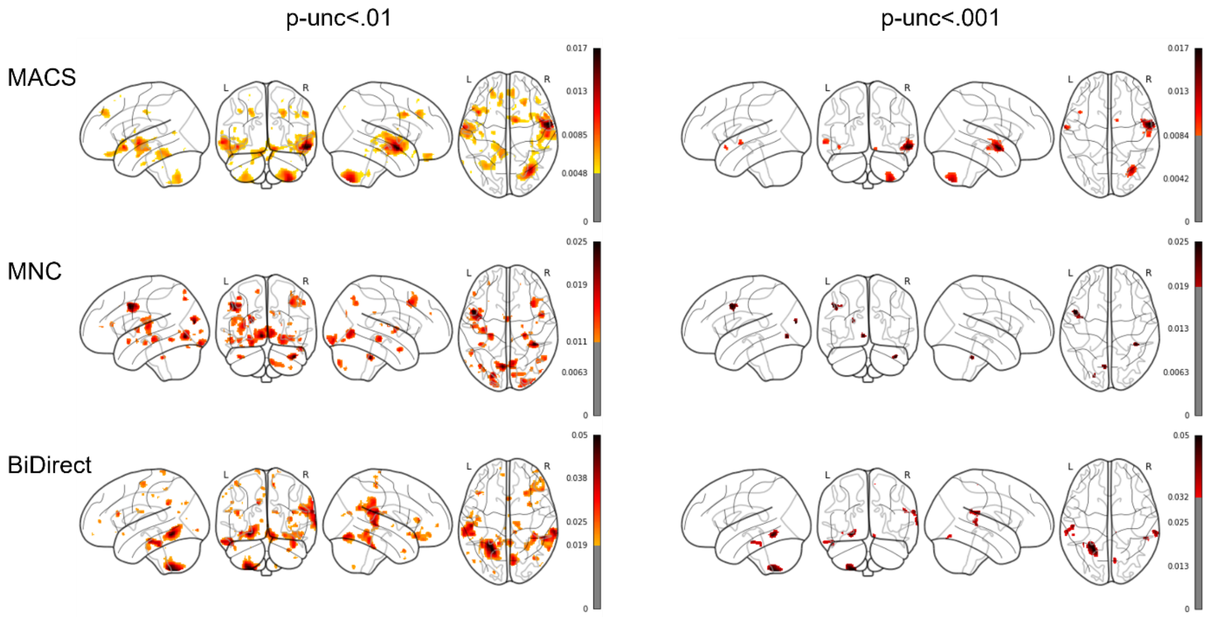


*Note.* Glass brains are shown with maximum intensity projections. Color bars represent the partial $R^2$ of the maltreatment predictor. Results are shown thresholded using the uncorrected thresholds $p_{unc}<.01$ (left column) and $p_{unc}<.001$ (right column).

**Figure S23**

*Significant clusters across cohort-wise analyses (replicability analysis) for* ***model 3*** *– CTQ sum as a predictor in HC subsamples –* ***sex-stratified for female subsample***

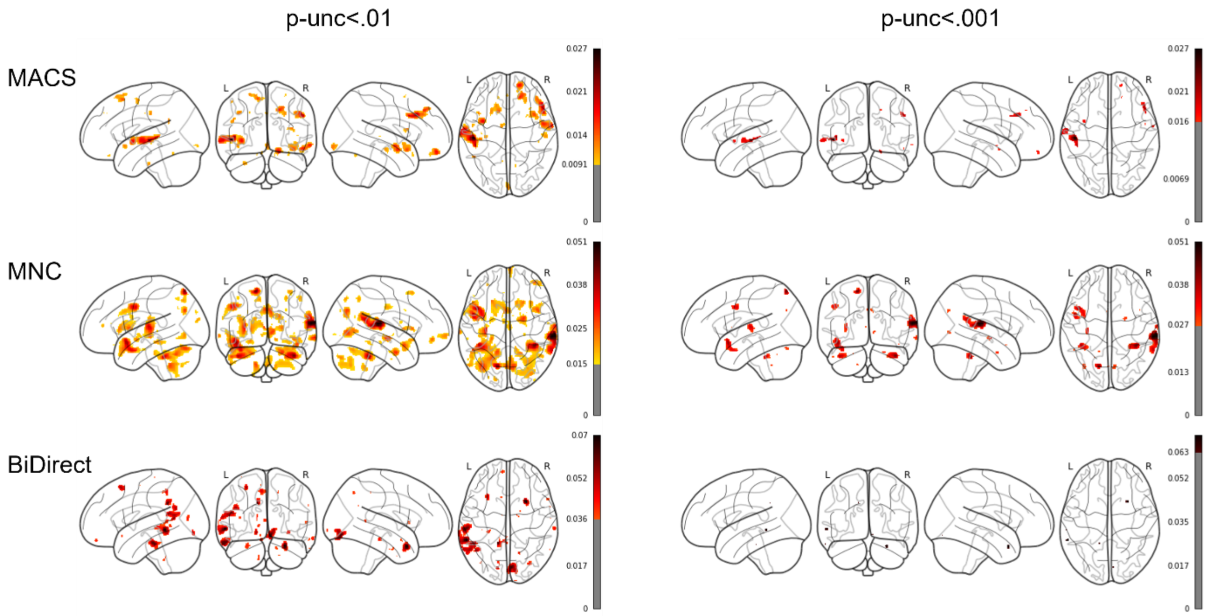


*Note.* Glass brains are shown with maximum intensity projections. Color bars represent the partial $R^2$ of the maltreatment predictor. Results are shown thresholded using the uncorrected thresholds $p_{unc}<.01$ (left column) and $p_{unc}<.001$ (right column).

**Figure S24**

*Significant clusters across cohort-wise analyses (replicability analysis) for* ***model 4*** *– CTQ sum as a predictor in MDD subsamples –* ***sex-stratified for female subsample***

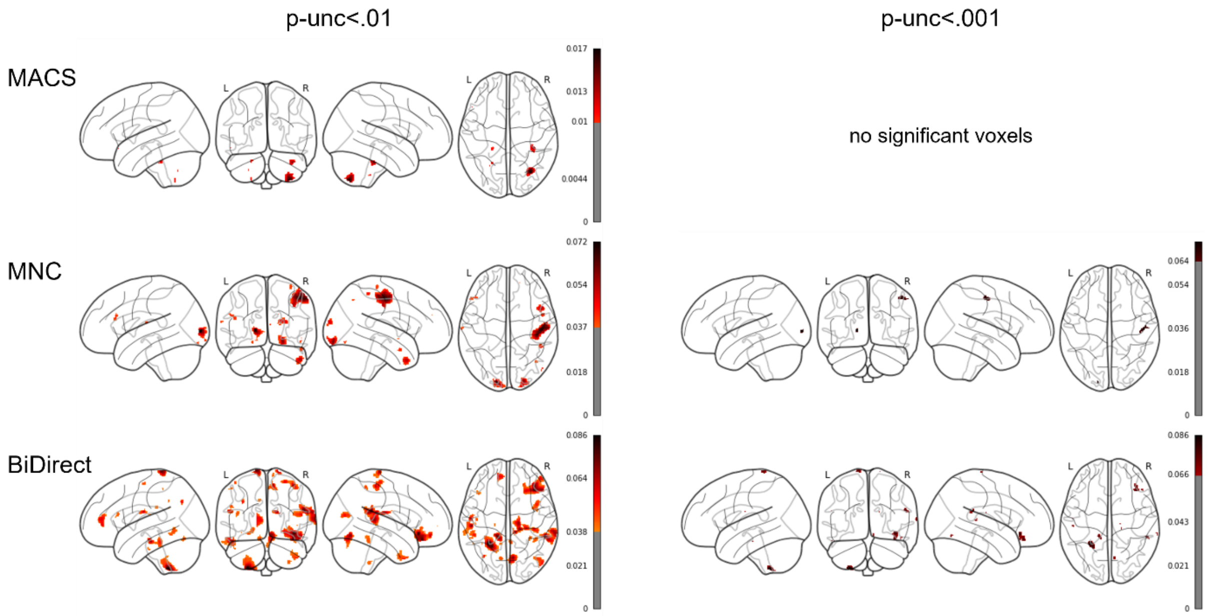


*Note.* Glass brains are shown with maximum intensity projections. Color bars represent the partial $R^2$ of the maltreatment predictor. Results are shown thresholded using the uncorrected thresholds $p_{unc}<.01$ (left column) and $p_{unc}<.001$ (right column).

**Figure S25**

*Significant clusters across cohort-wise analyses (replicability analysis) for **model 5** – CTQ abuse subscales as a predictor in HC and MDD samples controlling for MDD diagnosis – **sex-stratified for female subsample***

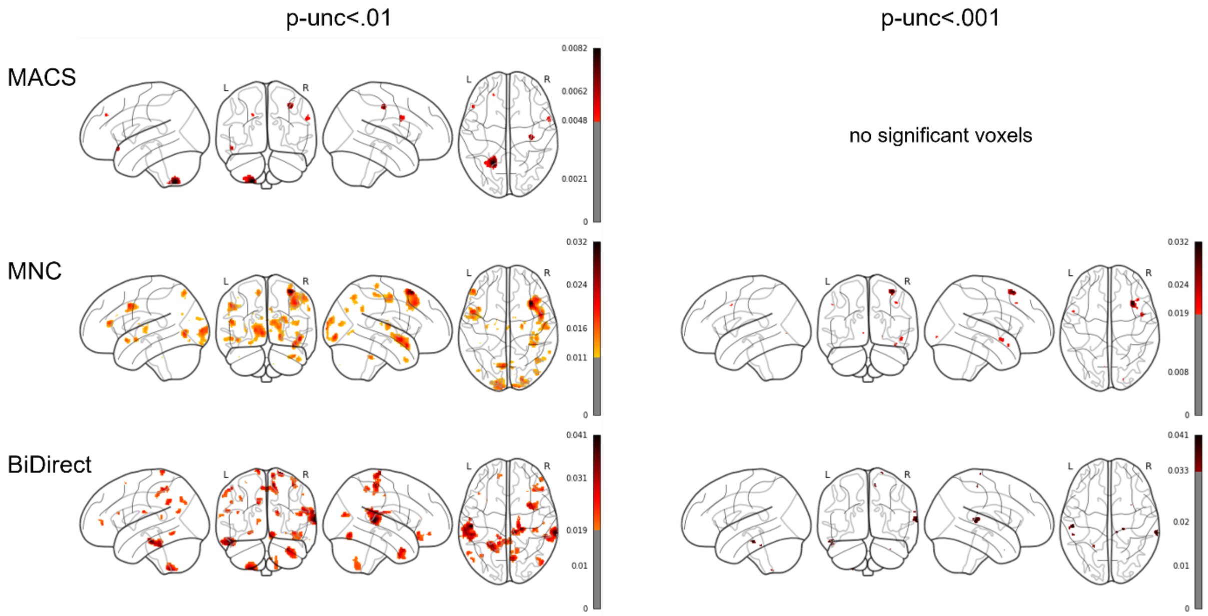


*Note.* Glass brains are shown with maximum intensity projections. Color bars represent the partial $R^2$ of the maltreatment predictor. Results are shown thresholded using the uncorrected thresholds $p_{unc}<.01$ (left column) and $p_{unc}<.001$ (right column).

**Figure S26**

*Significant clusters across cohort-wise analyses (replicability analysis) for* ***model 6*** *– CTQ neglect subscales as a predictor in HC and MDD samples controlling for MDD diagnosis –* ***sex-stratified for female subsample***

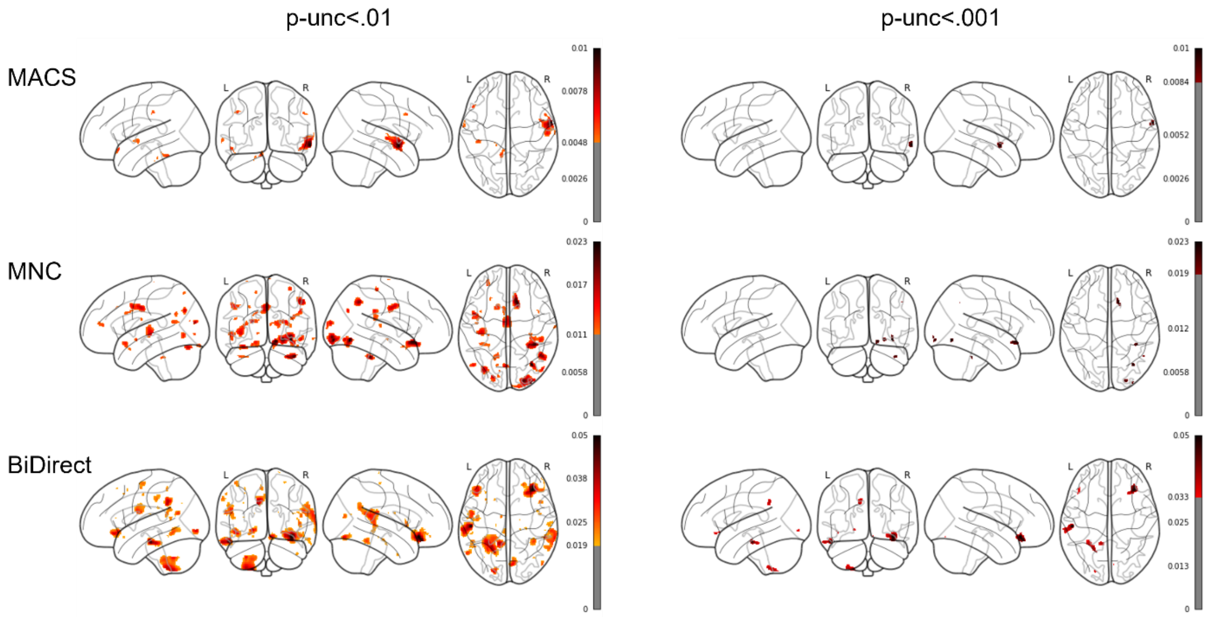


*Note.* Glass brains are shown with maximum intensity projections. Color bars represent the partial $R^2$ of the maltreatment predictor. Results are shown thresholded using the uncorrected thresholds $p_{unc}<.01$ (left column) and $p_{unc}<.001$ (right column).

**Figure S27**

*Significant clusters across cohort-wise analyses (replicability analysis) for* ***model 7*** *– CTQ subscale emotional abuse as a predictor in HC and MDD samples controlling for MDD diagnosis –* ***sex-stratified for female subsample***

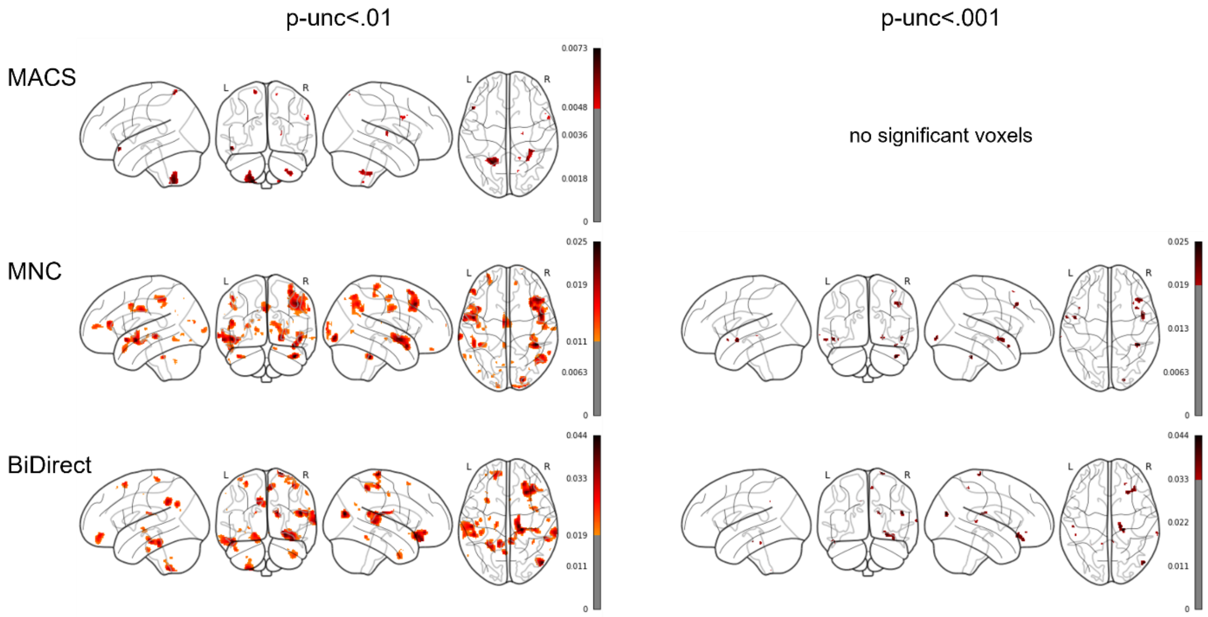


*Note.* Glass brains are shown with maximum intensity projections. Color bars represent the partial $R^2$ of the maltreatment predictor. Results are shown thresholded using the uncorrected thresholds $p_{unc}$<.01 (left column) and $p_{unc}$<.001 (right column).

**Figure S28**

*Significant clusters across cohort-wise analyses (replicability analysis) for **model 8** – CTQ subscale physical abuse as a predictor in HC and MDD samples controlling for MDD diagnosis – **sex-stratified for female subsample***

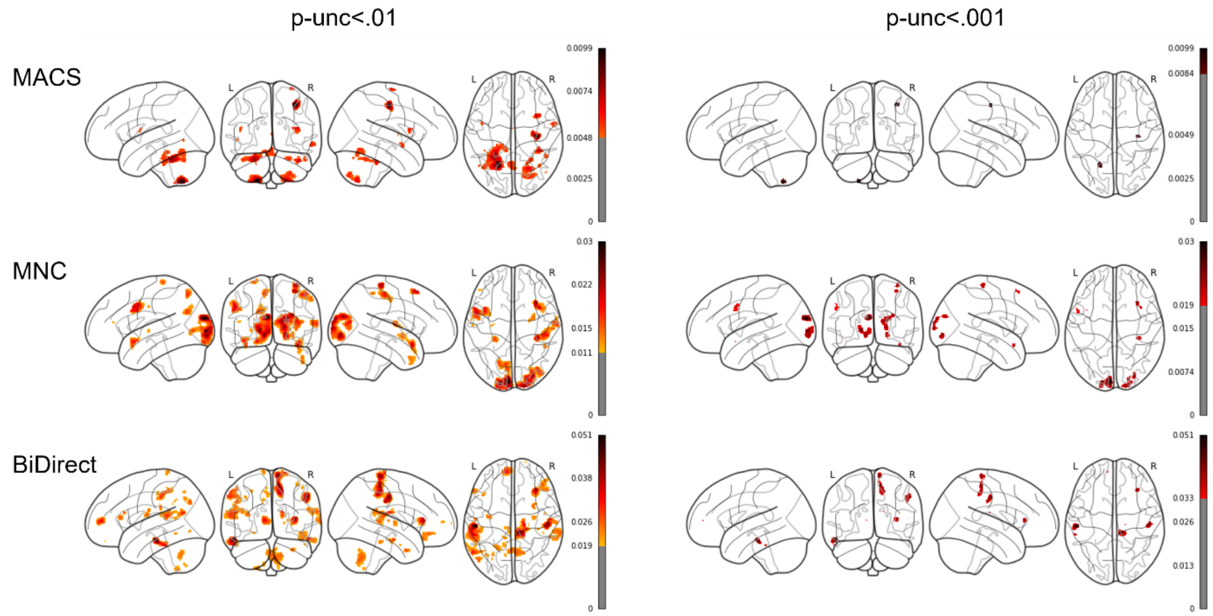


*Note.* Glass brains are shown with maximum intensity projections. Color bars represent the partial $R^2$ of the maltreatment predictor. Results are shown thresholded using the uncorrected thresholds $p_{unc}<.01$ (left column) and $p_{unc}<.001$ (right column).

**Figure S29**

*Significant clusters across cohort-wise analyses (replicability analysis) for* ***model 9*** *– CTQ subscale sexual abuse as a predictor in HC and MDD samples controlling for MDD diagnosis –* ***sex-stratified for female subsample***

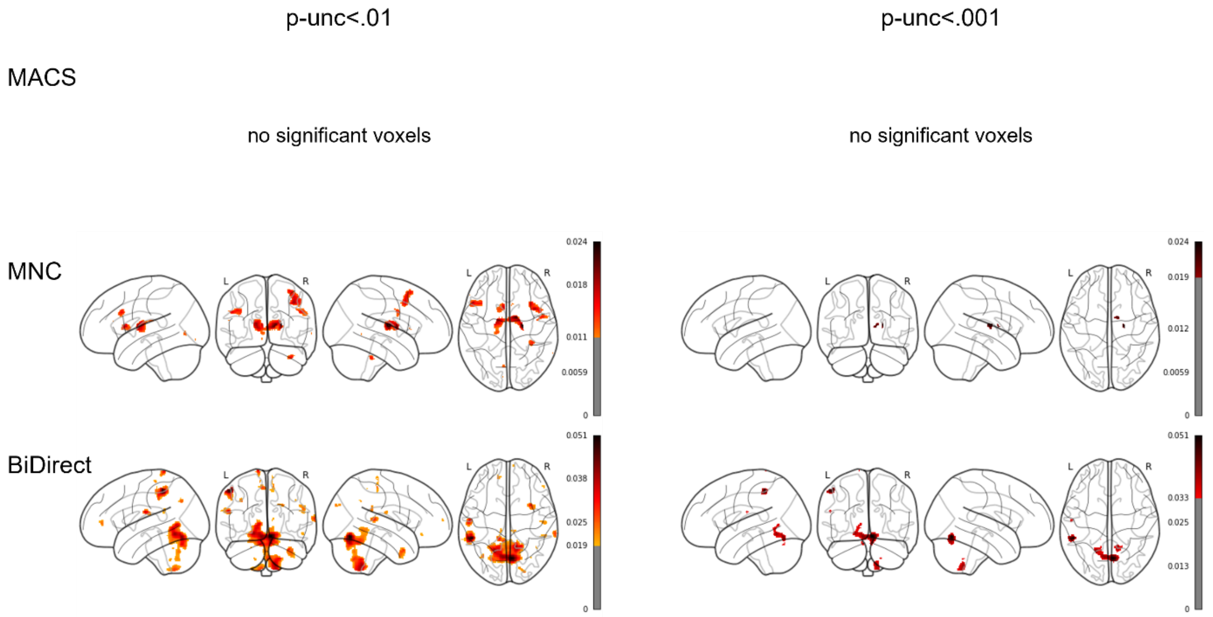


*Note.* Glass brains are shown with maximum intensity projections. Color bars represent the partial $R^2$ of the maltreatment predictor. Results are shown thresholded using the uncorrected thresholds $p_{unc}$<.01 (left column) and $p_{unc}$<.001 (right column).

**Figure S30**

*Significant clusters across cohort-wise analyses (replicability analysis) for* ***model 10*** *– CTQ subscale emotional neglect as a predictor in HC and MDD samples controlling for MDD diagnosis –* ***sex-stratified for female subsample***

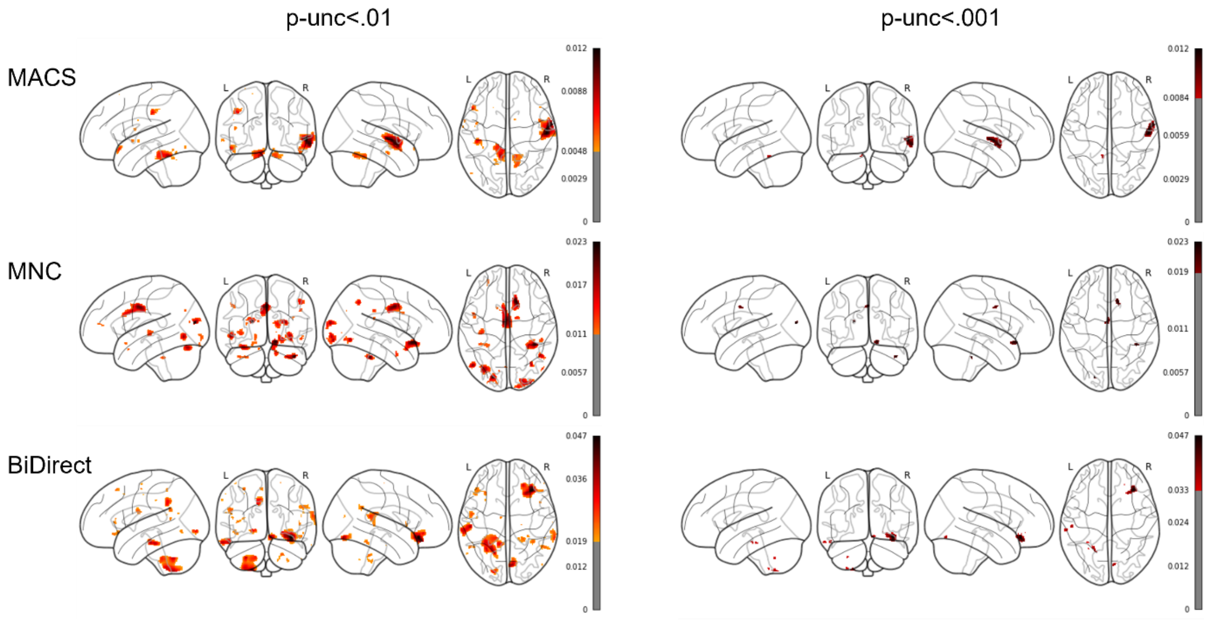


*Note.* Glass brains are shown with maximum intensity projections. Color bars represent the partial $R^2$ of the maltreatment predictor. Results are shown thresholded using the uncorrected thresholds $p_{unc}<.01$ (left column) and $p_{unc}<.001$ (right column).

**Figure S31**

*Significant clusters across cohort-wise analyses (replicability analysis) for* ***model 11*** *– CTQ subscale physical neglect as a predictor in HC and MDD samples controlling for MDD diagnosis –* ***sex-stratified for female subsample***

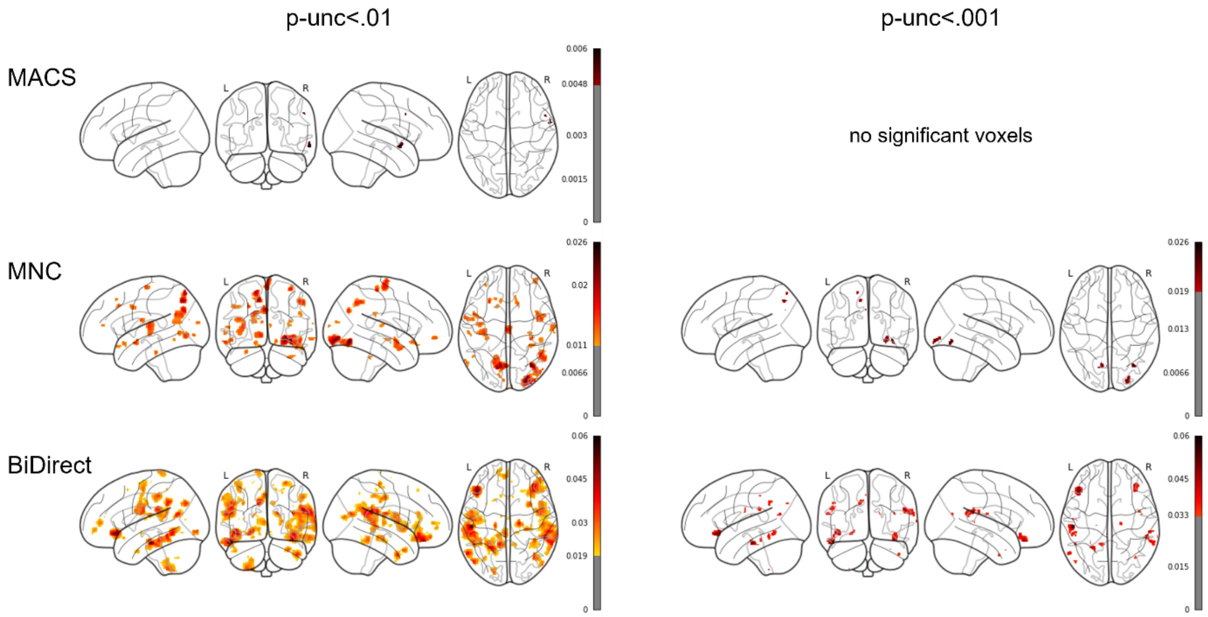


*Note.* Glass brains are shown with maximum intensity projections. Color bars represent the partial $R^2$ of the maltreatment predictor. Results are shown thresholded using the uncorrected thresholds $p_{unc}<.01$ (left column) and $p_{unc}<.001$ (right column).

**Figure S32**

*Significant clusters across cohort-wise analyses (replicability analysis) for* ***model 12*** *– CTQ extreme severity compared to none in HC and MDD samples controlling for MDD diagnosis –* ***sex-stratified for female subsample***

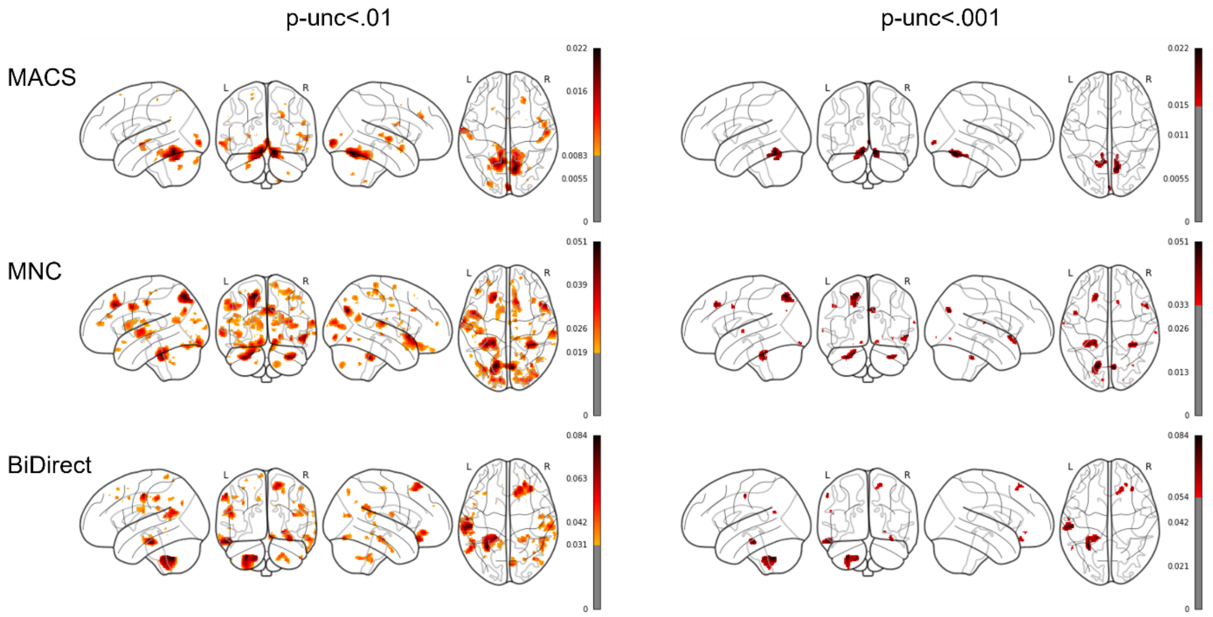


*Note.* Glass brains are shown with maximum intensity projections. Color bars represent the partial $R^2$ of the maltreatment predictor. Results are shown thresholded using the uncorrected thresholds $p_{unc}$<.01 (left column) and $p_{unc}$<.001 (right column).

**Figure S33**

*Significant clusters across cohort-wise analyses (replicability analysis) for* ***model 13*** *– CTQ extreme severity compared to none in HC and MDD samples without controlling for MDD diagnosis –* ***sex-stratified for female subsample***

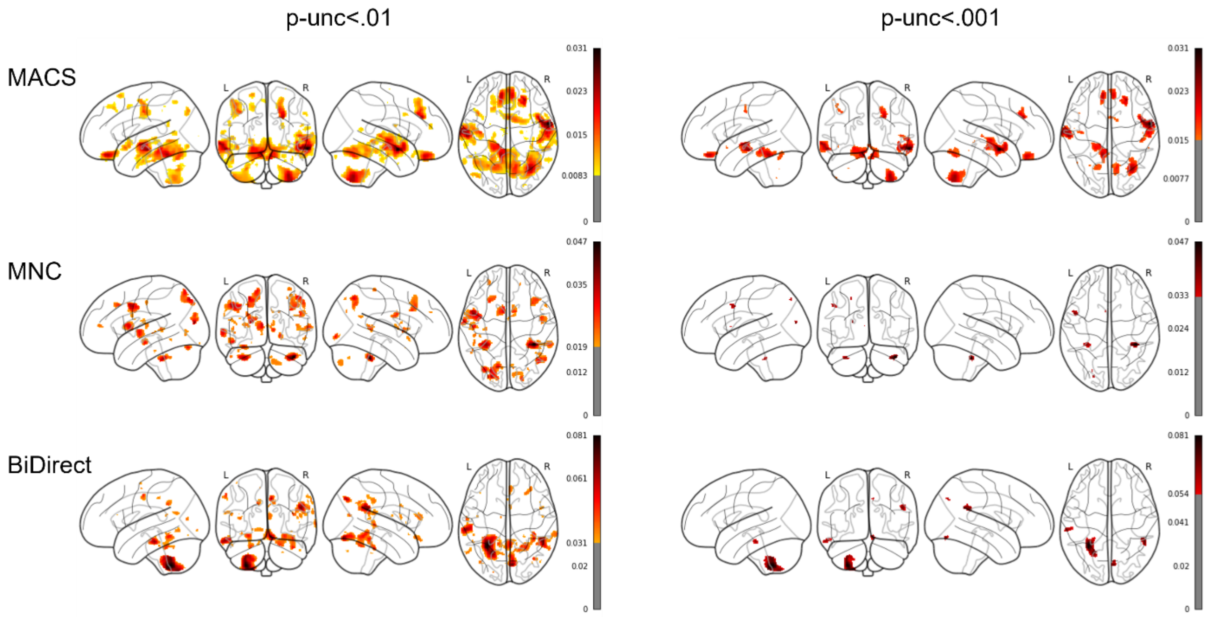


*Note.* Glass brains are shown with maximum intensity projections. Color bars represent the partial $R^2$ of the maltreatment predictor. Results are shown thresholded using the uncorrected thresholds $p_{unc}<.01$ (left column) and $p_{unc}<.001$ (right column).

**Figure S34**

*Significant clusters across cohort-wise analyses (replicability analysis) for **model 14** – CTQ extreme severity compared to none in HC subsamples – **sex-stratified for female subsample***

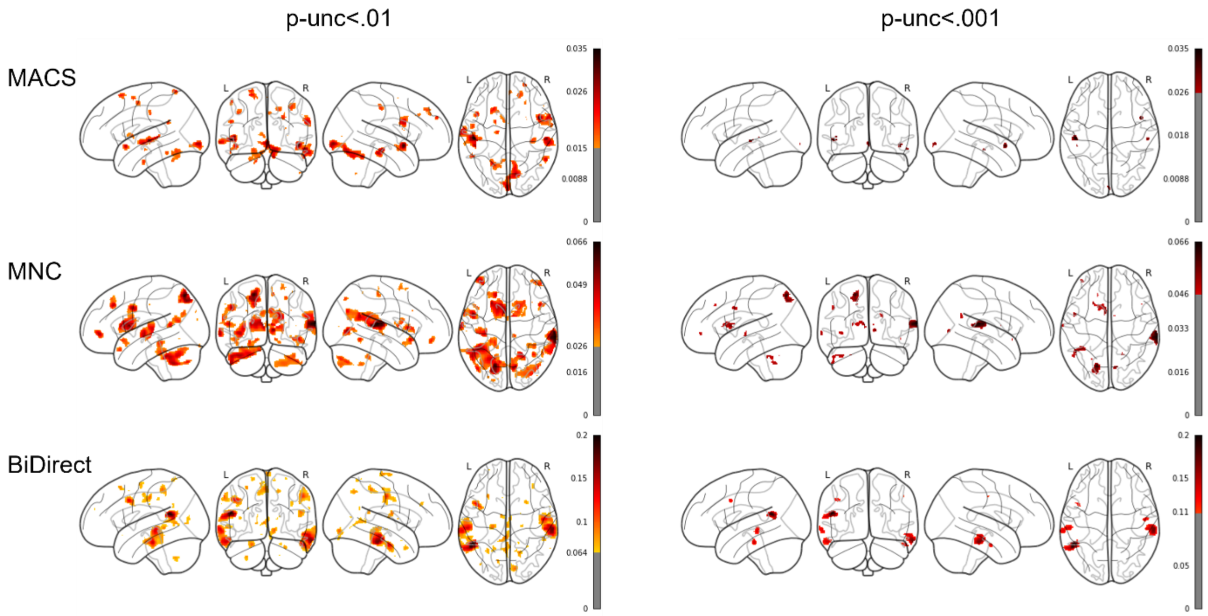


*Note.* Glass brains are shown with maximum intensity projections. Color bars represent the partial $R^2$ of the maltreatment predictor. Results are shown thresholded using the uncorrected thresholds $p_{unc}<.01$ (left column) and $p_{unc}<.001$ (right column).

**Figure S35**

*Significant clusters across cohort-wise analyses (replicability analysis) for **model 15** – CTQ extreme severity compared to none in MDD subsamples – **sex-stratified for female subsample***

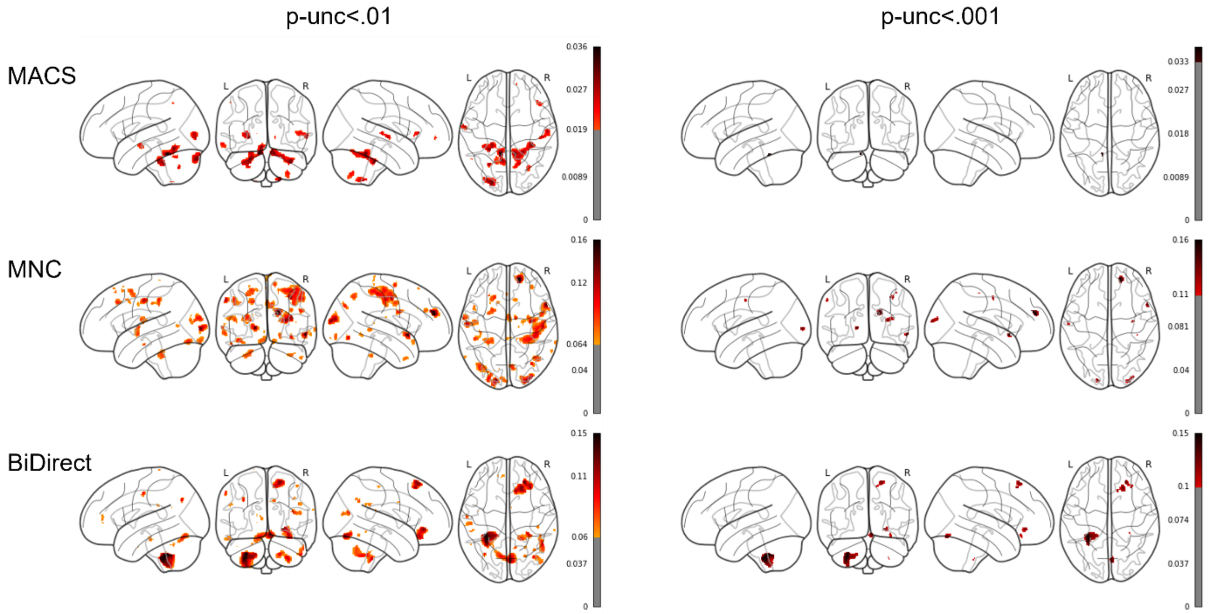


*Note.* Glass brains are shown with maximum intensity projections. Color bars represent the partial $R^2$ of the maltreatment predictor. Results are shown thresholded using the uncorrected thresholds $p_{unc}<.01$ (left column) and $p_{unc}<.001$ (right column).

**Figure S36**

*Significant clusters across cohort-wise analyses (replicability analysis) for* ***model 1*** *– CTQ sum in HC and MDD samples controlling for MDD diagnosis –* ***sex-stratified for male subsample***

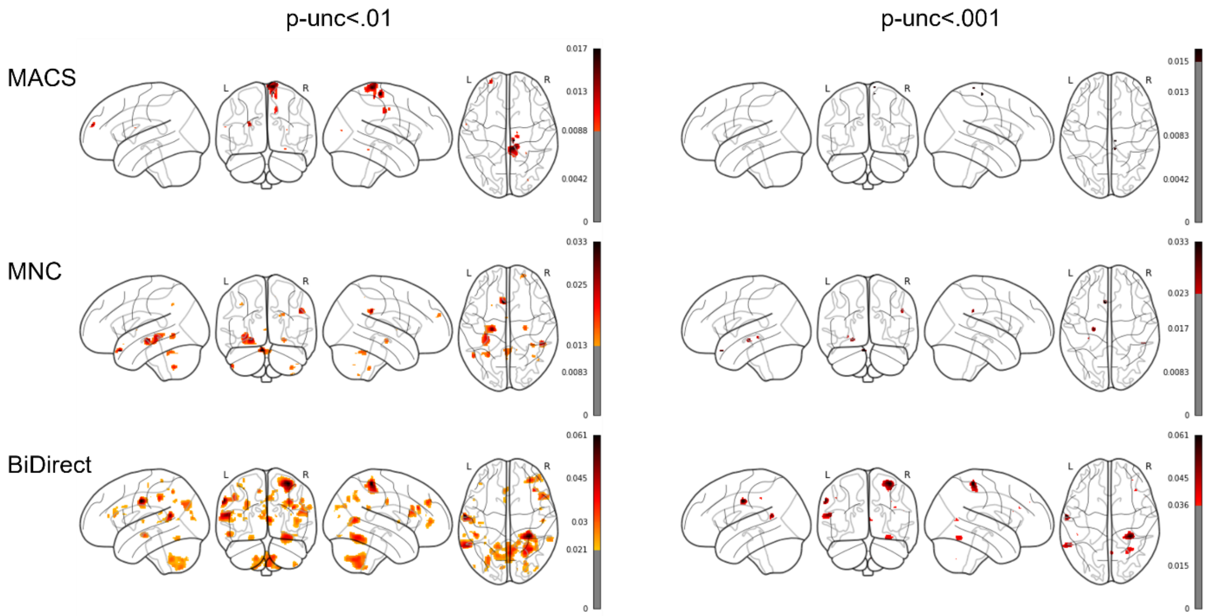


*Note.* Glass brains are shown with maximum intensity projections. Color bars represent the partial $R^2$ of the maltreatment predictor. Results are shown thresholded using the uncorrected thresholds $p_{unc}$<.01 (left column) and $p_{unc}$<.001 (right column).

**Figure S37**

*Significant clusters across cohort-wise analyses (replicability analysis) for* ***model 2*** *– CTQ sum as a predictor in HC and MDD samples without controlling for MDD diagnosis –* ***sex-stratified for male subsample***

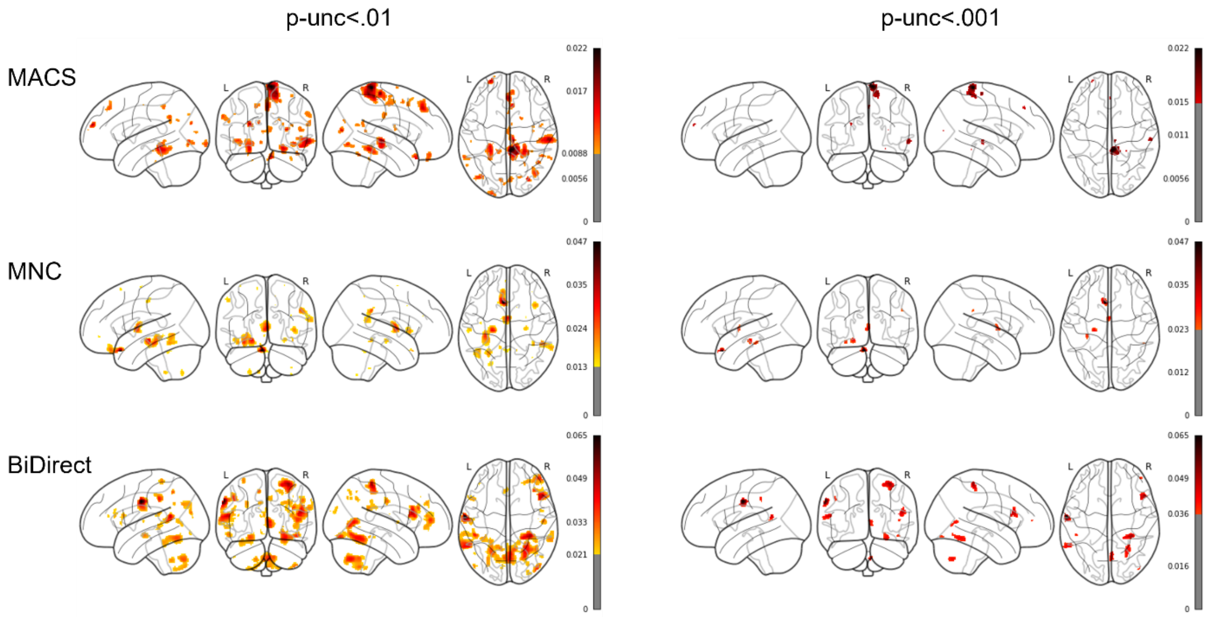


*Note.* Glass brains are shown with maximum intensity projections. Color bars represent the partial $R^2$ of the maltreatment predictor. Results are shown thresholded using the uncorrected thresholds $p_{unc}<.01$ (left column) and $p_{unc}<.001$ (right column).

**Figure S38**

*Significant clusters across cohort-wise analyses (replicability analysis) for* ***model 3*** *– CTQ sum as a predictor in HC subsamples –* ***sex-stratified for male subsample***

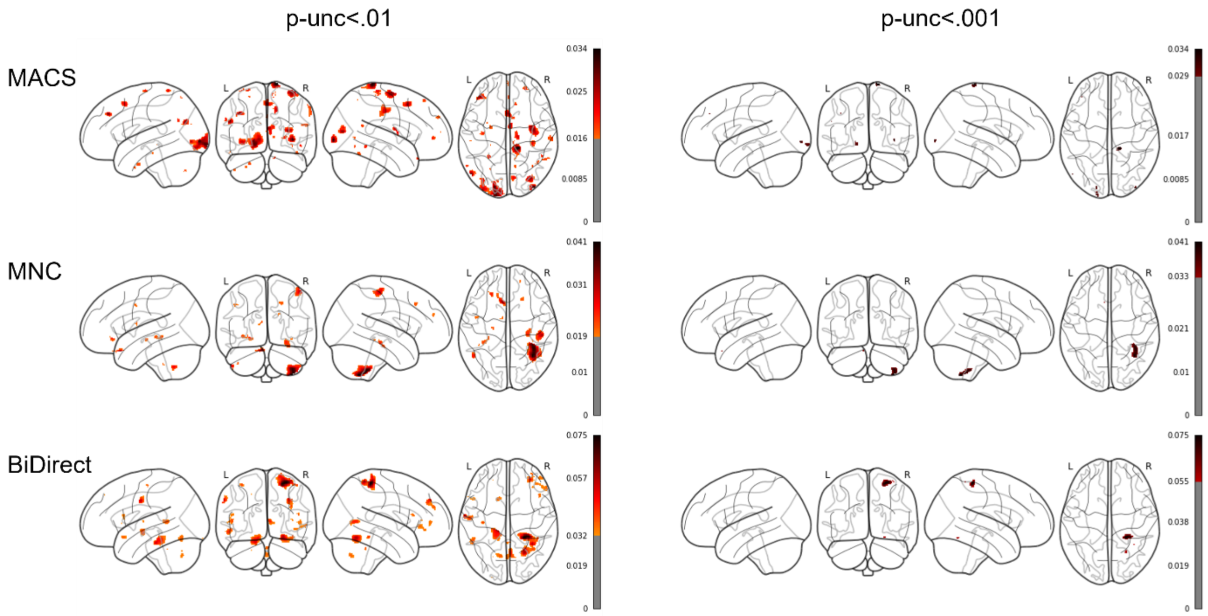


*Note.* Glass brains are shown with maximum intensity projections. Color bars represent the partial $R^2$ of the maltreatment predictor. Results are shown thresholded using the uncorrected thresholds $p_{unc}<.01$ (left column) and $p_{unc}<.001$ (right column).

**Figure S39**

*Significant clusters across cohort-wise analyses (replicability analysis) for* ***model 4*** *– CTQ sum as a predictor in MDD subsamples –* ***sex-stratified for male subsample***

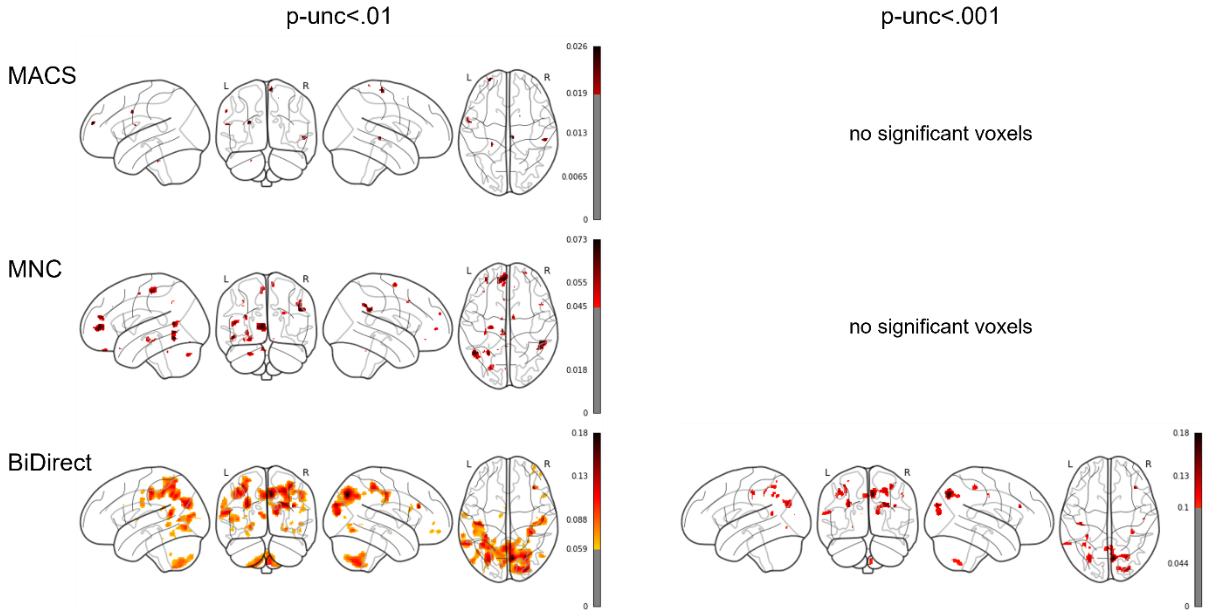


*Note.* Glass brains are shown with maximum intensity projections. Color bars represent the partial $R^2$ of the maltreatment predictor. Results are shown thresholded using the uncorrected thresholds $p_{unc}<.01$ (left column) and $p_{unc}<.001$ (right column).

**Figure S40**

*Significant clusters across cohort-wise analyses (replicability analysis) for* ***model 5*** *– CTQ abuse subscales as a predictor in HC and MDD samples controlling for MDD diagnosis –* ***sex-stratified for male subsample***

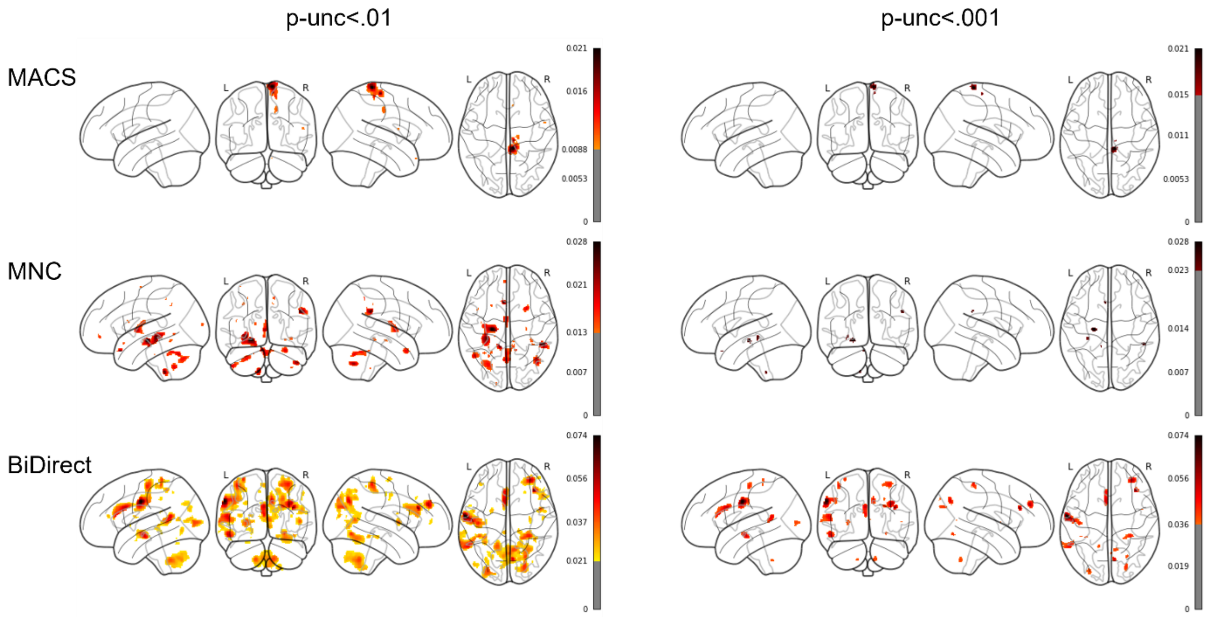


*Note.* Glass brains are shown with maximum intensity projections. Color bars represent the partial $R^2$ of the maltreatment predictor. Results are shown thresholded using the uncorrected thresholds $p_{unc}<.01$ (left column) and $p_{unc}<.001$ (right column).

**Figure S41**

*Significant clusters across cohort-wise analyses (replicability analysis) for* ***model 6*** *– CTQ neglect subscales as a predictor in HC and MDD samples controlling for MDD diagnosis –* ***sex-stratified for male subsample***

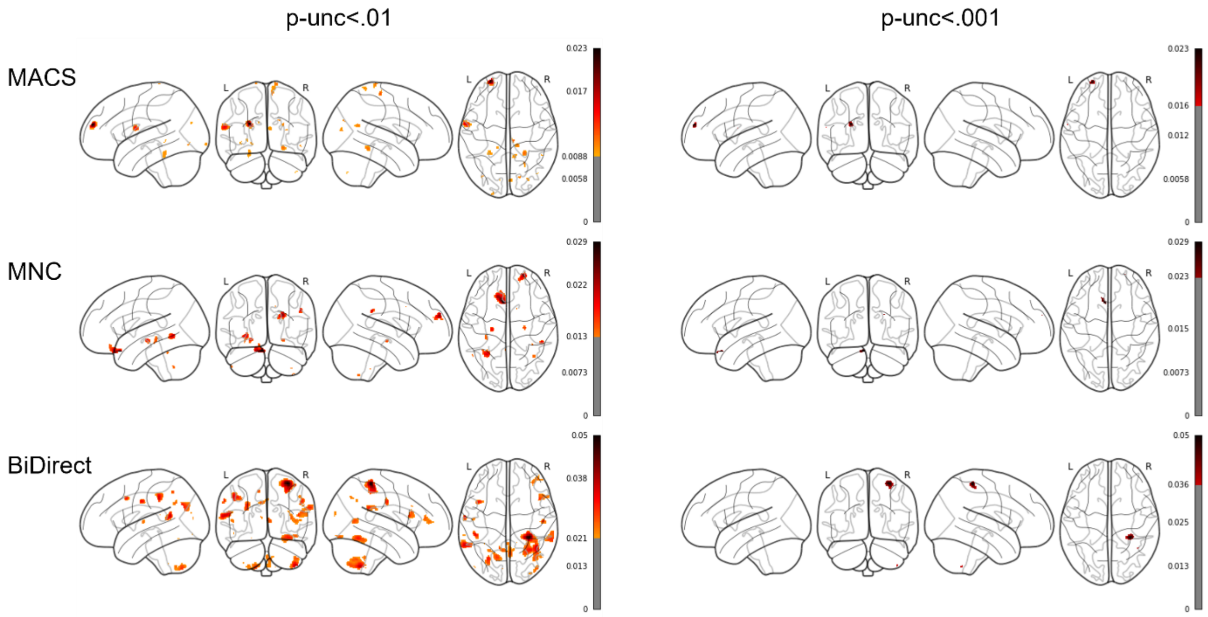


*Note.* Glass brains are shown with maximum intensity projections. Color bars represent the partial $R^2$ of the maltreatment predictor. Results are shown thresholded using the uncorrected thresholds $p_{unc}<.01$ (left column) and $p_{unc}<.001$ (right column).

**Figure S42**

*Significant clusters across cohort-wise analyses (replicability analysis) for* ***model 7*** *– CTQ subscale emotional abuse as a predictor in HC and MDD samples controlling for MDD diagnosis –* ***sex-stratified for male subsample***

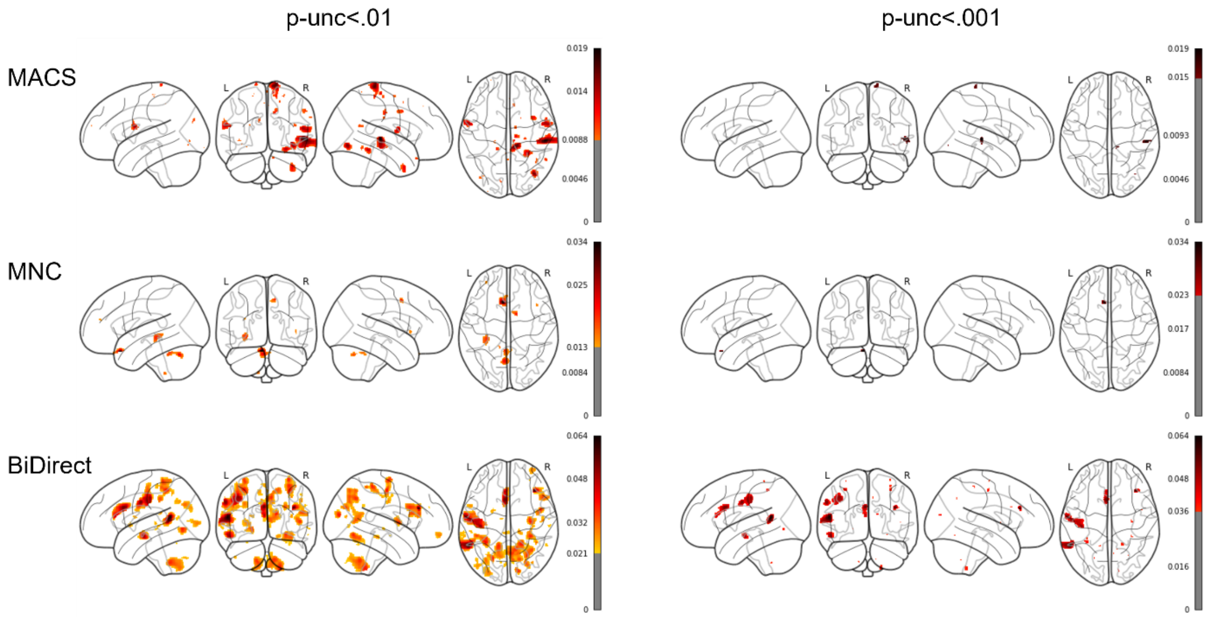


*Note.* Glass brains are shown with maximum intensity projections. Color bars represent the partial $R^2$ of the maltreatment predictor. Results are shown thresholded using the uncorrected thresholds $p_{unc}$<.01 (left column) and $p_{unc}$<.001 (right column).

**Figure S43**

*Significant clusters across cohort-wise analyses (replicability analysis) for **model 8** – CTQ subscale physical abuse as a predictor in HC and MDD samples controlling for MDD diagnosis – **sex-stratified for male subsample***

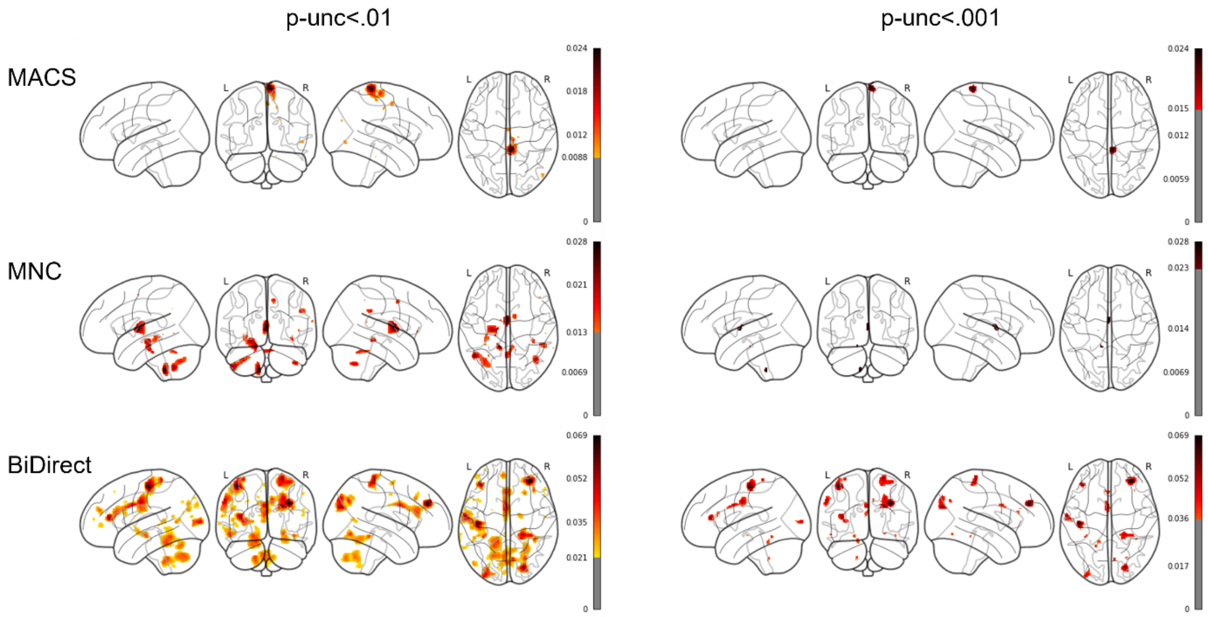


*Note.* Glass brains are shown with maximum intensity projections. Color bars represent the partial $R^2$ of the maltreatment predictor. Results are shown thresholded using the uncorrected thresholds $p_{unc}<.01$ (left column) and $p_{unc}<.001$ (right column).

**Figure S44**

*Significant clusters across cohort-wise analyses (replicability analysis) for **model 9** – CTQ subscale sexual abuse as a predictor in HC and MDD samples controlling for MDD diagnosis – **sex-stratified for male subsample***

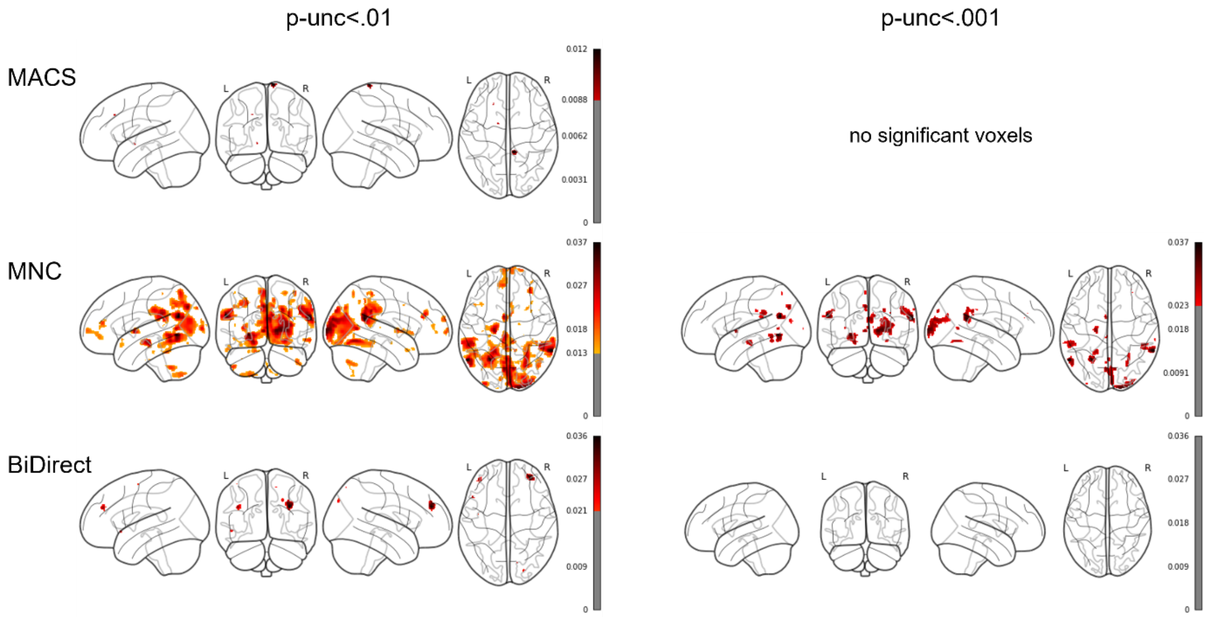


*Note.* Glass brains are shown with maximum intensity projections. Color bars represent the partial $R^2$ of the maltreatment predictor. Results are shown thresholded using the uncorrected thresholds $p_{unc}<.01$ (left column) and $p_{unc}<.001$ (right column).

**Figure S45**

*Significant clusters across cohort-wise analyses (replicability analysis) for **model 10** – CTQ subscale emotional neglect as a predictor in HC and MDD samples controlling for MDD diagnosis – **sex-stratified for male subsample***

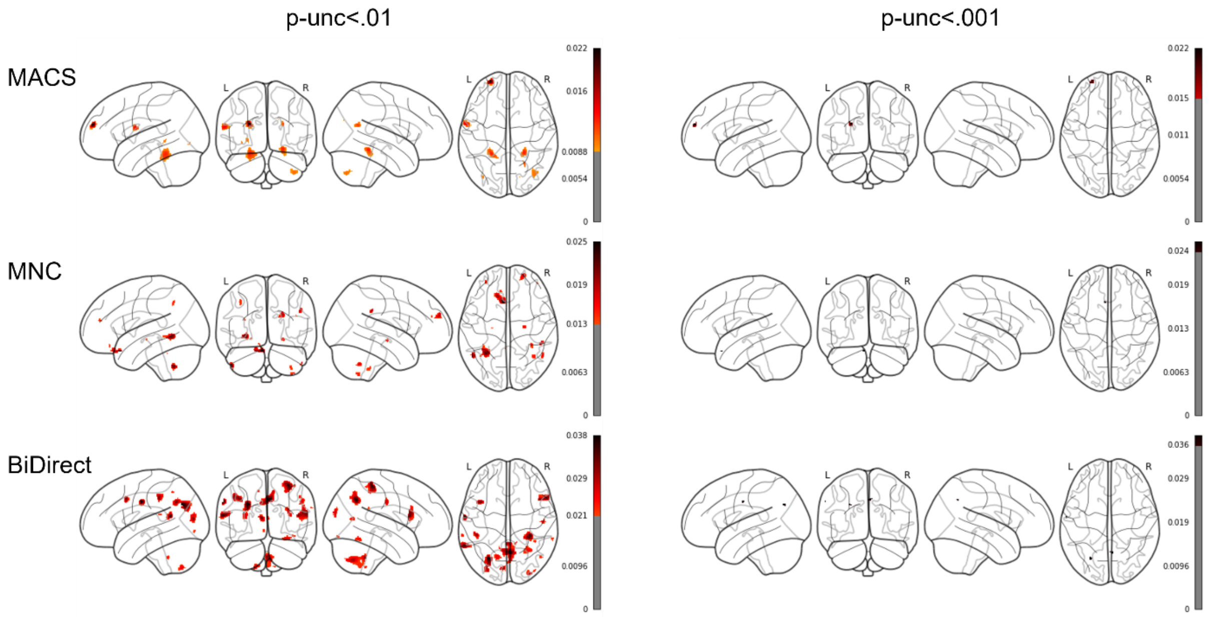


*Note.* Glass brains are shown with maximum intensity projections. Color bars represent the partial $R^2$ of the maltreatment predictor. Results are shown thresholded using the uncorrected thresholds $p_{unc}<.01$ (left column) and $p_{unc}<.001$ (right column).

**Figure S46**

*Significant clusters across cohort-wise analyses (replicability analysis) for* ***model 11*** *– CTQ subscale physical neglect as a predictor in HC and MDD samples controlling for MDD diagnosis –* ***sex-stratified for male subsample***

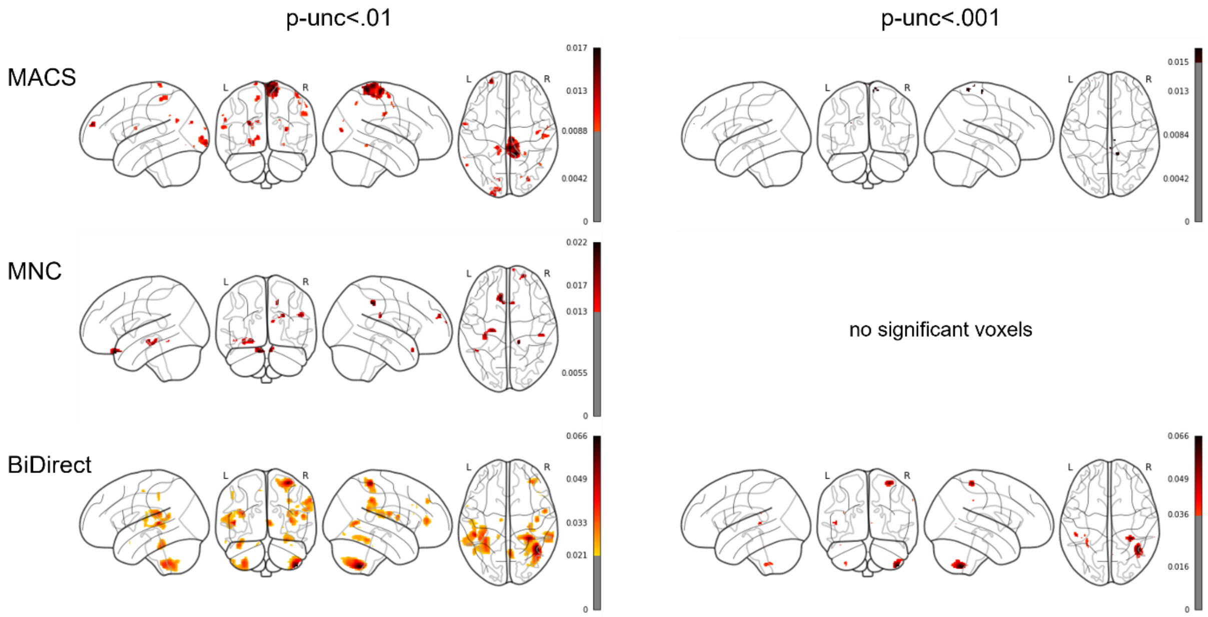


*Note.* Glass brains are shown with maximum intensity projections. Color bars represent the partial $R^2$ of the maltreatment predictor. Results are shown thresholded using the uncorrected thresholds $p_{unc}<.01$ (left column) and $p_{unc}<.001$ (right column).

**Figure S47**

*Significant clusters across cohort-wise analyses (replicability analysis) for* ***model 12*** *– CTQ extreme severity compared to none in HC and MDD samples controlling for MDD diagnosis –* ***sex-stratified for male subsample***

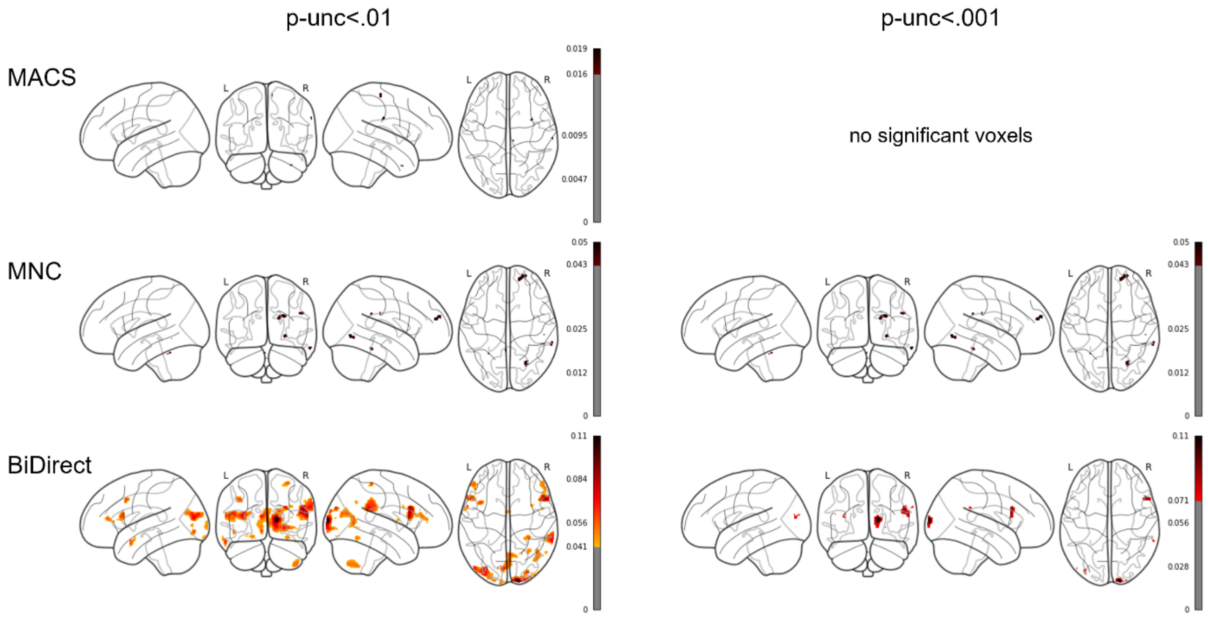


*Note.* Glass brains are shown with maximum intensity projections. Color bars represent the partial $R^2$ of the maltreatment predictor. Results are shown thresholded using the uncorrected thresholds $p_{unc}<.01$ (left column) and $p_{unc}<.001$ (right column).

**Figure S48**

*Significant clusters across cohort-wise analyses (replicability analysis) for* ***model 13*** *– CTQ extreme severity compared to none in HC and MDD samples without controlling for MDD diagnosis –* ***sex-stratified for male subsample***

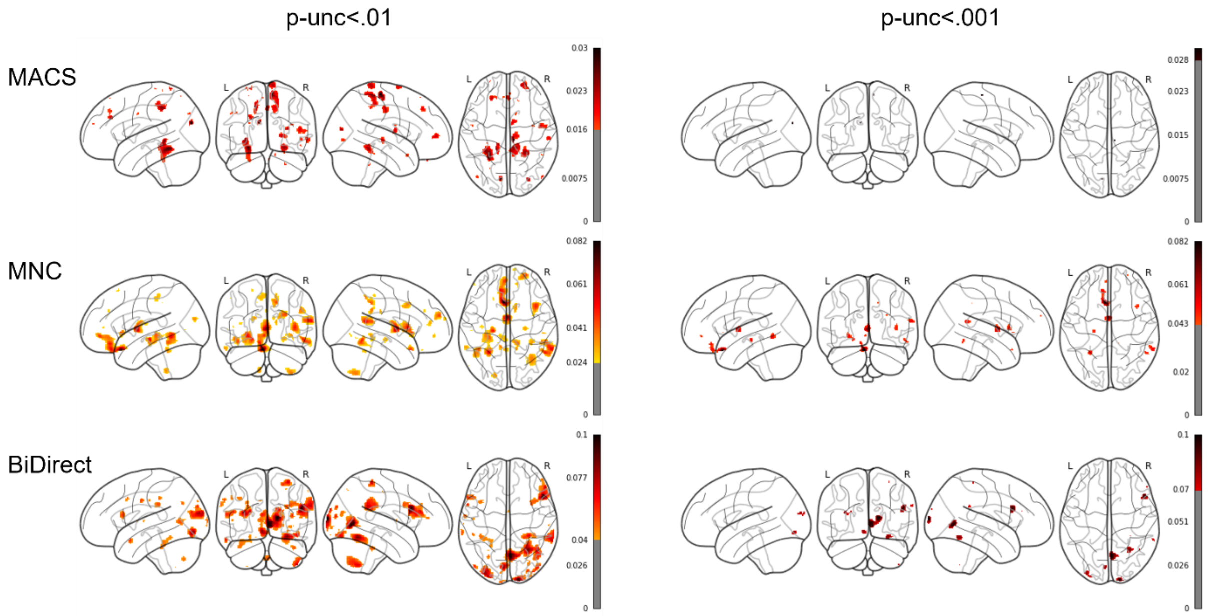


*Note.* Glass brains are shown with maximum intensity projections. Color bars represent the partial $R^2$ of the maltreatment predictor. Results are shown thresholded using the uncorrected thresholds $p_{unc}$<.01 (left column) and $p_{unc}$<.001 (right column).

**Figure S49**

*Significant clusters across cohort-wise analyses (replicability analysis) for* ***model 14*** *– CTQ extreme severity compared to none in HC subsamples –* ***sex-stratified for male subsample***

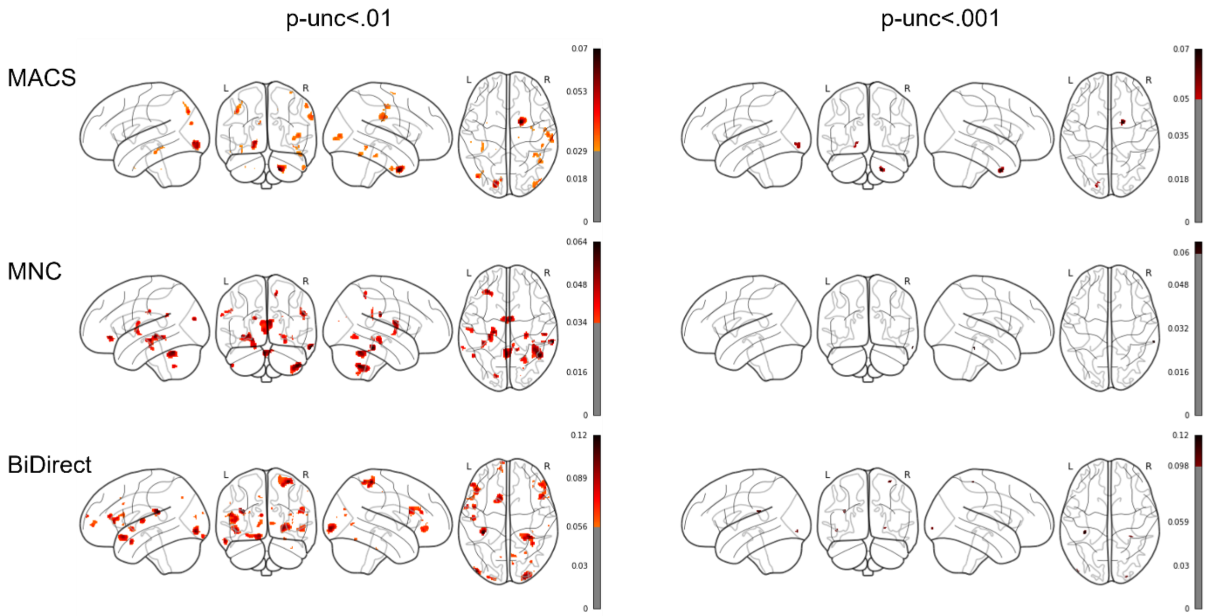


*Note.* Glass brains are shown with maximum intensity projections. Color bars represent the partial $R^2$ of the maltreatment predictor. Results are shown thresholded using the uncorrected thresholds $p_{unc}$<.01 (left column) and $p_{unc}$<.001 (right column).

**Figure S50**

*Significant clusters across cohort-wise analyses (replicability analysis) for* ***model 15*** *– CTQ extreme severity compared to none in MDD subsamples –* ***sex-stratified for male subsample***

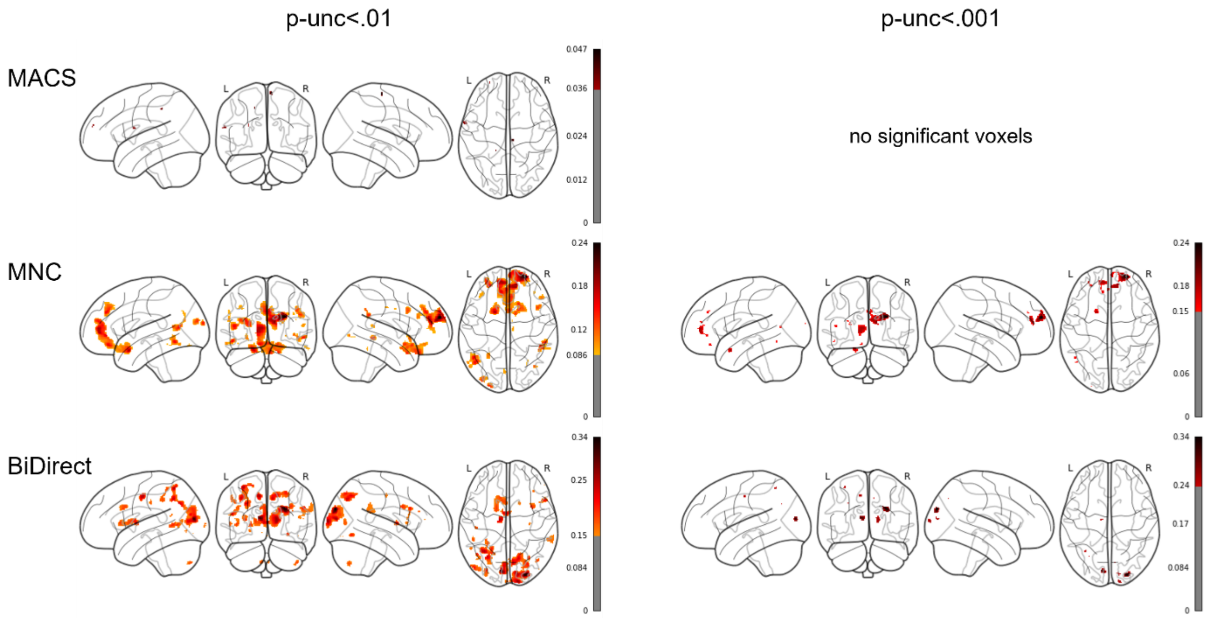


*Note.* Glass brains are shown with maximum intensity projections. Color bars represent the partial $R^2$ of the maltreatment predictor. Results are shown thresholded using the uncorrected thresholds $p_{unc}$<.01 (left column) and $p_{unc}$<.001 (right column).
